# Different orthology inference algorithms generate similar predicted orthogroups among Brassicaceae species

**DOI:** 10.1101/2024.05.21.595184

**Authors:** Irene T. Liao, Karen E. Sears, Lena C. Hileman, Lachezar A. Nikolov

## Abstract

- Premise – Orthology inference is crucial for comparative genomics, and multiple algorithms have been developed to identify putative orthologs for downstream analyses. Despite the abundance of proposed solutions, including publicly available benchmarks, it is difficult to assess which tool to best use for plant species, which commonly have complex genomic histories.
- Methods – We explored the performance of four orthology inference algorithms – OrthoFinder, SonicParanoid, Broccoli, and OrthNet – on eight Brassicaceae genomes in two groups: one group comprising only diploids and another set comprising the diploids, two mesopolyploids, and one recent hexaploid genome.
- Results – Orthogroup compositions reflect the species’ ploidy and genomic histories. Additionally, the diploid set had a higher proportion of identical orthogroups. While the diploid+higher ploidy set had a lower proportion of orthogroups with identical compositions, the average degree of similarity between the orthogroups was not different from the diploid set.
- Discussion – Three algorithms – OrthoFinder, SonicParanoid, and Broccoli – are helpful for initial orthology predictions. Results from OrthNet were generally an outlier but could provide detailed information about gene colinearity. With our Brassicaceae dataset, slight discrepancies were found across the orthology inference algorithms, necessitating additional analyses, such as tree inference to fine-tune results.

## INTRODUCTION

Performing genetic and genomic comparisons across species is central to phylogenetic inference and comparative methods, genome annotations, and functional genomics, which enable the transfer of knowledge from well-studied model systems to less genetically tractable species, such as crops. Thus, identifying the appropriate set of genes for such comparisons is critical. Broadly, genes or loci sharing common ancestry are known as homologs; from a genomic perspective, these genes exhibit sequence similarity. More specifically, genes in different species that originated as a result of a speciation event are defined as orthologs whereas genes that have arisen due to duplications are defined as paralogs (Fitch, 1970). Orthologs are often the target genes for comparative studies, as they represent the “same” gene in different species (Nehrt et al., 2011; Altenhoff et al., 2019; Stamboulian et al., 2020).

The traditional practice to identify orthologs between two species includes reciprocal one-to-one sequence alignment (*e.g.,* BLAST); however, gene duplications and losses, gene conversion events, and whole genome duplications make accurate homology inferences difficult because one-to-one gene correspondence is broken (Wendel, 2015; Altenhoff et al., 2019; Conover et al., 2021). The extent to which these complexities are present and confound homology inference is dependent on the time since species divergence. Additional challenges arise when comparing more than two species at a time. All homologous genes from two or more species descended from a single gene in their most recent common ancestor, whether they are orthologs or paralogs, together form a cluster of orthologous genes, or an orthogroup (Tatusov et al., 1997; Altenhoff et al., 2019; Emms and Kelly, 2019). Thus, compared to the traditional practice to infer one-to-one orthologs through reciprocal searches, an orthogroup approach provides a broader comparable gene space for inferring orthologs for comparative analyses among species, including species with complex gene lineage histories.

Many orthology inference algorithms exist for determining single-copy orthologs among multiple species, without a clear agreement for which algorithm to use for one’s own projects. A consortium of researchers, known as the Quest for Orthologs, was formed to assess best practices and resources for the scientific community (Dessimoz et al., 2012; Nevers et al., 2022). One of the resources is Orthology Benchmark, a repository for method developers to submit the results from their algorithms on a reference set of proteomes (Quest for Orthologs consortium et al., 2016). The results from newly developed algorithms are compared with results from existing algorithms and assessed for the degree of accuracy and sensitivity, which facilitates algorithm choice for other researchers. Additionally, several databases have orthology designations for species across all domains of life, such as OMA (Altenhoff et al., 2021), OrthoDB (Kuznetsov et al., 2023), and eggNOG (Huerta-Cepas et al., 2019); see a full list of databases here: https://questfororthologs.org/orthology_databases. These databases include well-developed model organisms with publicly available genomic resources.

There are several limitations to relying on a database for orthology and homology inference. From the perspective of a plant researcher, many of these repositories and databases lack a broad representation of plant species. According to the Encyclopedia of Life, Viridiplantae (also called Chloroplastida) represent 18.9% of described Eukaryotic species (378543/2003399; Parr et al., 2014). Some larger databases include 8-27% of Viridiplantae species: Orthology Benchmark reference proteome set: 5/34 – 14.7%; OMA: 83/713 – 11.6% (Altenhoff et al., 2021); OrthoDB: 171/1,952 – 8.7% (Kuznetsov et al., 2023), PANTHER: 38/143 – 26.6% (Thomas et al., 2022). There are several plant-specific databases, including Phytozome (Goodstein et al., 2012), GreenPhylDB (Guignon et al., 2021), and PLAZA (Van Bel et al., 2018, 2022), which have incorporated orthology inference as part of their resources. There are 134 Viridiplantae species represented in PLAZA and 46 species represented in GreenPhylDB, with active maintenance and updates to both databases. These resources are useful on a gene-by-gene basis, but more difficult to use on a global genome level, for example, performing a *de novo* orthology inference for a newly annotated genome. Additionally, these databases vary in the frequency of updates – a limitation given that genome annotations, even for well-characterized species, are continuously being improved – and many more genomes are being sequenced and made publicly available on a regular basis. Thus, orthology inference algorithms that allow for species customization is important.

Several commonly used algorithms allow for user-supplied genomic data. OrthoFinder (Emms and Kelly, 2015, 2019) is a phylogenetically informed tree-based inference algorithm where users can select among software packages for sequence alignment and tree inference. SonicParanoid (Cosentino and Iwasaki, 2019) is a graph-based inference algorithm that was modified from the InParanoid algorithm (Sonnhammer and Östlund, 2015), but does not incorporate phylogenetic information in its orthogroup and orthology inference. Both OrthoFinder and SonicParanoid use the Markov Clustering algorithm (MCL, Van Dongen, 2008) to distinguish clusters of similar sequences. Broccoli (Derelle et al., 2020) is a tree-based algorithm and uses network analyses determine orthology networks. All three programs consider gene length biases before clustering proteins based on sequence similarity. Synteny between genes may assist in orthology inferences. CLfinder-OrthNet (Oh and Dassanayake, 2019) is one such workflow that incorporates this information for determining orthogroups, and it also uses MCL to cluster sequences.

The Brassicaceae family, which includes the model species *Arabidopsis thaliana* and several important agricultural crops (e.g. *Brassica* spp., *Sinapis alba*, *Camelina sativa, Thlaspi arvense*) is a model clade for a wide range of comparative studies (Franzke et al., 2011; Nikolov and Tsiantis, 2017; Hendriks et al., 2023; Mabry et al., 2023). *Arabidopsis thaliana* is arguably the most well-studied plant species, with extensive genetic and genomic resources; it often serves as the reference for comparative analyses across plants. Other species in the Brassicaceae have been developed as model systems for studies in evolutionary ecology (*e.g.*, *Boechera stricta*; Rushworth et al., 2011), fruit and leaf morphology (*e.g.*, *Cardamine hirsuta*; Hay and Tsiantis, 2016), and domestication (e.g., *Brassica rapa*; McAlvay et al., 2021), and many of these species have well-annotated genomes. Additionally, a resolved Brassicaceae phylogeny has recently been published (Nikolov et al., 2019; Hendriks et al., 2023). All Brassicaceae species share several whole genome paleopolyploidization events, the most recent along the stem lineage leading to the contemporary diversity in the family after the divergence of its sister family Cleomaceae (Hall et al., 2002; Schranz and Mitchell-Olds, 2006; Nikolov and Tsiantis, 2017). Additionally, lineage, tribe and genus-specific duplication events have created a complex genomic landscape where orthology assessment has been challenging (Couvreur et al., 2010; Hendriks et al., 2023; Mabry et al., 2023; Walden and Schranz, 2023). Given the variation in genome complexity and the ample genomic resources, Brassicaceae species can serve as a model to compare the performance of orthology inference algorithms in species with different ploidies, including mesopolyploid and recent polyploid species.

In this study, we leveraged eight Brassicaceae genomes to infer orthogroups and compare the performance of several orthology inference algorithms. We have opted to use the term “orthogroup inference,” but refer to the algorithms as “orthology inference algorithms” in line with previous literature (Nevers et al., 2022). We focused on two species sets: one set consisting of five diploid species (diploid set), and a second set including the five diploids, two mesopolyploids, and one recent allohexaploid species (diploid+higher ploidy set). We compared the performance of orthology inference algorithms based on the number of species represented in an orthogroup and the distribution of the number of genes from a given species in the orthogroups. We examined the degree of similarity between orthogroup compositions inferred from each algorithm. We found that most of the algorithms infer orthogroups that have similar distributions in the number of species and the number of genes per species regardless of whether the species belonged to the diploid set or to the diploid+higher ploidy set. We found fewer matching orthogroup compositions in the diploid+higher ploidy set, but overall, the orthology inference algorithms yield similar average orthogroup similarity scores across the two species sets.

## METHODS

### Plant genomes

We selected eight Brassicaceae species (Fig. 1, Table 1): the diploid species *Arabidopsis thaliana* (Cheng et al., 2017), *Capsella rubella* (Slotte et al., 2013), *Cardamine hirsuta* (Gan et al., 2016), *Thlaspi arvense* (Nunn et al., 2022), and *Aethionema arabicum* (Fernandez-Pozo et al., 2021), which share the eudicot- and Brassicaceae-specific paleopolyploidization events; the mesopolyploids *Brassica rapa* (Zhang et al., 2018, 2023) and *Sinapis alba* (Yang et al., 2023), which share an additional whole-genome triplication event that defines the Brassiceae tribe (The Brassica rapa Genome Sequencing Project Consortium et al., 2011; Hendriks et al., 2023; Yang et al., 2023; Fig. 1); and the recent hexaploid, *Camelina sativa* (Kagale et al., 2014; Mandáková et al., 2019). Custom scripts were used to extract putative primary transcripts for *Cardamine hirsuta* and *Camelina sativa* (the modified .fasta and .gtf files used as inputs can be found on GitHub and Dryad.

**Fig. 1:**
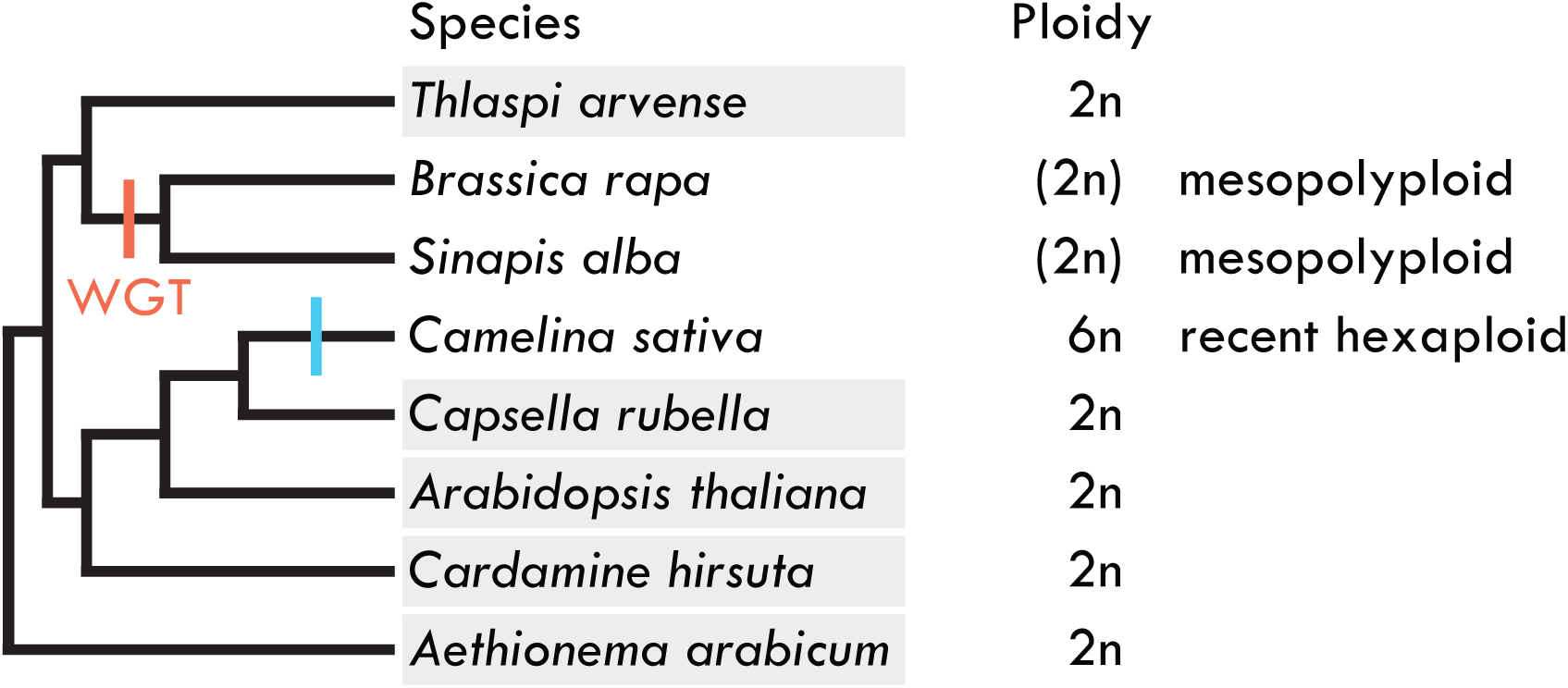
Phylogenetic tree of the species used in the study, including the proposed ploidy of each species (Table 1). Highlighted in light gray are species included in the diploid set; all eight species are included in the diploid+higher ploidy set. Parentheses indicate the mesopolyploid species *Brassica rapa* and *Sinapis alba*, which share a whole genome triplication event (WGT, in red) and has undergone genome fractionation. Blue bar is the *Camelina sativa* specific hexaploidization event.

**Table 1:**
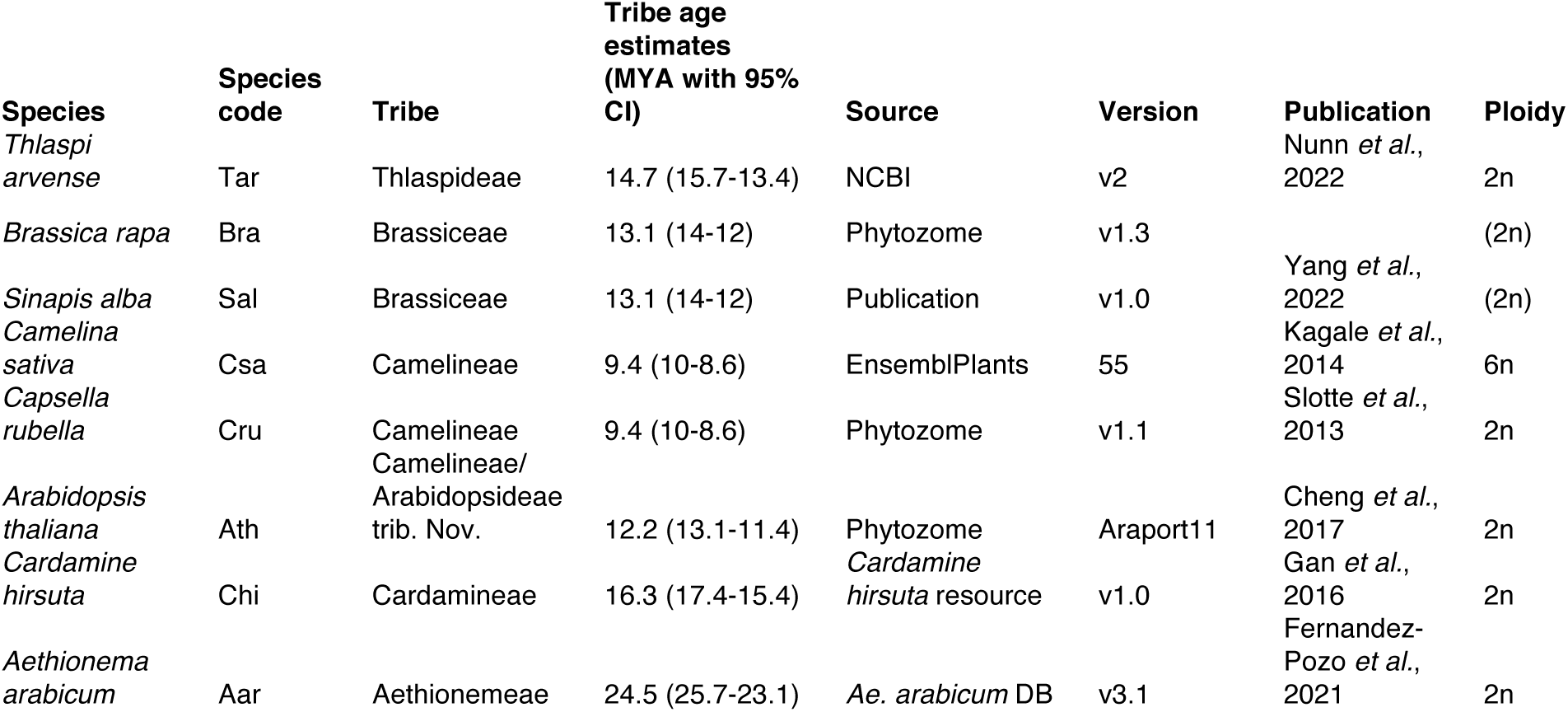
List of species, the genomes used in the study, and the composition of the species sets. Species: species name. Species code: shorthand abbreviation of the species. Tribe age estimates: the mean stem tribe age estimates from nuclear data from Hendriks *et al*., 2023. Source: database/website the genomes was obtained. Version: genome version used. Publication: citation for the genome (if there is a publication associated with it). Ploidy: predicted ploidy. Parentheses indicate that the species is a mesopolyploid.

### Orthology inference algorithms

We tested four software tools: OrthoFinder (Emms and Kelly, 2015, 2019), SonicParanoid (Cosentino and Iwasaki, 2019), Broccoli (Derelle et al., 2020), and CLfinder-OrthNet (Oh and Dassanayake, 2019). The first three were selected based on the overall metrics from the Orthology Benchmark. We included CLfinder-OrthNet, referred to as OrthNet hereafter, to test whether synteny could provide additional information for fine-tuning orthogroup assignments; OrthNet requires GFF annotations as additional input. OrthoFinder is the only algorithm that inferred species-specific orthogroups – these were removed from subsequent analyses.

We tested a total of seven variations of the four orthology algorithms, as summarized in Table 2: Broccoli, OrthoFinder-BLAST, OrthoFinder-DIAMOND, OrthoFinder-MMseqs2, SonicParanoid-DIAMOND, SonicParanoid-MMseqs2, and OrthNet. Because all four algorithms use different default alignment software, we ran OrthoFinder and SonicParanoid with BLAST (Camacho et al., 2009), DIAMOND (Buchfink et al., 2015), and MMseqs2 (Steinegger and Söding, 2017) to examine whether different alignment algorithms contribute to differences in orthology inferences. For OrthoFinder, *Aethionema arabicum* was used as the outgroup species for tree inference. We ran the algorithms with the default settings, except OrthNet, where we changed the MCL inflation parameter from 1.2 to 1.5 to match the default settings of OrthoFinder and SonicParanoid, a change which increases the degree of cluster splitting for OrthNet outputs compared to the default. We refer to each of these seven variations as “algorithms.”

**Table 2:**
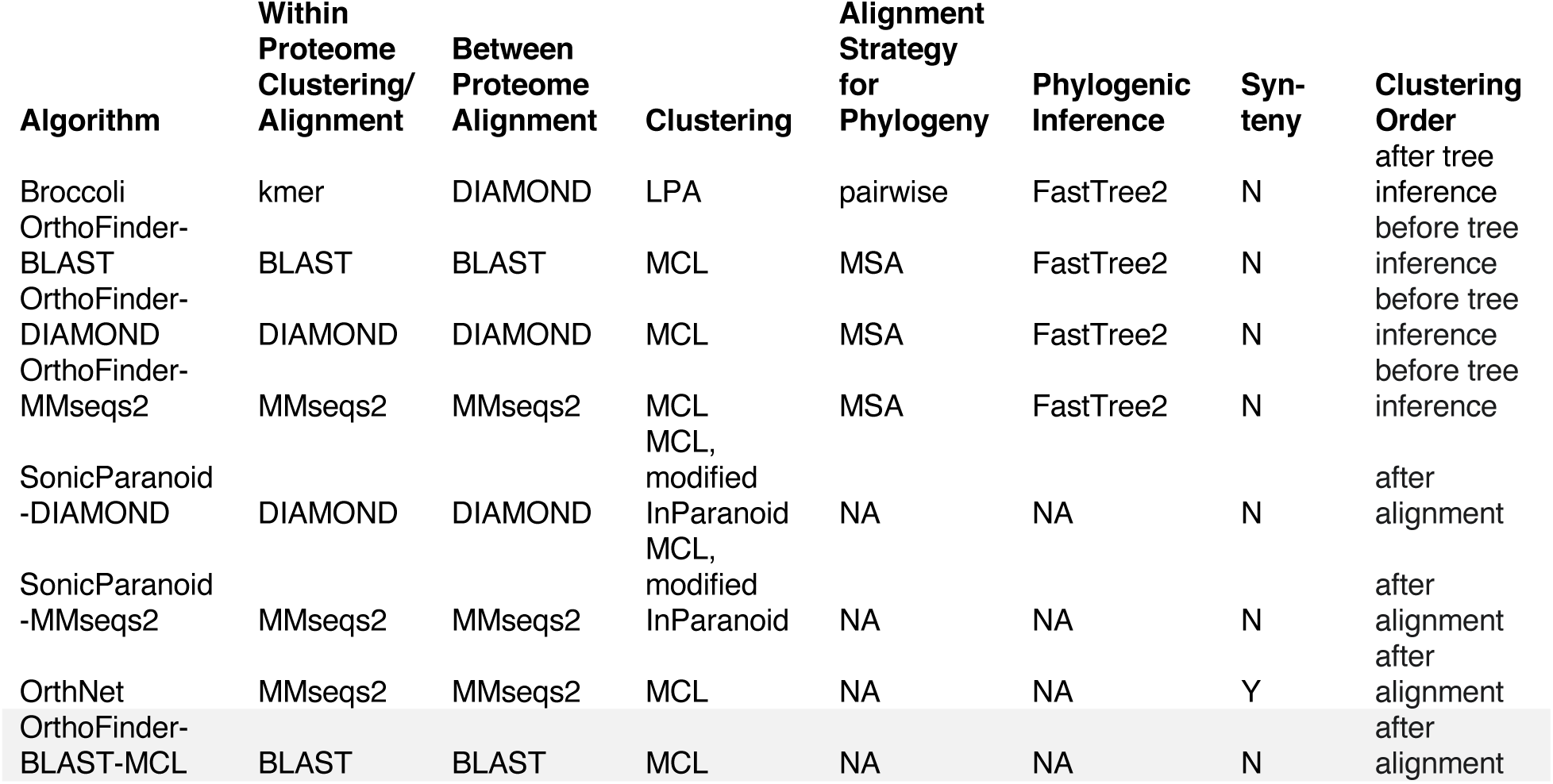
Summary of the orthology inference algorithms tested. Highlighted in gray is the algorithm used as the baseline to compare species pairs orthology inferences with inferences from other algorithms – no phylogenetic inference was run. Algorithm: algorithm name and alignment software used. Within Proteome Clustering/Alignment: alignment method used for clustering proteome sequences within species. Between Proteome Alignment: alignment method used for clustering proteome sequences between species. Clustering: clustering method. Alignment Strategy for Phylogeny: how sequences were aligned before building phylogenetic trees. Phylogenetic Inference: tree inference method. Synteny: whether the method incorporates syntenic information. Clustering Order: step when clustering occurs. LPA: label propagation algorithm. MCL: Markov Clustering Algorithm. MSA: multiple sequence alignment.

The orthogroup sets can be found on GitHub.

### Summary statistics

Each algorithm computes orthogroup sets of genes and provides: 1) the species represented in an orthogroup, and 2) the number of genes per species found in an orthogroup. We used ggplot2 3.4.2 (Wickham, 2016) and ComplexUpset 1.3.3 (Lex et al., 2014; Krassowski, 2020) in R 4.0.2 to process and plot the results. To test whether the distribution of number of species in an orthogroup and the distribution of number of genes per species in an orthogroup differed among algorithms, we performed Kruskal-Wallis rank sum test on all the algorithms and pairwise Wilcoxon rank sum tests between algorithms with multiple hypotheses accounted for with FDR; these tests were chosen because the distributions of the residuals were non-normal.

### Comparing orthogroup composition across algorithms

We then compared orthogroup gene compositions across the seven algorithms for the two species sets (Table 2). To establish correspondence between orthogroups generated with different algorithms, we used the *Arabidopsis* genes as anchors; consequently, we omitted orthogroups without *Arabidopsis* genes. We compared orthogroups on a gene-by-gene basis using the results from two algorithms and presented the results of the pairwise comparisons as the proportion of identical orthogroups and their average similarity scores across all *Arabidopsis* genes.

To assess the degree of similarity among the orthogroups, we calculated three similarity score metrics: Rand Score (RS), Adjusted Rand Score (ARS), and Jaccard similarity Index (JI). RS measures the similarity between two orthogroups, whereas ARS measures the similarity between two orthogroups and corrects for chance gene clustering. Both scores examine the number of gene pairs that are the same between two orthogroups and the number of gene pairs that are different between the two orthogroups. RS and ARS require both orthogroups to contain the same number of genes; for each pair of orthogroups, we determined the union of the genes of the two orthogroups and used the function from scikit-learn v1.0.2 (Pedregosa et al., 2011) to calculate RS and ARS. For example, if orthogroup X generated by one algorithm has three genes (A, B, C) and orthogroup Y generated by another algorithm has four genes (A, B, C, D), to calculate RS and ARS, the union of the four genes (A, B, C, D) is found. Each gene is then coded by whether it is present in both orthogroups (indicated as 0) or not (indicated as 1). In this case, orthogroup X is coded as [0,0,0,1], since “D” was not originally found in this orthogroup, whereas orthogroup Y is coded as [0,0,0,0]. These matrices are compared to calculate RS and ARS.

JI is the ratio of intersection over union:

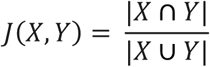

and is calculated in the following manner:

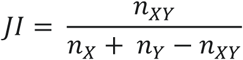

where *n_XY_* is the number of genes that are in common between the orthogroups containing the same *Arabidopsis* gene from algorithm X and from algorithm Y, *n_X_* is the number of genes in the orthogroup from algorithm X, and *n_Y_* is the number of genes in the orthogroup from algorithm Y. JI does not require orthogroups of the same size.

We determined the proportion of identical orthogroups among all the algorithms in a pairwise manner for the diploid set and the diploid+higher ploidy set. We also calculated the average values for each metric to summarize the degree of similarity for all comparisons. We plotted these results as a heatmap in R.

### Examining orthogroups by species pairs

The number of genes in an orthogroup for a pair of species can be categorized as one-to-one (1:1), one-to-many (1:M), many-to-one (M:1), and many-to-many (M:M). To compare whether these distributions differ between algorithms, we estimated orthogroups using OrthoFinder with BLAST alignment and MCL clustering but without tree inference (OrthoFinder-BLAST-MCL) as a baseline; this was done for the 10 species pairs in the diploid set and the 28 species pairs in the diploid+higher ploidy set. We then compared the number of orthogroups in each of these categories between the baseline algorithm and the other algorithms. We also calculated JI to determine the number and proportion of identical orthogroups in a species pair and the average JI value to describe the degree of orthogroup similarity between the baseline algorithm and each of the algorithms tested.

### Case study: orthogroup inference of a small plant-specific gene family

We studied the YABBY transcription factor family, a small plant-specific gene family which consists of six paralogs in *Arabidopsis thaliana*: AT1G69180.1 (CRC), AT2G45190.1 (FIL/YAB1), AT1G08465.1 (YAB2), AT4G00180.1 (YAB3), AT2G26580.1 (YAB5), and AT1G23420.2 (INO).

For the diploid+higher ploidy set, we extracted all genes found in the same orthogroup as the *Arabidopsis* YABBY family genes as long as the gene was found by at least one of the six synteny-agnostic algorithms. To build gene trees for each orthogroup, we used the *Aethionema* sequence as the outgroup, except for YAB3, where no *Aethionema* YAB3 homolog was found. We used MAFFT (https://www.ebi.ac.uk/Tools/msa/mafft/) without manual modification to align the sequences and visualized the alignments in Aliview v1.28 (Larsson, 2014). We used RAxML v8 2.12 (Stamatakis, 2014) using the default settings with 1000 bootstraps via the CIPRES portal (Miller et al., 2010), visualized the trees in FigTree v1.4.4 (http://tree.bio.ed.ac.uk/software/figtree/), and mapped the presence/absence of each gene in the results from each algorithm. To examine reciprocal colinearity from OrthNet, we visualized the clusters from the diploid set and diploid+higher ploidy set using Cytoscape v3.9.1 (Shannon et al., 2003).

## RESULTS

### The majority of orthogroups include all examined species across all algorithms

We first examined the number of orthogroups generated by the seven algorithms. For the diploid set, the number of orthogroups ranged from 19596 to 22191, while for the diploid+higher ploidy set, the number ranged from 20492 to 24875 (Appendices S1-2; see Supporting Information with this article). For both sets, OrthNet yielded the smallest number of orthogroups (diploid set: 19596, diploid+higher ploidy set: 20492).

We then examined the species composition of each orthogroup derived under the seven algorithms. For the diploid set, 60-74.1% of the orthogroups contain all five species (*Arabidopsis, Capsella, Cardamine*, *Thlaspi*, and *Aethionema*) and 50.7-69.5% of the orthogroups contain all eight species for the diploid+higher ploidy set (Figure 2, Appendices S1, S3). For the diploid set, 62.3-83.8% of orthogroups from non-OrthNet algorithms are single-copy orthogroups, yet only 49.6% of such orthogroups are single copy for OrthNet (Appendix S3). Additionally, OrthNet resulted in a markedly higher number of orthogroups containing all species from both the diploid set (14524, 74.1%) and the diploid+higher ploidy set (14251, 69.5%). Under the examined parameter, OrthNet produced a higher mean number of species per orthogroup and fewer orthogroups overall compared to all the other algorithms (Appendices S1-4).

**Fig. 2:**
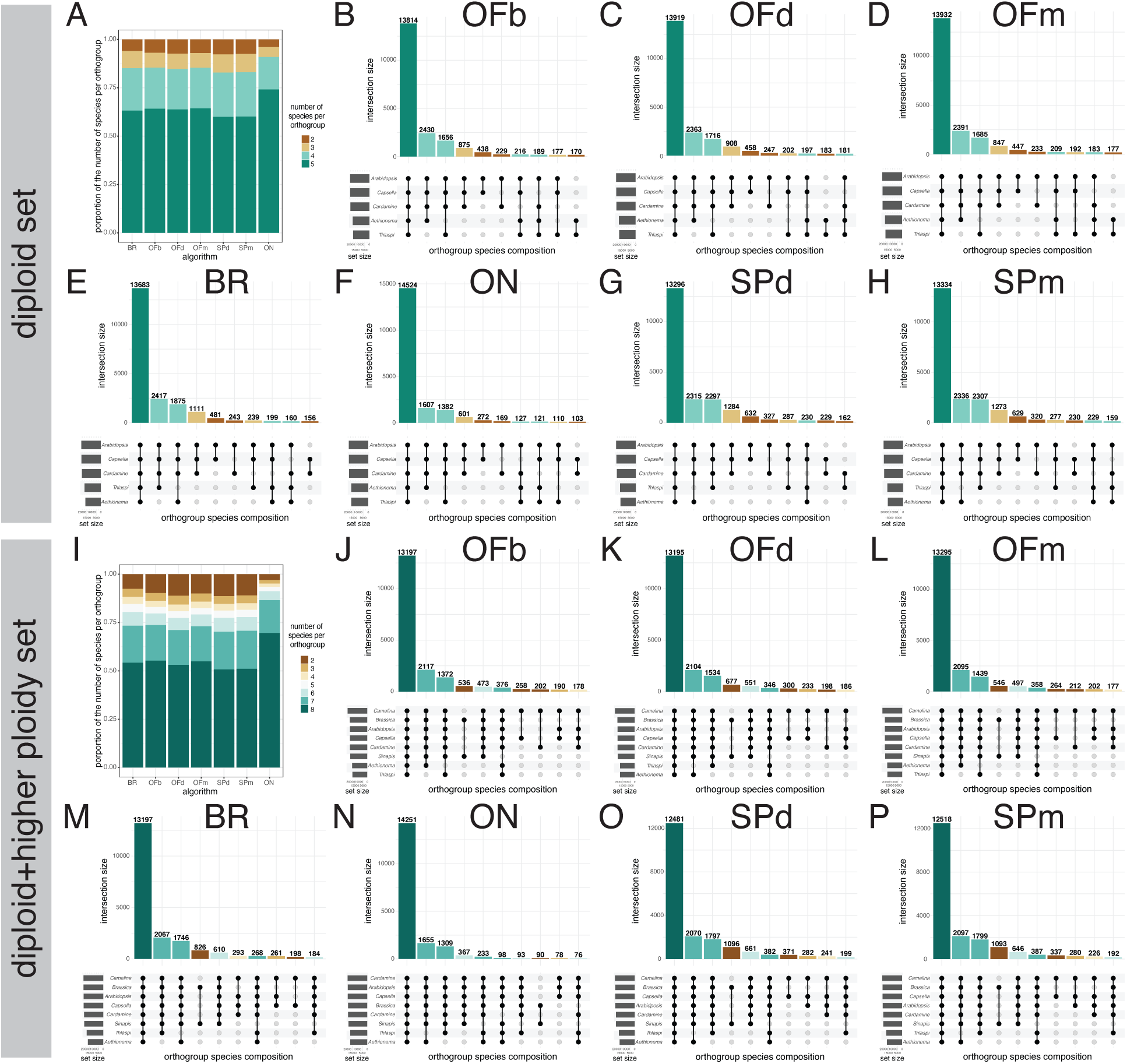
Distribution of the number of species found in an orthogroup is similar across most algorithms. (A, I) Stacked bar plots of the (A) diploid set and (I) diploid+higher ploidy set (I). Each bar represents the results from the seven algorithms tested, and each color represents the specific number of species found in an orthogroup. (B-H, J-P) Upset plots reveal the specific distribution of the orthogroup species compositions. Horizontal bar plots on the left indicate the number of times the species is found in an orthogroup. Horizontal lines on the bottom represents each species, and circles indicate the presence (filled) or absence (empty) of the species in an orthogroup. Vertical bars with numbers indicated the number of orthogroups that have the specific species composition. The 10 most abundant species compositions are displayed. (B-H) Diploid species. (J-P) Diploid+higher ploidy set. Corresponds to Appendix S1. BR: Broccoli. OFb: OrthoFinder-BLAST. OFd: OrthoFinder-DIAMOND. OFm: OrthoFinder-MMseqs2. SPd: SonicParanoid-DIAMOND. SPm: SonicParanoid-MMseqs2. ON: OrthNet.

For both diploids and diploids+higher ploidy sets, most orthogroups included all species, followed by orthogroups including four of five species (diploid set) or seven of eight species (diploid+higher ploidy set; Fig. 2). More orthogroups included *Aethionema arabicum* and missing *Thlaspi arvense* genes across all orthology inference algorithms, except Broccoli. We also identified Brassiceae-(*Sinapis alba* and *Brassica rapa*), Camelinae-(*Capsella rubella* and *Camelina sativa*) and Lineage I-specific orthogroups (*Arabidopsis thaliana*, *Capsella rubella*, *Cardamine hirsuta*, and/or *Camelina sativa*).

Different algorithms produced orthogroups with different distributions of the number of species per orthogroup (Fig. 2, Table 3, Appendix S5). Species number per orthogroup is significantly different across algorithms for the diploid set (Kruskal-Wallis test, χ^2^ = 1250.9; *P* < 2.2 E -16) and the diploid+higher ploidy set (Kruskal-Wallis test, χ^2^ = 2648.3; *P* < 2.2 E -16). These significance values hold even when excluding OrthNet results from the analyses (Appendix S5); in pairwise comparisons, results from OrthNet had significantly different distributions from all other algorithms (Table 3). Comparing the distributions in a pairwise manner using the Wilcoxon rank sum test, for the diploid set, all results from SonicParanoid algorithms (SonicParanoid–DIAMOND, SonicParanoid–MMseqs2) were statistically different from all the OrthoFinder algorithms (OrthoFinder–BLAST, OrthoFinder–DIAMOND, OrthoFinder–MMseqs2) and Broccoli (*P* < 0.05). The same pattern was found for the diploid+higher ploidy set with additional significant differences between OrthoFinder–DIAMOND and Broccoli, OrthoFinder– DIAMOND and OrthoFinder–BLAST, and OrthoFinder–DIAMOND and OrthoFinder–MMseqs2.

**Table 3:**
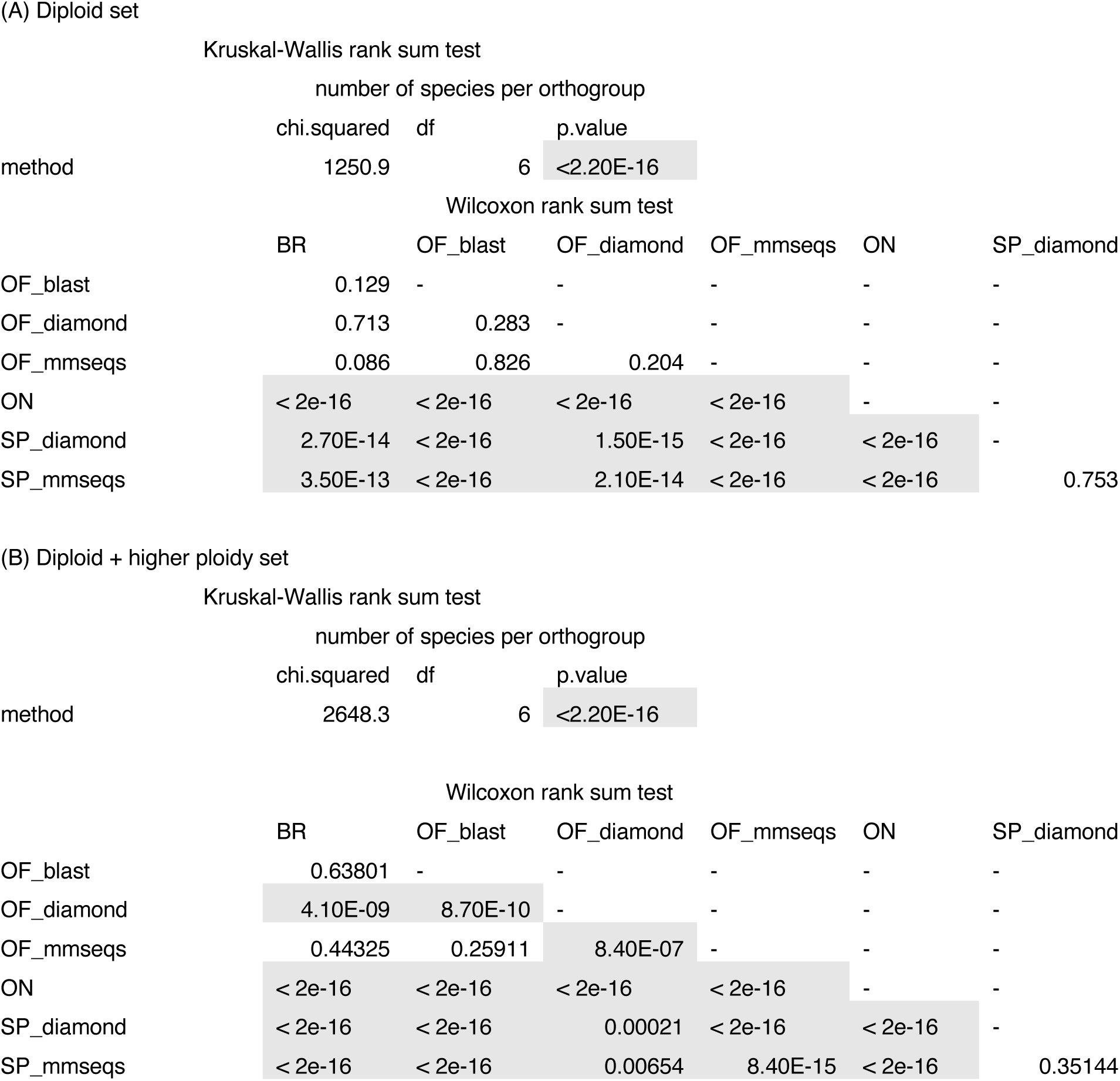
Comparison of the number of species per orthogroup detected across orthology interference methods. Asterisks or gray highlight indicate significance at P < 0.05 after a FDR adjustment for multiple comparisons. (A) Diploid set. (B) Diploid+higher ploidy set. Top table: results from a Kruskal-Wallis rank sum test. Bottom table: p-values from all possible pairwise comparisons of the algorithms tested through a Wilcoxon rank sum test after a FDR correction. chi.squared is the test statistic used to calculate the p-value. BR: Broccoli. OF_blast: OrthoFinder + BLAST. OF_diamond: OrthoFinder-DIAMOND. OF_mmseqs: OrthoFinder-MMseqs2. SP_diamond: SonicParanoid-DIAMOND. SP_mmseqs: SonicParanoid-MMseqs2. ON: OrthNet. Corresponds to Fig. 2.

### All algorithms recover the ploidy of the species based on the number of genes in an orthogroup

The number of genes per orthogroup for each species shows evidence of their shared and lineage-specific whole genome multiplication(s) (Table 1). For diploid species, the majority of the orthogroups are expected to include a single gene per species (i.e., 1:1:1:1:1 orthologs, the most common category used in comparative analyses), whereas for mesopolyploids and recent polyploids, the majority of the orthogroups are expected to include additional genes from each species (*e.g.,* tetraploid – two genes; hexaploid – three genes). This pattern is consistent with our observations for all algorithms (Figure 3, Appendices S6-7). The majority of the orthogroups contained a single gene for the diploid species *Arabidopsis thaliana*, *Capsella rubella*, *Cardamine hirsuta*, *Thlaspi arvense*, and *Aethionema arabicum* in analyses based on the diploid and the diploid+higher ploidy set (Fig. 3A). In the mesopolyploids *Brassica rapa* and *Sinapis alba,* fewer orthogroups contain only one gene and more orthogroups contain two genes compared to diploids (Fig. 3B). Finally, the majority of orthogroups contain three genes for the recent allohexaploid *Camelia sativa* (Fig. 3C).

**Fig. 3:**
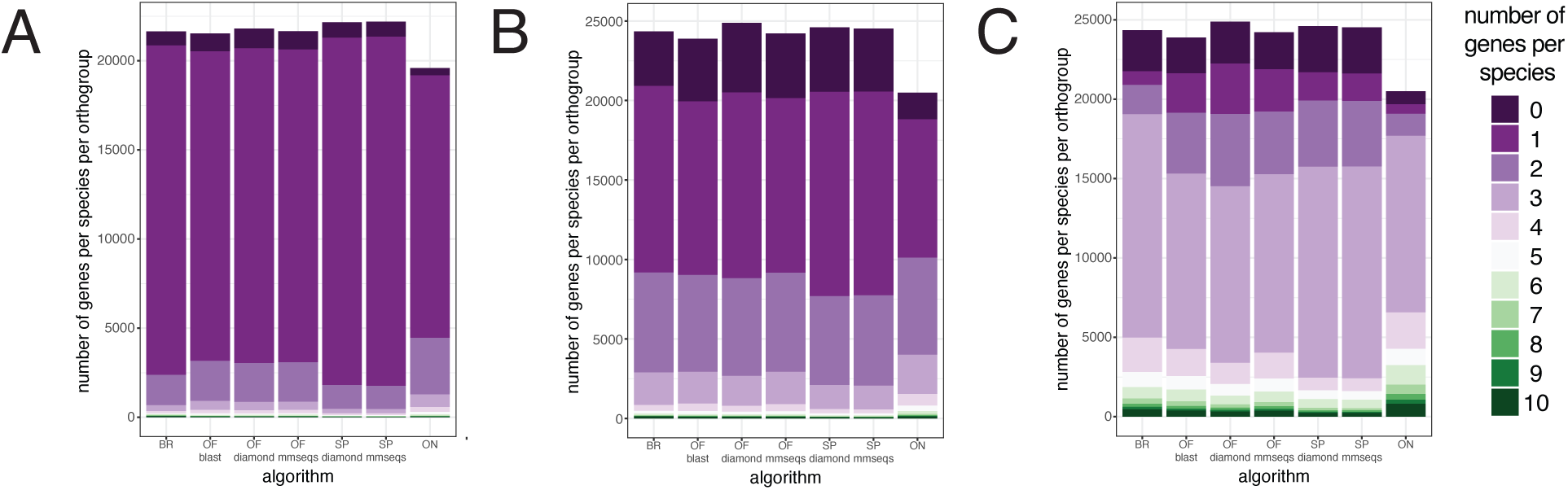
Distributions of the number of genes per species found in an orthogroup reflect the predicted ploidy of the species. Stacked bar plots displaying representative (A) diploid species (*Arabidopsis thaliana*), (B) mesopolyploid species (*Sinapis alba*), and (C) hexaploid species (*Camelina sativa*). Diploid species used in both diploid and diploid+higher ploidy sets show the same patterns. Each plot displays the results for one species across the different algorithms, and each color represents the specific number of genes per species found in an orthogroup. Corresponds to Appendix S6. BR: Broccoli. OF_blast: OrthoFinder-BLAST. OF_diamond: OrthoFinder-DIAMOND. OF_mmseqs: OrthoFinder-MMseqs2. SP_diamond: SonicParanoid-DIAMOND. SP_mmseqs: SonicParanoid-MMseqs2. ON: OrthNet.

The distributions of the number of genes in an orthogroup for each species varied across algorithms (Appendix S8). For example, the distributions of the number of *Arabidopsis* genes in an orthogroup was different among all tested algorithms (Kruskal-Wallis rank sum test, chi-squared = 2400.4, *df* = 6, *P* < 2.20 E-16). Pairwise comparisons for *Arabidopsis* show most comparisons between algorithms are significantly different, except the results comparing SonicParanoid-DIAMOND and SonicParanoid-MMseqs2 *(P* = 0.743), OrthoFinder-BLAST and OrthoFinder-MMseqs2 (*P* = 0.228), and OrthoFinder-DIAMOND and OrthoFinder-MMseqs2 (*P* = 0.126). For all comparisons within each species, the results from SonicParanoid-DIAMOND and SonicParanoid-MMseqs2 were consistently not significantly different from each another (*P* > 0.4).

### Orthogroup composition is variable across algorithms, but more so for the diploid+higher ploidy set than the diploid set

No two algorithms produced identical orthogroup gene compositions for every orthogroup inferred based on the RS, ARS, and JI metrics (Figure 4, Appendix S9-10). The highest degree of similarity was found between algorithms that used the same suite of software but different alignment tool. The proportion of orthogroups with identical gene compositions was highest for SonicParanoid-DIAMOND and SonicParanoid-MMseqs2 (diploid: 0.935, diploid+higher ploidy: 0.858) and among OrthoFinder-BLAST, OrthoFinder-DIAMOND, and OrthoFinder-MMseqs2 (diploid set average: 0.856, diploid+higher ploidy set average: 0.627) compared to any other pairwise comparisons (Fig. 4, upper left triangles). For all other pairwise comparisons, the proportions of identical orthogroups between algorithms range from 0.511–0.660 for the diploid set and 0.288–0.437 for the diploid+higher ploidy set (Fig. 4, Appendix S10). Overall, the proportion of orthogroups with identical composition is higher for the diploid set. On the other hand, the average orthogroup similarity scores are similar between the diploid set and diploid+higher ploidy set, with average JI values in the 0.7-0.9 range, which are higher in the diploid set compared to the diploid+higher ploidy set (Appendix S11).

**Fig. 4:**
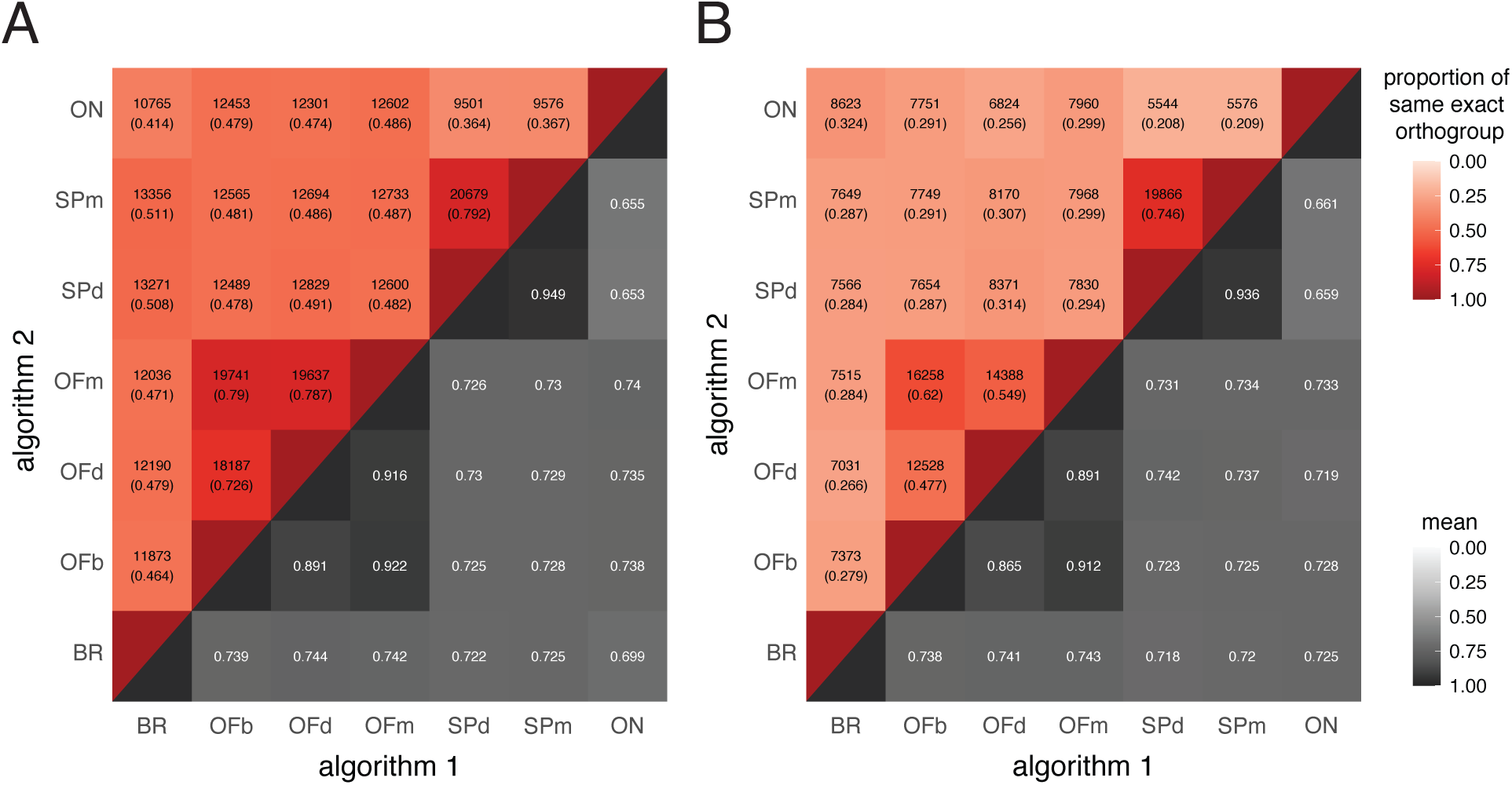
Orthogroup gene compositions are more similar across algorithms tested for diploid species than for those from diploid + higher ploidy species. The Jaccard Index (JI) was calculated for all algorithms in a pairwise manner. The upper left triangle represents the number of orthogroups with identical gene composition (JI = 1) with the numbers in parentheses and the red color gradient representing the proportion of orthogroups with the same composition. The lower right triangle represents the mean JI value; the gray gradient represents the mean values. (A) Diploid set. (B) Diploid+higher ploidy set. Corresponds to Appendix S13. BR: Broccoli. OFb: OrthoFinder-BLAST. OFd: OrthoFinder-DIAMOND. OFm: OrthoFinder-MMseqs2. SPd: SonicParanoid-DIAMOND. SPm: SonicParanoid-MMseqs2. ON: OrthNet.

### Species pairs gene copy ratios reveal general patterns of additional subclustering in orthology inference algorithms

We compared all orthogroup algorithms and a baseline algorithm – OrthoFinder-BLAST-MCL without tree inference – for pairs of species to quantify the similarity between orthogroup compositions produced with different algorithms. The orthogroup algorithm results were partitioned into two-species orthogroups, 10 species pairs in the diploid set and 28 species pairs in the diploid+higher ploidy set.

Our expectation for diploid species is that the majority of genes are single copy (1:1). If one species is a diploid and the other species has a different ploidy (mesopolyploid or hexaploid), we expect most of the genes to consist of one-to-many genes. Finally, if both species are non-diploids, we expect the majority relationship to be many-to-many genes. For all species pairs in the diploid set, the majority of orthogroups indeed consisted of a single gene copy from each species (1:1), regardless of the orthology inference algorithm (Fig. 5A, Appendices S12-13). Orthology inferences between *Arabidopsis thaliana* and *Capsella rubella* using OrthoFinder-BLAST yielded the highest proportion of identical orthogroups (0.634) and average similarity (JI = 0.708) with the baseline algorithm for the diploid set (Appendices S14-15). In general, all the algorithms (besides OrthNet) generated more 1:1 single-copy orthogroups than the baseline algorithm (OrthoFinder-BLAST-MCL).

**Fig. 5:**
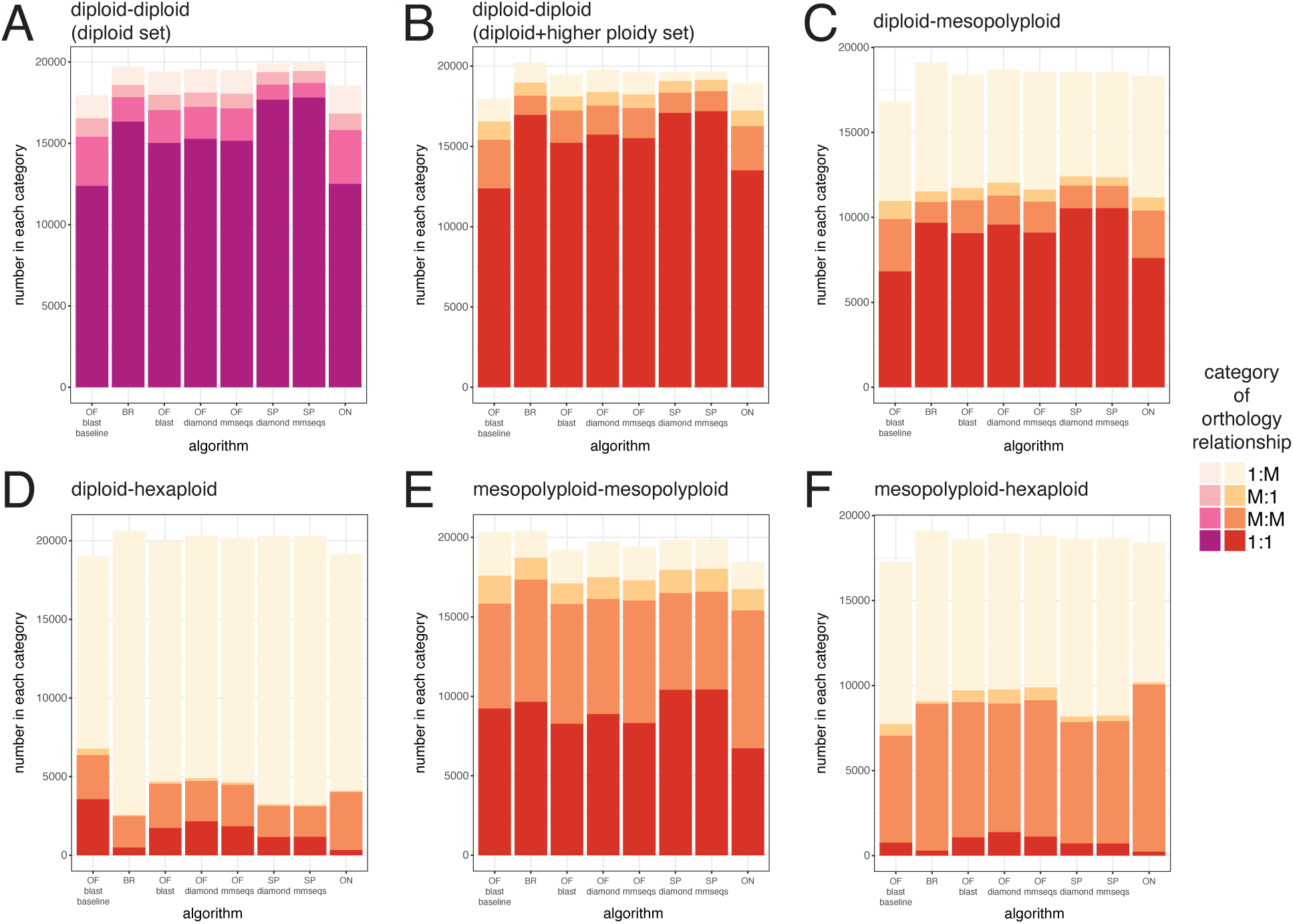
Proportion of predicted orthology relationships between all species pairs across algorithms for the diploid set and the diploid+higher ploidy set. Stacked bar plots displaying representative (A-B) diploid-diploid species pair (*Arabidopsis thaliana* and *Cardamine hirsuta*) from the (A) diploid set and (B) the diploid+higher species set; (C) diploid-mesopolyploid species pair (*Arabidopsis thaliana* and *Sinapis alba*); (D) diploid-hexaploid species pair (*Arabidopsis thaliana* and *Camelina sativa*); (E) mesopolyploid-mesopolyploid species pair (*Sinapis alba* and *Brassica rapa*); (F) mesopolyploid-hexaploid species pair (*Sinapis alba* and *Camelina sativa*). Each stacked bar represents the algorithm used, and the colors represent the category of orthology relationships: 1:1 (one-to-one), 1:M (one-to-many), M:1 (many-to-one), M:M (many-to-many). OFb_base: OrthoFinder-BLAST-MCL, the baseline to compare the results from all other algorithms. BR: Broccoli. OFb: OrthoFinder-BLAST. OFd: OrthoFinder-DIAMOND. OFm: OrthoFinder-MMseqs2. SPd: SonicParanoid-DIAMOND. SPm: SonicParanoid-MMseqs2. ON: OrthNet. Corresponds to Appendices S12, S13.

Species pairs in the diploid+higher ploidy set are more complex given the evolutionary history of the mesopolyploid and the recent hexaploid genomes (Fig. 5B-E, Appendices S12B-E, S13B, S14B, S15B). Including these species did not affect the diploid species pairs (*Arabidopsis*, *Cardamine*, *Capsella*, *Thlaspi*, *Aethionema*) where levels of 1:1 orthogroups were consistent with the expectations for diploids. Similar to the diploid set, for the diploid+higher ploidy set, the highest proportion of identical orthogroups was found for the *Arabidopsis-Capsella* species pair (OrthoFinder-MMseqs, 0.625). The highest average similarity was found between the baseline and either OrthoFinder-BLAST or OrthoFinder-MMseqs (both approximately JI = 0.697). Species pairs that included *Brassica* or *Sinapis* generally had a smaller proportion of orthogroups consisting of 1:1 orthologs and a greater proportion of one-to-many or many-to-one orthogroups, relative to the comparison between two diploid species. Finally, for species pairs that included *Camelina* and a diploid species, most orthogroups contained many-to-one genes. Generally, the lowest similarity values resulted from inferences that included *Camelina sativa* as one of the species in the species pair.

### Case study: YABBY sequence features affect the inclusion of the sequence in orthogroups for orthology inference algorithms

Each algorithm identified six orthogroups corresponding to the six Arabidopsis YABBY paralogs (Fig. 6, Appendix S16), but the orthogroup compositions varied in at least one of the inferences. For example, all the algorithms except Broccoli produced the same gene composition for the INO orthogroup, the gene tree is identical to the species tree, and the genes have a high degree of reciprocal colinearity for both the diploid set and diploid+higher ploidy set (Fig. 6A). While the same high degree of reciprocal colinearity is observed for the CRC orthogroup, the gene tree does not match the species tree precisely (Fig. 6B). Additionally, several algorithms did not include one of the three *Camelina* paralogs (Csa07g035840.1, Fig. 6B) in their results, whereas OrthNet included an extra *Sinapis* gene (Sal09g27760L) in the CRC orthogroup, which is not colinear with other genes in the orthogroup.

**Fig. 6:**
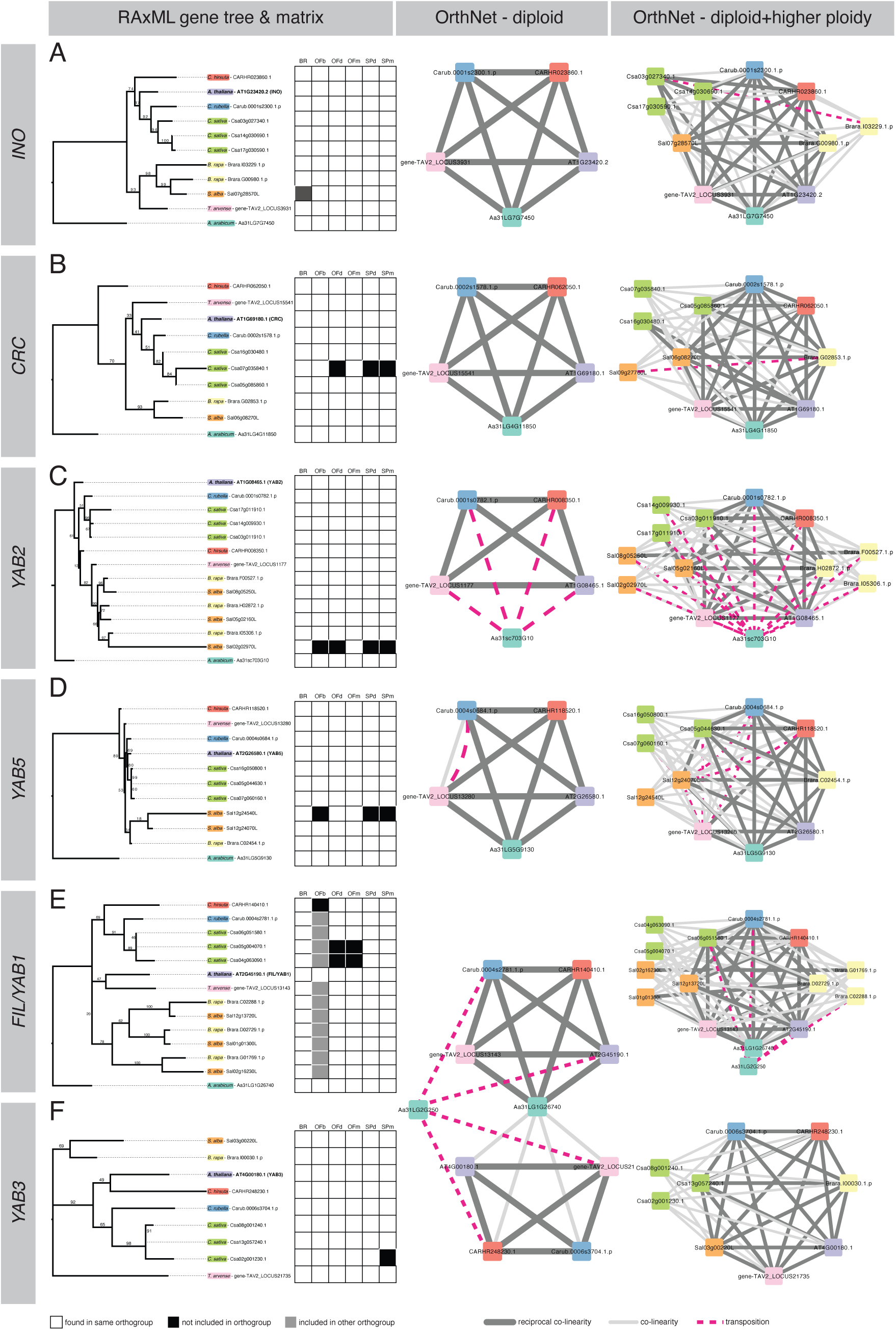
Orthogroup compositions of YABBY genes vary for the diploid+higher ploidy set. (A) Gene trees for individual YABBY orthogroups reflect the most inclusive gene composition from all algorithms except OrthNet. Matrix next to the genes indicates whether the gene is found in the same orthogroup. Each row represents a gene, and each column represents the algorithm tested. Colors represent whether the gene was found in the same orthogroup (white), a different orthogroup (light gray), or not found in any orthogroup (black) resulting from each algorithm. (B-C) Clusters from OrthNet for the (B) diploid set and (C) diploid+higher ploidy set with lines indicating reciprocal co-linearity (solid dark gray), co-linearity (solid light gray), and a transposition in one or more of the genes compared (dashed dark pink). BR: Broccoli. OFb: OrthoFinder-BLAST. OFd: OrthoFinder-DIAMOND. OFm: OrthoFinder-MMseqs2. SPd: SonicParanoid-DIAMOND. SPm: SonicParanoid-MMseqs2.

The YAB2 and YAB5 orthogroups also showed variation in the gene composition. In these cases, *Sinapis* proteins Sal02g02970L in YAB2 and Sal12g24540L in YAB5 were missing from several orthology inference results (Fig. 6C, D). Examining the protein alignments revealed that while conserved regions of the protein align well, the specific gene annotations lack or include additional amino acids (*e.g., Sinapis* proteins Sal02g02970L in YAB2; Appendix S17B). These sequence variations did not appear to affect the OrthNet inference, and additional colinearity information for these paralogs was observed as well.

In the FIL/YAB1 orthogroup, only the *Arabidopsis* and *Aethionema* genes were found consistently in the same orthogroup. The OrthoFinder-BLAST inference resulted in orthogroup splitting after the initial MCL clustering. Additionally, for both diploid and the diploid+higher ploidy set for FIL/YAB1 (AT2G45190.1), all algorithms included Aa31LG1G26740; for the diploid set, OrthoFinder-BLAST included an extra *Aethionema* gene, Aa31LG2G250 (Appendix S18). Conversely, both the diploid and diploid+higher ploidy sets do not include an *Aethionema* gene in their orthogroups for YAB3 (AT4G00180.1). This pattern is also reflected in the OrthNet clusters, where the *Aethionema* sequence Aa31LG1G26740 was shared between the two groups, preventing additional subclustering. However, in the diploid+higher ploidy set, OrthNet included Aa31LG1G26740 in the FIL/YAB1 orthogroup, given that it is reciprocally colinear with all the other FIL/YAB1 homologs. The other *Aethionema* copy Aa31LG2G250 is retained in the diploid+higher ploidy set for FIL/YAB1, but is considered positionally in a non-syntenic region, relative to both FIL/YAB1 and YAB3 homologs.

## DISCUSSION

### Similarities and differences among orthology inference algorithms

Here, we compared several customizable orthology inference algorithms, OrthoFinder, SonicParanoid, and Broccoli, and one pipeline that incorporates synteny (OrthNet), to examine their performance on two sets of plant species with complex genomic histories. Without a “ground-truth” to compare our results, we first examined the overall summary statistics from the orthology algorithms. At a broad scale, we found that inferences by all algorithms produced mostly orthogroups that contained genes from all species (five for the diploid set; eight for the diploid+higher ploidy set), and the number of gene copies in an orthogroup matched the predicted ploidy of the species. The number of orthogroups found was also similar across all the algorithms, except for OrthNet, which recovered less orthogroups, likely due to decreased granularity under the examined parameter regime.

Oh and Dassanayake (2019) developed CLfinder-OrthNet, which incorporates syntenic information to identify colinear, duplicated, or transposed orthologs, to identify duplication and gene transposition events across six Brassicaceae diploid species and to infer patterns of adaptation and speciation in extremophytes. They compared their results to OrthoFinder and found that 70.1% of OrthNet orthogroups had the same composition as orthogroups from OrthoFinder. However, in our study, the percentage of identical orthogroups between OrthNet and the three OrthoFinder results were much lower (diploid set: 47.4-48.6%; diploid+higher ploidy set: 25.6-29.9%; Fig. 4), even when adjusting the default MCL inflation parameter of 1.2 to 1.5. The difference in percentages could be due to the species sets used in the analyses, the divergence times of the species included in both studies, and our inclusion of *Aethionema* as the outgroup.

While the proportions of identical orthogroup compositions were higher for the diploid set, the average orthogroup similarity was nearly the same between the diploid set and the diploid+higher ploidy set. Species pair orthology inferences compared to our baseline algorithm (OrthoFinder-BLAST-MCL) were higher when both species are diploids, indicating that the traditional approach for inferring orthologs, such as reciprocal BLAST search, often identifies the same orthologs and is more efficient for diploid species. Phylogenetic distance also plays a role; for instance, the highest proportion of identical orthogroups and highest degree of similarity between the baseline approach and all other orthology inference algorithms was found in *Arabidopsis* and *Capsella* (Appendices S14-15), which diverged approximately 9.4 mya (Hendriks et al., 2023). When higher ploidy species are included, comparing the results between the baseline and other approaches yielded lower similarity scores, possibly due to fewer single-copy orthogroups identified in the baseline approach for diploid-mesopolyploid (Fig. 5C, Appendix S12C) and diploid-hexaploid species pairs (Fig. 5D, Appendix S12D). These results imply that resolving the relationships among complex genomes requires comparisons with more genomes rather than a one-to-one comparison. Finally, some of the differences between the baseline and the other algorithms may result from the lack of additional sub-clustering in the baseline as the number of orthogroups detected with the baseline algorithm were generally lower compared to all other algorithms (except OrthNet in some cases; Appendices S12-13).

The lack of complete congruence between inference algorithms has been shown in other studies across a broad spectrum of algorithms using more distantly related species (Deutekom et al., 2021; Nevers et al., 2022). Similar to the findings of Cosentino and Iwasaki (2023), we generally find that changing parameters and alignment software for an algorithm introduces variation in the orthogroup inference. We also find that the average degree of similarity between the orthogroups across all algorithms is high regardless of whether the set of compared species contain just diploids or species with higher ploidy. Further examination showed that a comparison with only diploid species results in a higher number of orthogroups with identical composition versus a comparison including species of different ploidy. This pattern is not surprising given the preponderance of paralogs from more species with more complex genomes, including mesopolyploid species, which make it difficult to determine whether certain gene copies should be included in an orthogroup. Because of potential discrepancies in orthogroup inference, attempts to generate orthology inferences with a broader consensus by aggregating the results from multiple algorithms using meta-methods have been implemented in repositories for genetic information from model organisms, such as HGNC Comparison of Orthology Predictions (Yates et al., 2021) and DIOPT (Hu et al., 2011). These derived inferences lead to higher precision but lower recall (Altenhoff et al., 2019) and limit the shared gene space for comparative studies.

Orthology inference algorithms are constantly being updated and improved upon, oftentimes for scalability and speed. For instance, SonicParanoid2 incorporates machine-learning in its inference pipeline, which increases the speed and yields a similar accuracy result (Cosentino and Iwasaki, 2023). Other algorithms incorporate synteny to visualize patterns of orthology and genomic positional information, such as GENESPACE (Lovell et al., 2022) and pSONIC (Conover et al., 2021), both of which build upon OrthoFinder results. Synteny can assist with distinguishing paralogs and identifying syntenic orthologs to use for species tree reconstruction; however, for a set of Brassicaceae species, incorporating the additional syntenic information did not lead to different species trees compared to previous Brassicaceae phylogenies (Huang et al., 2016; Nikolov et al., 2019; Hendriks et al., 2023; Walden and Schranz, 2023). Finally, a new tool TOGA (Tool to infer Orthologs from Genome Alignments), combines orthology inference and gene annotation with machine learning, reporting more accurate orthologous loci throughout the genome (Kirilenko et al., 2023). Although TOGA has only been used in mammals and birds, it will be interesting to see whether TOGA may also improve orthology inference in plants.

### Brassicaceae genomic history is reflected in orthogroup analyses

For both the diploid and diploid+higher ploidy set, *Aethionema arabicum* is sister to the rest of Brassicaceae and can serve as an outgroup for the clade composed of the rest of the species (Fig 2). However, the species composition of the second highest number of orthogroups included *Aethionema arabicum* and excluded *Thlaspi arvense*. The more divergent position of *Aethionema* may resulted in less sequence similarity to the other species but the exclusion of *Thlaspi* is surprising, with both biological (*e.g.* unusual rate of molecular evolution) and technical (*e.g.*, quality of the genomic resources) factors shaping the result. Walden and Schranz (2023) identified syntenic orthologs and paralogs across 11 diploid Brassicaceae species and found 6058 syntenic orthologs, of which 3833 were single-copy across all species; they also used OrthoFinder to identify 3463 orthogroups comprising single-copy orthologs. They also found 95.9% of the syntenic orthologs were found in single orthogroups, but 40% of syntenic paralogs can be found across multiple orthogroups. These findings suggest that orthology inference algorithms, most of which rely on clustering based on sequence similarity, will likely be prone to errors and introduce greater variation when higher ploidy species are included in orthogroup inferences. In our study, we found 7204-11230 single-copy orthogroups (depending on the algorithm used) across all five diploid species that span a range of divergence times (Appendix S3); this number is likely to be reduced if additional species were included. Additionally, our results are reflective of the predictions about greater variation when including species with higher ploidy – while there is no “ground-truth” to compare the results, in the pairwise comparison between algorithm results, there were fewer identical orthogroups between orthology algorithms, although the average degree of similarity was not very different (Fig. 4).

Ploidy can be inferred from orthogroup analyses based on the majority number of genes per species in an orthogroup. For instance, *Brassica* and *Sinapis* are mesopolyploids with the majority of the orthogroups containing either one or two gene copies, indicative of genome fractionation after the Brassiceae tribe-specific whole-genome triplication event (Yang et al., 2023; Fig. 1). Similarly, the majority of the orthogroups contain three *Camelina* gene copies, reflecting its polyploid origin and that it has not undergone extensive genome fractionation since polyploidization (Fig. 3C; Kagale et al., 2014; Mandáková et al., 2019).

Our case study of YABBY genes indicates some of the strengths and limitations of genome-wide orthology inference. For instance, the *Brassica rapa* homologs recovered for each YABBY are the same with those found in a phylogenetic study on the YABBY gene family (Lu et al., 2021). We were unable to recover the *Aethionema* ortholog of YAB3 (AT4G00180.1; Fig 5), but the phylogenetic study considered Aa31LG2G250 as the YAB3 ortholog. Interestingly, Aa31LG2G250 sequence is more similar to *Cleome violacea*, which is sister to the Brassicaceae. This finding indicates that there are subtleties in the gene family evolution that can only be uncovered with a more thorough investigation into those specific genes and a broader sampling of species. A closer look at the alignments and gene phylogeny can also reveal whether certain gene copies, especially paralogous copies that might have additional protein variation, should be included in an orthogroup. Some of this variation could occur from improper gene annotation, misalignment, or choosing an alternative primary transcript to represent the gene copy, and additional tools may provide complementary solutions. For example, NovelTree (Celebi et al., 2023) can improve orthogroup assignments by trimming unaligned sequences. Additionally, Broccoli uses k-mer clustering in its workflow to group regions of the protein within a species and can assign a protein to multiple orthogroups, allowing the detection of chimeric proteins.

## AUTHOR CONTRIBUTIONS

I.T.L. and L.A.N. conceived of the project. I.T.L. performed the analyses. All authors wrote, read, and reviewed the manuscript.

## ACKNOWLEDGEMENTS

The authors thank Dr. Bryan Piatkowski for early discussions regarding orthology and algorithms and Drs. Philip Shushkov and Matthew W. Hahn for helpful feedback on the manuscript. I.T.L. was supported by NSF Postdoctoral Research Fellowship in Biology (DBI – 2010944). LN is supported by startup funds from UCLA and Indiana University.

## DATA AVAILABILITY STATEMENT

Scripts and specific output files are openly available on GitHub (https://github.com/itliao/BrassicaceaeOrthology) and Dryad (https://doi.org/10.5061/dryad.8sf7m0cw8). Supporting data are provided in the Supporting Information.

## SUPPORTING INFORMATION

Additional Supporting Information may be found online in the Supporting Information section at the end of the article.

Appendix S1: Most orthogroups contain the maximum number of species across the orthology algorithms tested.

Appendix S2: Summary statistics describing the number of species in an orthogroup across the methods tested.

Appendix S3: Orthogroups with single copy genes with all species represented for the diploid species set from all methods.

Appendix S4: The average number of species in an orthogroup is generally similar across the algorithms, except for OrthNet.

Appendix S5 Comparison of the number of species per orthogroup detected across orthology interference methods, except OrthNet.

Appendix S6: Distributions of the number of genes per species found in an orthogroup reflect the predicted ploidy of the species.

Appendix S7: Stacked bar plots and heatmaps infer the ploidy of the species by displaying the number of genes per species in each orthogroup.

Appendix S8: Testing differences in the number of genes for each species per orthogroup across methods.

Appendix S9: Orthogroup gene compositions are more similar across algorithms tested for diploid species than for those from higher ploidy species.

Appendix S10: Summary statistics for metrics calculated for all-against-all comparisons of orthogroup compositions among all algorithms.

Appendix S11: Distribution of pairwise comparisons between orthology inference algorithms.

Appendix S12: Proportion of predicted orthology relationships between all species pairs across algorithms for the diploid set and the diploid+higher ploidy set.

Appendix S13: Species pair ortholog ratios across algorithms.

Appendix S14: The Jaccard Index was calculated from orthology inference results from each algorithm compared to the baseline orthology inference results (OrthoFinder-BLAST-MCL) for each species pair.

Appendix S15: Species pair orthogroup composition comparisons between the orthology inference algorithms and the baseline orthology algorithm (OrthoFinder-BLAST-MCL) using the Jaccard Index.

Appendix S16: RAxML tree of all genes from the six Arabidopsis YABBY orthogroups, after 1000 bootstraps.

Appendix S17: Screenshots of YABBY sequence alignments reveal sequence features that could affect whether an orthology inference algorithm incorporates the sequence into an orthogroup.

Appendix S18: Orthogroup composition outputs of select YABBYs from the diploid set

