## Supplementary Information for "Different orthology inference algorithms generate similar predicted orthogroups among Brassicaceae species"

Appendix S1-Fig. S1: Most orthogroups contain the maximum number of species across the orthology algorithms tested. Distribution of the number of species found in each orthogroup for each algorithm. Each bar represents the number of species in each orthogroup, with the number of orthogroups labeled on the top of the bar. (A) Diploid set. (B) Diploid+higher ploidy set. Corresponds to Figure 2.

<table>
<tr><th>algorithm</th><th>(A) Diploid set</th><th>(B) diploid+higher ploidy set</th></tr>
<tr><td>Broccoli</td><td>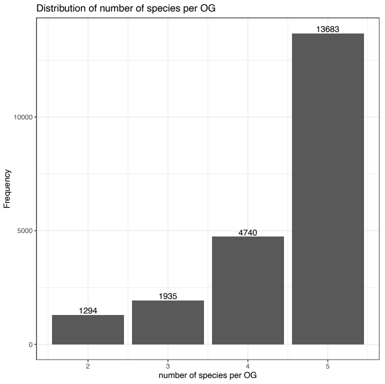
</td><td>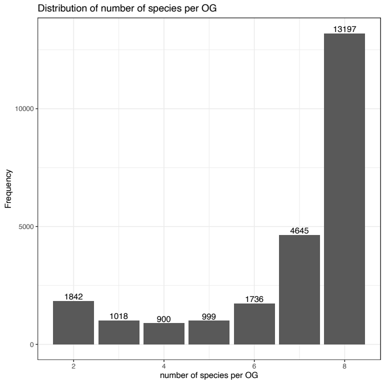
</td></tr>
<tr><td>OrthoFinder-BLAST</td><td>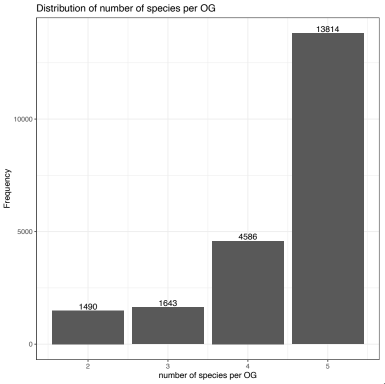
</td><td>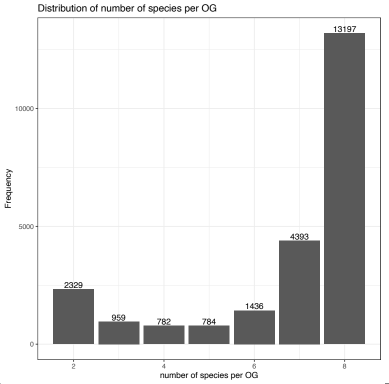
</td></tr>
<tr><td>OrthoFinder-DIAMOND</td><td>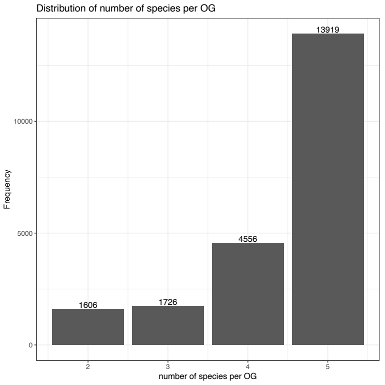
</td><td>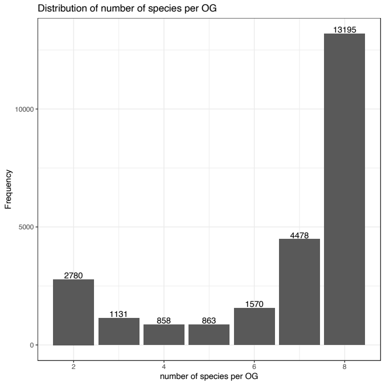
</td></tr>
</table>

OrthoFinder-MMseqs2

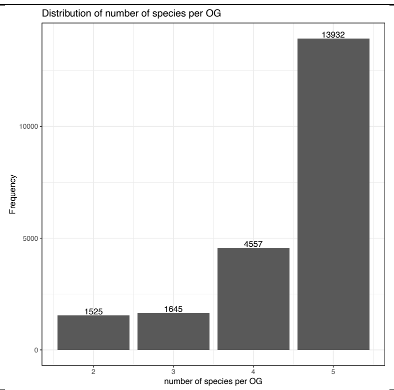

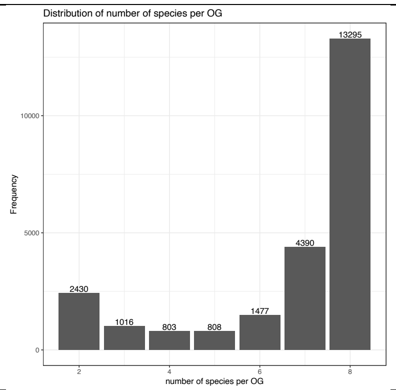

SonicParanoid-DIAMOND

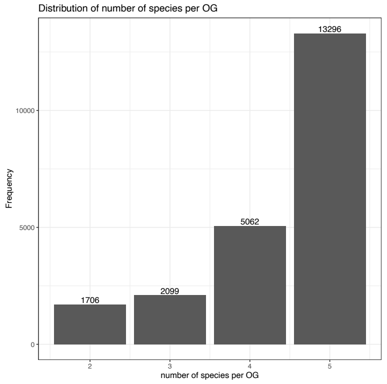

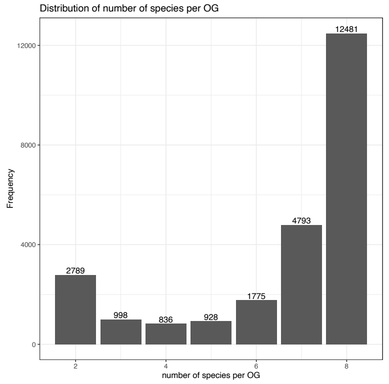

SonicParanoid-MMseqs2

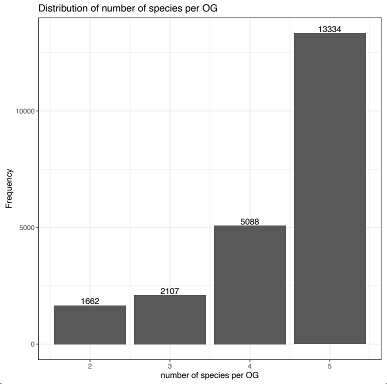

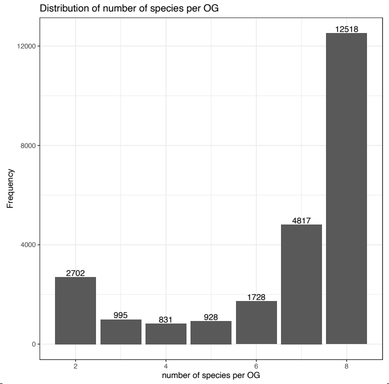

OrthNet

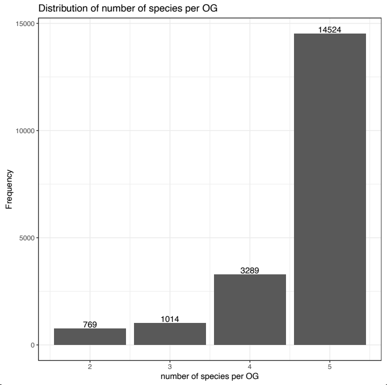

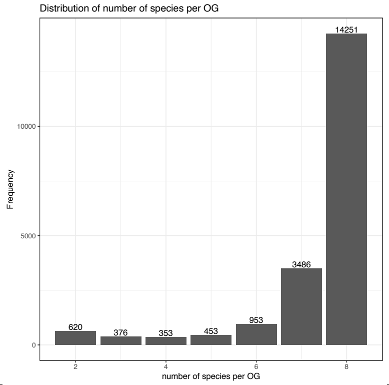

Appendix S2-Table S1: Summary statistics describing the number of species in an orthogroup across the methods tested. (A) Diploid set. (B) Diploid+higher ploidy set.

(A) Diploid set

| **algorithm** | **n** | **mean** | **median** | **sd** | **se** |
|---|---|---|---|---|---|
| Broccoli | 21652 | 4.42305561 | 5 | 0.88398712 | 0.00600754 |
| OrthoFinder-BLAST | 21533 | 4.42683323 | 5 | 0.90136787 | 0.00614257 |
| OrthoFinder-DIAMOND | 21807 | 4.41184023 | 5 | 0.91784571 | 0.00621544 |
| OrthoFinder-MMseqs2 | 21659 | 4.42647398 | 5 | 0.90498001 | 0.00614922 |
| OrthNet | 19596 | 4.61094101 | 5 | 0.75938676 | 0.00542474 |
| SonicParanoid-DIAMOND | 22163 | 4.35126111 | 5 | 0.93764707 | 0.00629833 |
| SonicParanoid-MMseqs2 | 22191 | 4.35613537 | 5 | 0.93199264 | 0.00625639 |
| (B) diploid+higher ploidy set | | | | | |
| **algorithm** | **n** | **mean** | **median** | **sd** | **se** |
| Broccoli | 24337 | 6.732136253 | 8 | 1.897487831 | 0.012163134 |
| OrthoFinder-BLAST | 23880 | 6.680318258 | 8 | 2.004387641 | 0.012970734 |
| OrthoFinder-DIAMOND | 24875 | 6.553809045 | 8 | 2.089332484 | 0.013247259 |
| OrthoFinder-MMseqs2 | 24219 | 6.652297783 | 8 | 2.025000646 | 0.01301209 |
| OrthNet | 20492 | 7.328372048 | 8 | 1.388396747 | 0.009698876 |
| SonicParanoid-DIAMOND | 24600 | 6.528658537 | 8 | 2.073096085 | 0.013217578 |
| SonicParanoid-MMseqs2 | 24519 | 6.549369876 | 8 | 2.058845412 | 0.013148383 |

Appendix S3-Table S2: Orthogroups with single copy genes with all species represented for the diploid species set from all methods. Each row represents one of the seven orthology inference algorithms tested. (A) Diploid set. (B) Diploid+higher ploidy set.

(A) Diploid set

| **Algorithm** | **Orthogroups with exactly 1 copy per species** | **Total number of orthogroups with all 5 species represented** | **Total number of orthogroups** | **Proportion of orthogroups with all species represented** | **Proportion of orthogroups with exactly 1 gene copy** |
|---|---|---|---|---|---|
| **Broccoli** | 10044 | 13683 | 21652 | 0.632 | 0.734 |
| **OrthoFinder-BLAST** | 8607 | 13814 | 21533 | 0.642 | 0.623 |
| **OrthoFinder-DIAMOND** | 8880 | 13919 | 21807 | 0.638 | 0.638 |
| **OrthoFinder-MMseqs2** | 8746 | 13932 | 21659 | 0.643 | 0.628 |
| **OrthNet** | 7204 | 14524 | 19596 | 0.741 | 0.496 |
| **SonicParanoid-DIAMOND** | 11146 | 13296 | 22163 | 0.600 | 0.838 |
| **SonicParanoid -MMseqs2** | 11230 | 13334 | 22191 | 0.601 | 0.842 |

(B) diploid+higher ploidy set

| | **Total number of orthogroups with all 8 species represented** | **Total number of orthogroups** | **Proportion of orthogroups with all species represented** |
|---|---|---|---|
| **Broccoli** | 13197 | 24337 | 0.542 |
| **OrthoFinder-BLAST** | 13197 | 23880 | 0.553 |
| **OrthoFinder-DIAMOND** | 13195 | 24875 | 0.530 |
| **OrthoFinder-MMseqs2** | 13295 | 24219 | 0.549 |
| **OrthNet** | 14251 | 20492 | 0.695 |
| **SonicParanoid-DIAMOND** | 12481 | 24600 | 0.507 |
| **SonicParanoid -MMseqs2** | 12581 | 24519 | 0.513 |

Appendix S4-Fig. S2: The average number of species in an orthogroup is generally similar across the algorithms, except for OrthNet. (A) Diploid set. (B) Diploid+higher ploidy set. BR: Broccoli. OF_blast: OrthoFinder-BLAST. OF_diamond: OrthoFinder-DIAMOND. OF_mmseqs: OrthoFinder-MMseqs2. SP_diamond: SonicParanoid-DIAMOND. SP_mmseqs: SonicParanoid-MMseqs2. ON: OrthNet.

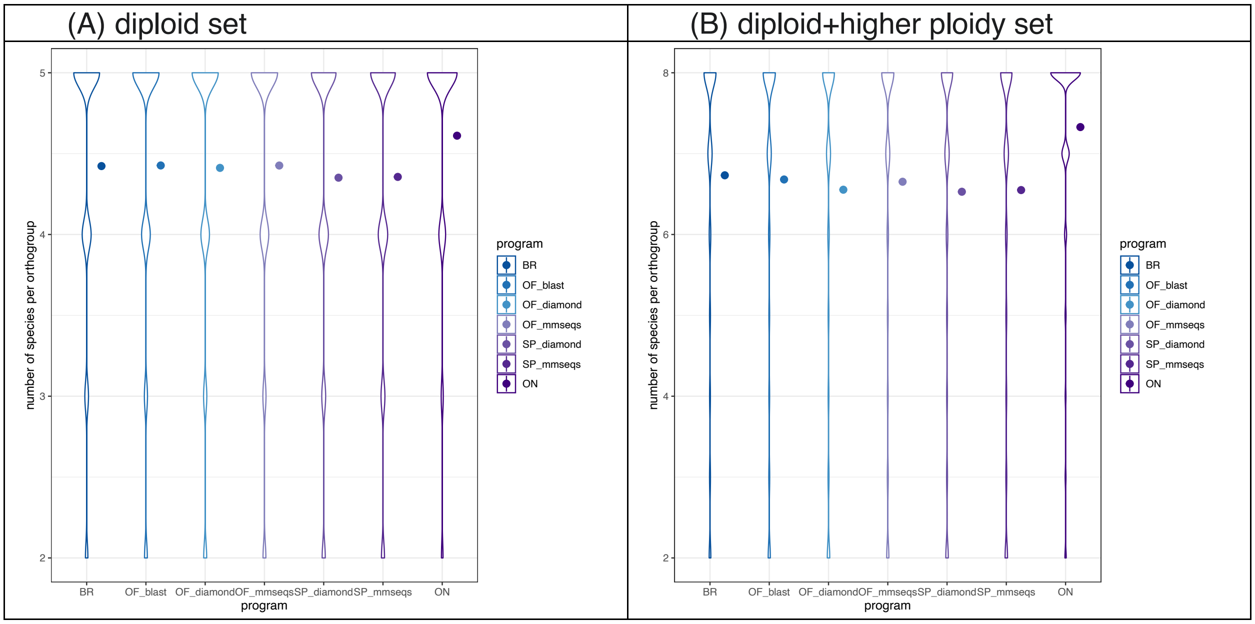

Appendix S5-Table S3: Comparison of the number of species per orthogroup detected across orthology interference methods, except OrthNet. Gray highlight indicate significance at $P < 0.05$ after a FDR adjustment for multiple comparisons. (A) Diploid set. (B) Diploid+higher ploidy set. Top table: results from a Kruskal-Wallis rank sum test. Bottom table: p-values from all possible pairwise comparisons of the methods tested through a Wilcoxon rank sum test after a FDR correction. chi.squared is the test statistic used to calculate the p-value. BR: Broccoli. OF_blast: OrthoFinder-BLAST. OF_diamond: OrthoFinder-DIAMOND. OF_mmseqs: OrthoFinder-MMseqs2. SP_diamond: SonicParanoid-DIAMOND. SP_mmseqs: SonicParanoid-MMseqs2. ON: OrthNet. Corresponds to Fig. 2.

(A) Diploid Set

Kruskal-Wallis rank sum test

| | number of species per orthogroup | | |
|---|---|---|---|
| | chi.squared | df | p.value |
| method | 197.37 | 5 | <2.20E-16 |

Wilcoxon rank sum test

| | BR | OF_blast | OF_diamond | OF_mmseqs | SP_diamond |
|---|---|---|---|---|---|
| OF_blast | 0.15 | - | - | - | - |
| OF_diamond | 0.74 | 0.3 | - | - | - |
| OF_mmseqs | 0.1 | 0.83 | 0.23 | - | - |
| SP_diamond | 3.50E-14 | < 2e-16 | 2.40E-15 | < 2e-16 | - |
| SP_mmseqs | 4.40E-13 | < 2e-16 | 3.10E-14 | < 2e-16 | 0.77 |

(B) Diploid+higher ploidy set

| | number of species per orthogroup | | |
|---|---|---|---|
| | chi.squared | df | p.value |
| method | 206.51 | 5 | <2.20E-16 |

Wilcoxon rank sum test

| | BR | OF_blast | OF_diamond | OF_mmseqs | SP_diamond |
|---|---|---|---|---|---|
| OF_blast | 0.63801 | - | - | - | - |
| OF_diamond | 5.10E-09 | 1.20E-09 | - | - | - |
| OF_mmseqs | 0.4523 | 0.27762 | 1.00E-06 | - | - |
| SP_diamond | < 2e-16 | < 2e-16 | 2.40E-04 | < 2e-16 | - |
| SP_mmseqs | < 2e-16 | < 2e-16 | 7.22E-03 | 1.20E-14 | 0.36689 |

Appendix S6-Fig. S3: Distributions of the number of genes per species found in an orthogroup reflect the predicted ploidy of the species. Stacked bar plots displaying species from (A) diploid set and (B) diploid+higher ploidy set. Each plot displays the results for one species across the different algorithms, and each color represents the specific number of genes per species found in an orthogroup. BR: Broccoli. OF_blast: OrthoFinder-BLAST. OF_diamond: OrthoFinder-DIAMOND. OF_mmseqs: OrthoFinder-MMseqs2. SP_diamond: SonicParanoid-DIAMOND. SP_mmseqs: SonicParanoid-MMseqs2. ON: OrthNet. Corresponds with Figure 3.

| (A) Diploid | (B) diploid+higher ploidy set |
|---|---|

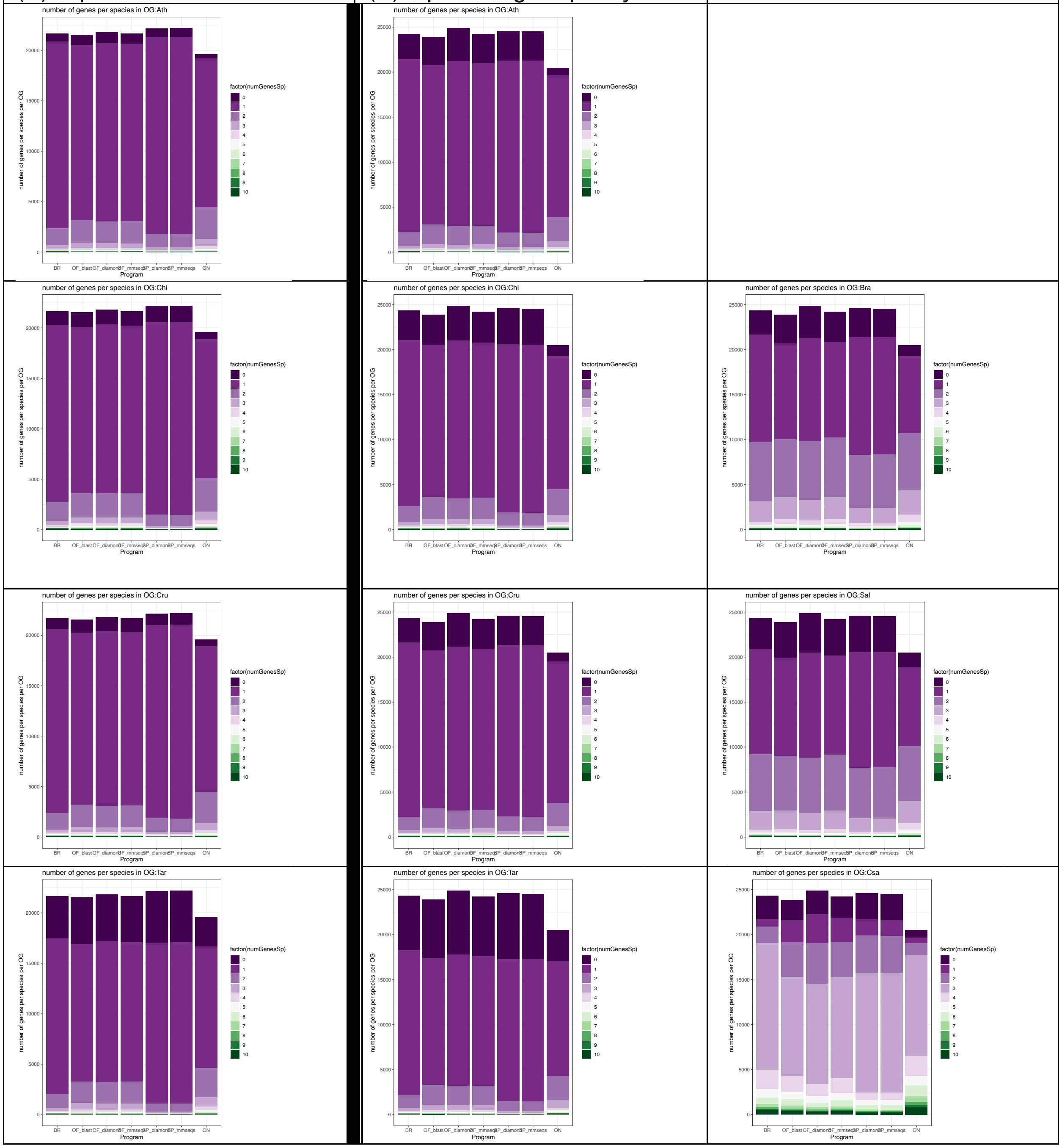

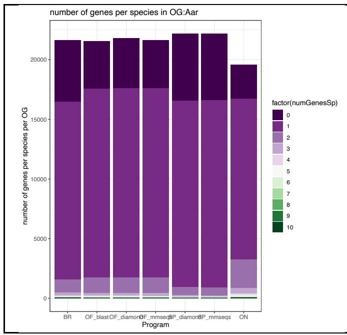
number of genes per species in OG:Aar
number of genes per species per OG
0
5000
10000
15000
20000
BR
OF_blast
SP_mmseqs
ON
Program
factor(numGenesSp)
0
1
2
3
4
5
6
7
8
9
10

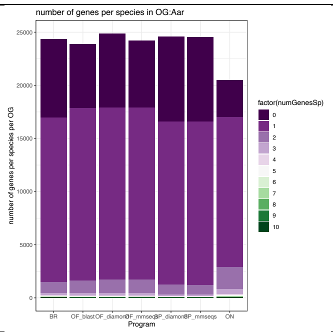
number of genes per species in OG:Aar
number of genes per species per OG
0
5000
10000
15000
20000
25000
BR
OF_blast
SP_mmseqs
ON
Program
factor(numGenesSp)
0
1
2
3
4
5
6
7
8
9
10

Appendix S7-Fig. S4: Stacked bar plots and heatmaps infer the ploidy of the species by displaying the number of genes per species in each orthogroup. All vertical bars (stacked bars) or columns (heatmap) represent a species. For stacked bars, colors indicate the number of genes found in a given orthogroup, with “10” representing orthogroups with 10 or more genes. For heatmaps, the colors represent the number of orthogroups with a certain number of genes with the numbers also listed in the rectangles. (A) Diploid set. (B) Diploid+higher ploidy set. Each row represents the output from an algorithm.

| algorithm | (A) Diploid set | (B) diploid+higher ploidy set |
|---|---|---|
| Broccoli | | |

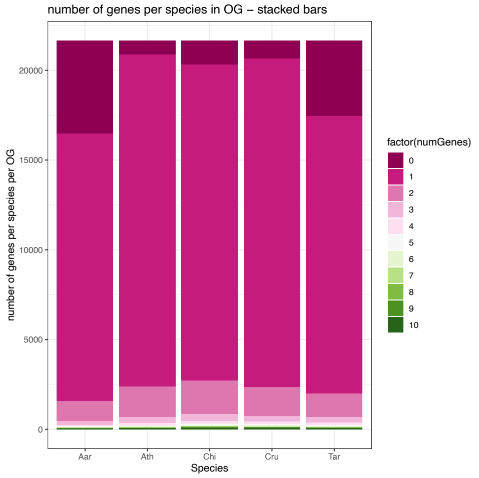

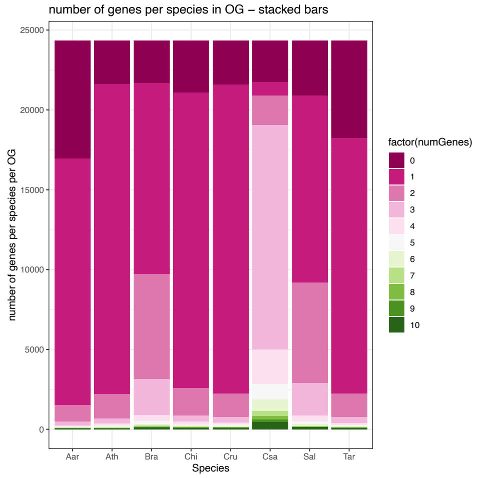

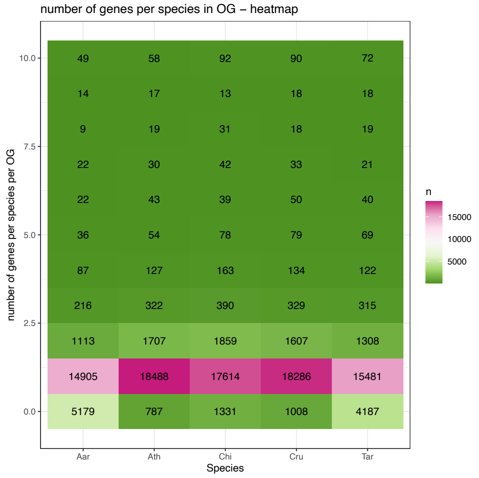

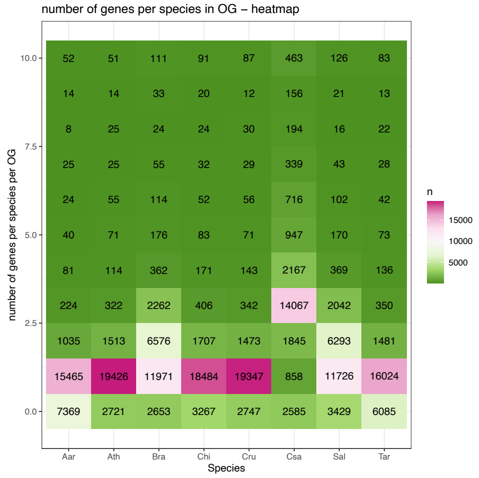

OrthoFinder-BLAST

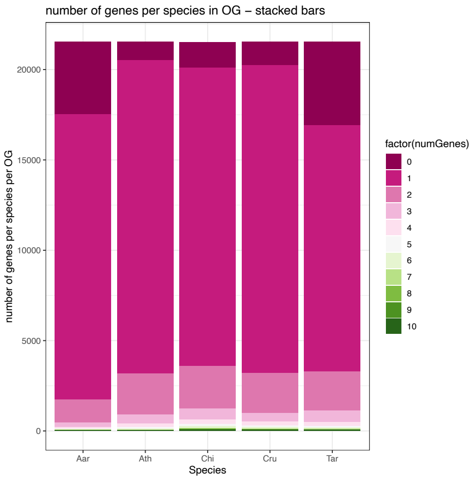

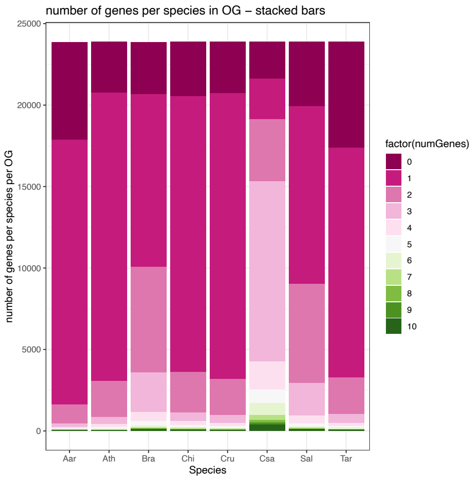

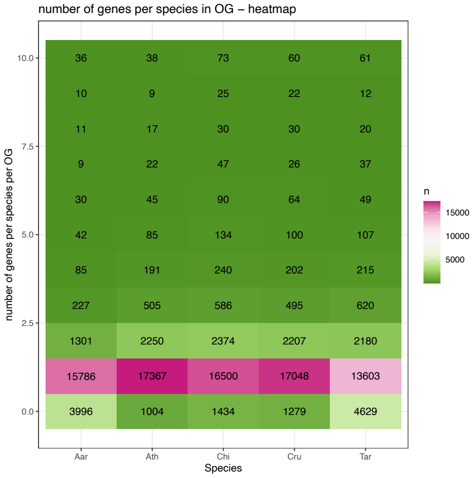

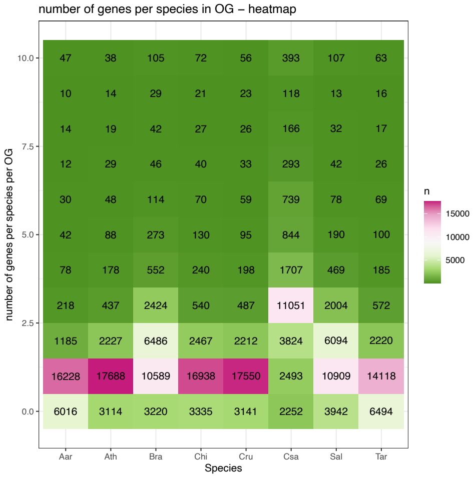

### OrthoFinder-DIAMOND

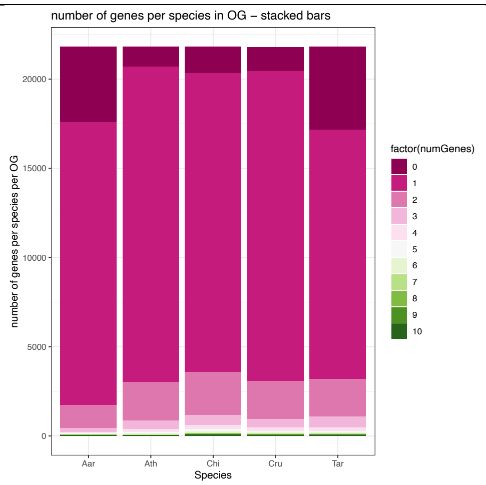

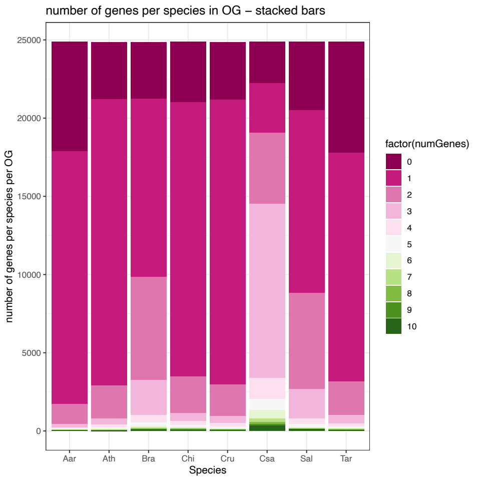

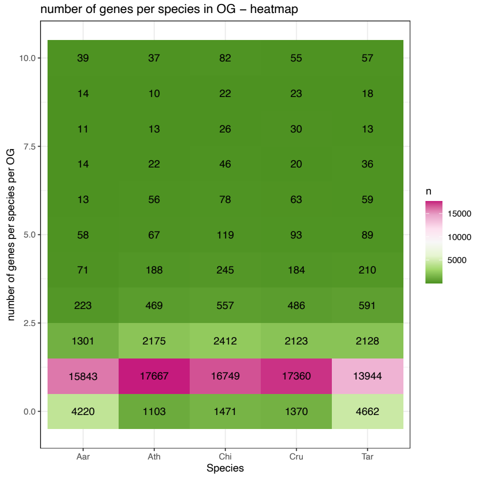

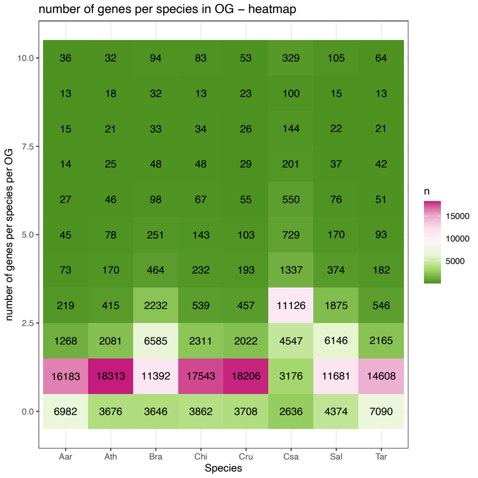

OrthoFinder-MMseqs2

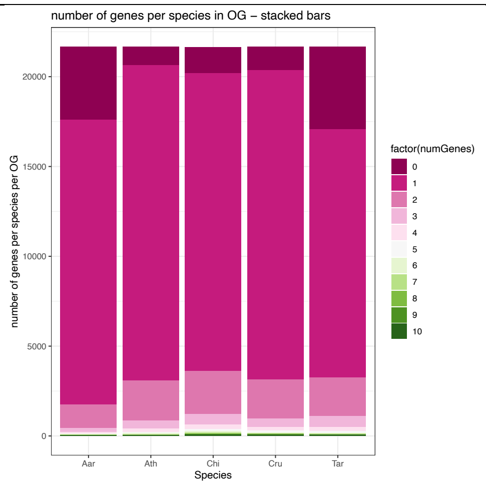

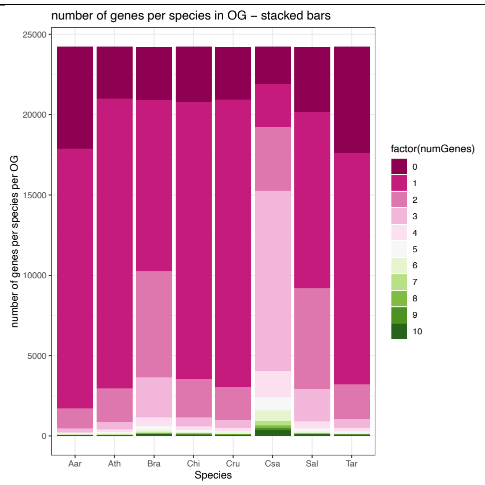

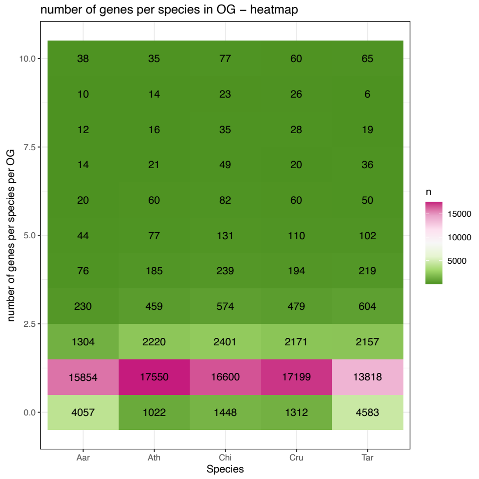

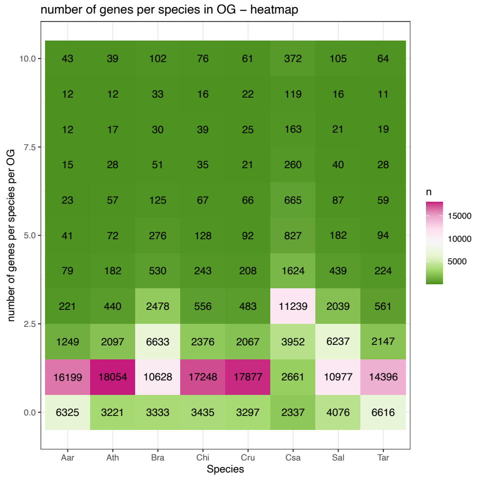

SonicParanoid
-DIAMOND

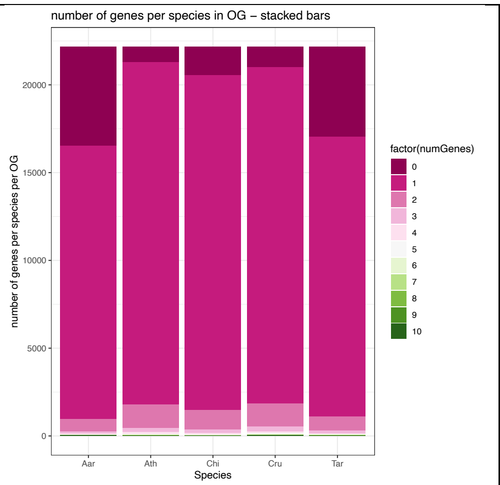

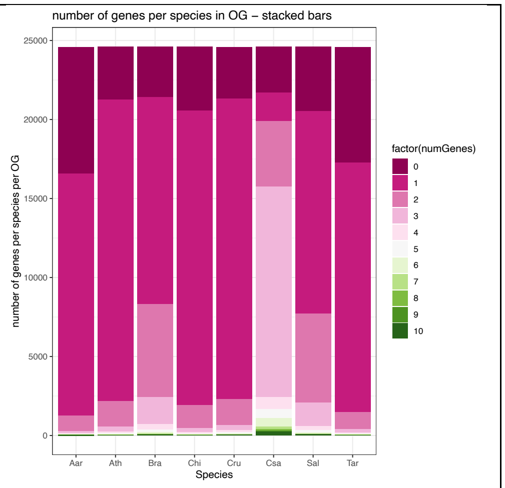

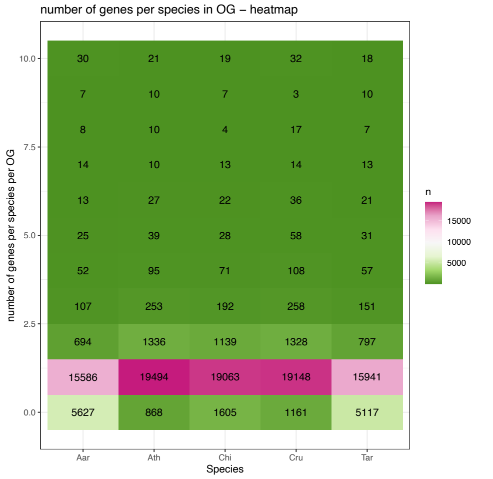

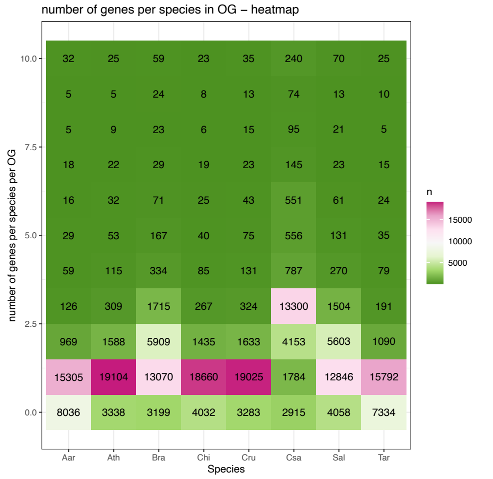

SonicParanoid
-MMseqs2

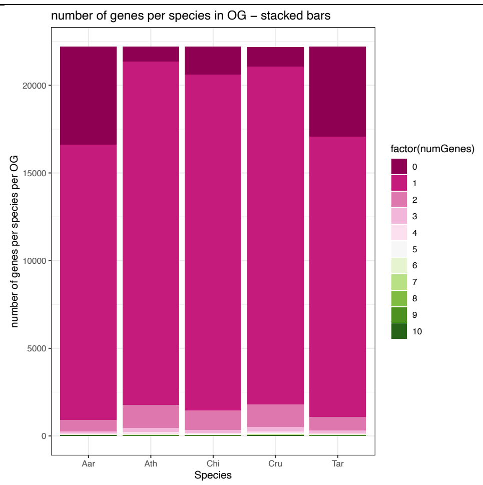

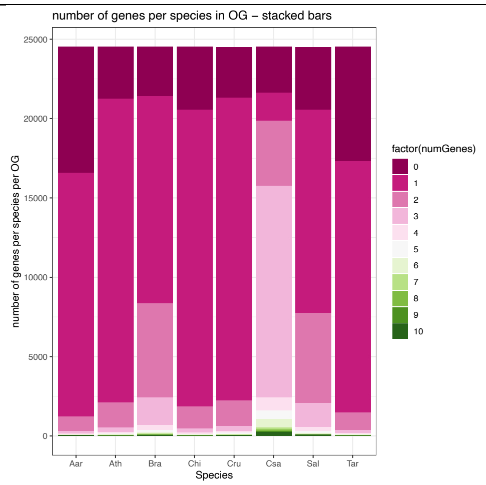

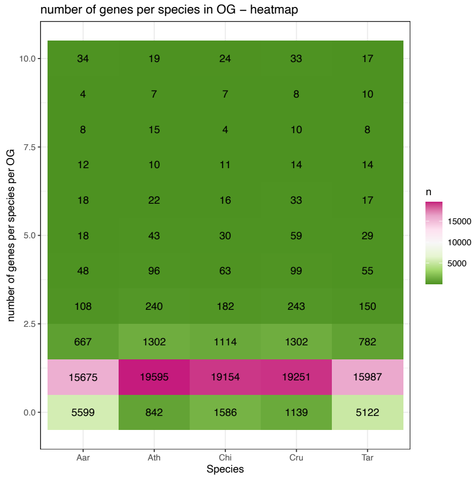

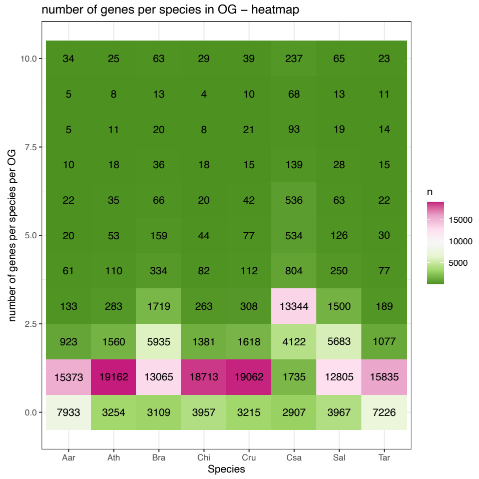

OrthNet

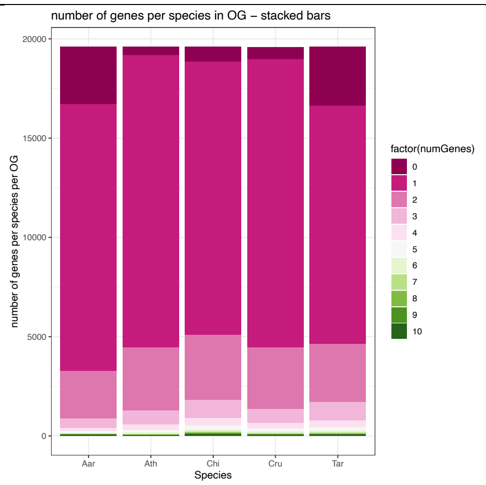
number of genes per species in OG – stacked bars
number of genes per species per OG
20000
15000
10000
5000
0
Aar
Ath
Chi
Cru
Tar
Species
factor(numGenes)
0
1
2
3
4
5
6
7
8
9
10

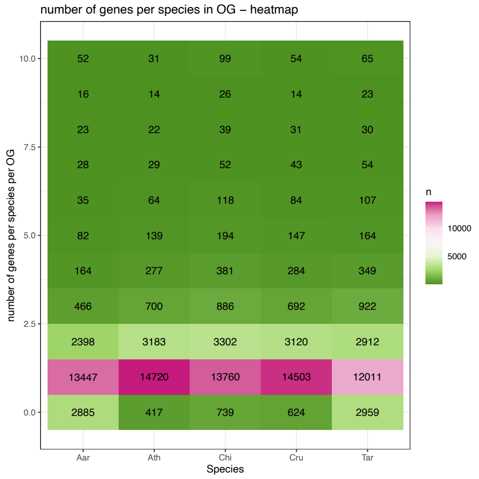
number of genes per species in OG – heatmap
number of genes per species per OG
10.0
7.5
5.0
2.5
0.0
52 31 99 54 65
16 14 26 14 23
23 22 39 31 30
28 29 52 43 54
35 64 118 84 107
82 139 194 147 164
164 277 381 284 349
466 700 886 692 922
2398 3183 3302 3120 2912
13447 14720 13760 14503 12011
2885 417 739 624 2959
Aar
Ath
Chi
Cru
Tar
Species
n
10000
5000

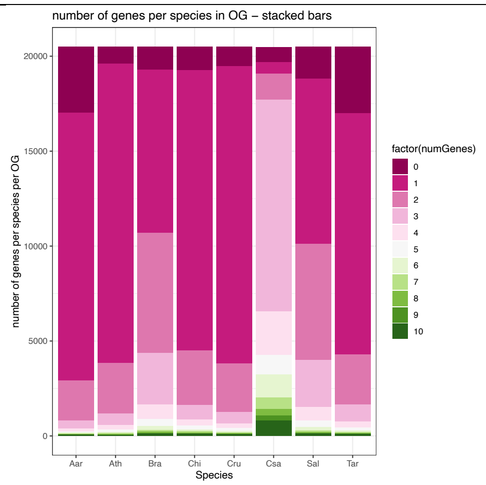
number of genes per species in OG – stacked bars
number of genes per species per OG
20000
15000
10000
5000
0
Aar
Ath
Bra
Chi
Cru
Csa
Sal
Tar
Species
factor(numGenes)
0
1
2
3
4
5
6
7
8
9
10

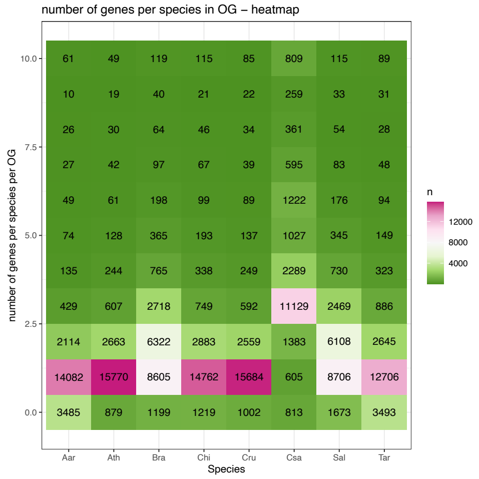
number of genes per species in OG – heatmap
number of genes per species per OG
10.0
7.5
5.0
2.5
0.0
61 49 119 115 85 809 115 89
10 19 40 21 22 259 33 31
26 30 64 46 34 361 54 28
27 42 97 67 39 595 83 48
49 61 198 99 89 1222 176 94
74 128 365 193 137 1027 345 149
135 244 765 338 249 2289 730 323
429 607 2718 749 592 11129 2469 886
2114 2663 6322 2883 2559 1383 6108 2645
14082 15770 8605 14762 15684 605 8706 12706
3485 879 1199 1219 1002 813 1673 3493
Aar
Ath
Bra
Chi
Cru
Csa
Sal
Tar
Species
n
12000
8000
4000

Appendix S8-Table S4: Testing differences in the number of genes for each species per orthogroup across methods. For each species, Kruskal-Wallis rank sum test was performed across all methods, and Wilcoxon rank sum test on all pairwise methods. Gray highlight indicates significance at P < 0.05 after a FDR adjustment for multiple comparisons. (A) Diploid set. (B) Diploid+higher ploidy set. BR: Broccoli. OF_blast: OrthoFinder-BLAST. OF_diamond: OrthoFinder-DIAMOND. OF_mmseqs: OrthoFinder-MMseqs2. ON: OrthNet. SP_diamond: SonicParanoid-DIAMOND. SP_mmseqs: SonicParanoid-MMseqs2.

(A) Diploid set

***Arabidopsis***

Kruskal-Wallis rank sum test

number of genes per species per orthogroup

| | chi.squared | df | p.value |
|---|---|---|---|
| program | 2400.4 | 6 | <2.20E-16 |

Wilcoxon rank sum test

| | BR | OF_blast | OF_diamond | OF_mmseqs | ON | SP_diamond |
|---|---|---|---|---|---|---|
| OF_blast | 3.85E-13 | NA | NA | NA | NA | NA |
| OF_diamond | 1.75E-05 | 0.00586652 | NA | NA | NA | NA |
| OF_mmseqs | 2.07E-09 | 0.22805291 | 0.12564797 | NA | NA | NA |
| ON | 1.77E-227 | 1.22E-125 | 1.28E-155 | 2.16E-139 | NA | NA |
| SP_diamond | 2.17E-20 | 1.54E-58 | 3.28E-39 | 4.73E-50 | 0 | NA |
| SP_mmseqs | 6.21E-22 | 2.98E-61 | 2.50E-41 | 1.56E-52 | 0 | 0.74271203 |

***Cardamine***

Kruskal-Wallis rank sum test

number of genes per species per orthogroup

| | chi.squared | df | p.value |
|---|---|---|---|
| program | 3937.9 | 6 | <2.20E-16 |

Wilcoxon rank sum test

| | BR | OF_blast | OF_diamond | OF_mmseqs | ON | SP_diamond |
|---|---|---|---|---|---|---|
| OF_blast | 3.58E-19 | NA | NA | NA | NA | NA |
| OF_diamond | 4.97E-16 | 0.45378406 | NA | NA | NA | NA |
| OF_mmseqs | 1.52E-18 | 0.85832543 | 0.54626378 | NA | NA | NA |
| ON | 1.06E-266 | 6.11E-140 | 1.60E-150 | 2.40E-142 | NA | NA |
| SP_diamond | 2.19E-73 | 1.26E-153 | 8.35E-145 | 5.34E-152 | 0 | NA |
| SP_mmseqs | 1.57E-76 | 3.53E-158 | 2.51E-149 | 1.50E-156 | 0 | 0.78491972 |

***Capsella***

| | Kruskal-Wallis rank sum test number of genes per species per orthogroup | | |
|---|---|---|---|
| | chi.squared | df | p.value |
| program | 2214.5 | 6 | <2.20E-16 |

Wilcoxon rank sum test

| | BR | OF_blast | OF_diamond | OF_mmseqs | ON | SP_diamond |
|---|---|---|---|---|---|---|
| OF_blast | 2.09E-12 | NA | NA | NA | NA | NA |
| OF_diamond | 2.16E-05 | 0.00996965 | NA | NA | NA | NA |
| OF_mmseqs | 4.47E-09 | 0.28173649 | 0.14067245 | NA | NA | NA |
| ON | 6.94E-216 | 6.75E-118 | 1.38E-145 | 2.19E-129 | NA | NA |
| SP_diamond | 2.67E-18 | 6.82E-53 | 6.63E-36 | 1.96E-45 | 0 | NA |
| SP_mmseqs | 3.37E-20 | 3.61E-56 | 1.60E-38 | 1.95E-48 | 0 | 0.65851822 |

***Thlaspi***

| | Kruskal-Wallis rank sum test number of genes per species per orthogroup | | |
|---|---|---|---|
| | chi.squared | df | p.value |
| program | 2966.3 | 6 | <2.20E-16 |

Wilcoxon rank sum test

| | BR | OF_blast | OF_diamond | OF_mmseqs | ON | SP_diamond |
|---|---|---|---|---|---|---|
| OF_blast | 1.04E-10 | NA | NA | NA | NA | NA |
| OF_diamond | 2.32E-08 | 0.41611927 | NA | NA | NA | NA |
| OF_mmseqs | 3.13E-11 | 0.90506722 | 0.3636386 | NA | NA | NA |
| ON | 3.03E-233 | 1.35E-130 | 1.89E-142 | 2.85E-131 | NA | NA |
| SP_diamond | 4.99E-55 | 1.76E-96 | 4.60E-90 | 3.56E-99 | 0 | NA |
| SP_mmseqs | 2.02E-56 | 3.00E-98 | 8.71E-92 | 5.62E-101 | 0 | 0.8972677 |

***Aethionema***

| | Kruskal-Wallis rank sum test number of genes per species per orthogroup | | |
|---|---|---|---|
| | chi.squared | df | p.value |
| program | 2742 | 6 | <2.20E-16 |

Wilcoxon rank sum test

| | BR | OF_blast | OF_diamond | OF_mmseqs | ON | SP_diamond |
|---|---|---|---|---|---|---|
| OF_blast | 5.05E-37 | NA | NA | NA | NA | NA |
| OF_diamond | 2.91E-27 | 0.05990035 | NA | NA | NA | NA |
| OF_mmseqs | 1.59E-34 | 0.65422795 | 0.15953662 | NA | NA | NA |
| ON | 1.14E-247 | 5.36E-117 | 1.78E-134 | 1.32E-121 | NA | NA |
| SP_diamond | 4.98E-16 | 2.69E-103 | 1.34E-85 | 5.35E-99 | 0 | NA |
| SP_mmseqs | 7.24E-16 | 2.69E-103 | 1.45E-85 | 5.35E-99 | 0 | 0.93248936 |

(B) Diploid+higher ploidy set

***Aethionema***

Kruskal-Wallis rank sum test

number of genes per species per orthogroup

| | chi.squared | df | p.value |
|---|---|---|---|
| program | 2920.4 | 6 | <2.20E-16 |

Wilcoxon rank sum test

| | BR | OF_blast | OF_diamond | OF_mmseqs | ON | SP_diamond |
|---|---|---|---|---|---|---|
| OF_blast | 1.37E-32 | NA | NA | NA | NA | NA |
| OF_diamond | 8.24E-09 | 9.13E-10 | NA | NA | NA | NA |
| OF_mmseqs | 4.18E-24 | 0.08489463 | 1.19E-05 | NA | NA | NA |
| ON | 0 | 1.35E-187 | 6.03E-261 | 2.13E-206 | NA | NA |
| SP_diamond | 1.70E-11 | 9.08E-79 | 1.86E-36 | 6.27E-65 | 0 | NA |
| SP_mmseqs | 2.25E-10 | 6.33E-76 | 1.96E-34 | 2.63E-62 | 0 | 0.67915531 |

***Arabidopsis***

Kruskal-Wallis rank sum test

number of genes per species per orthogroup

| | chi.squared | df | p.value |
|---|---|---|---|
| program | 2441.9 | 6 | <2.20E-16 |

Wilcoxon rank sum test

| | BR | OF_blast | OF_diamond | OF_mmseqs | ON | SP_diamond |
|---|---|---|---|---|---|---|
| OF_blast | 1.27E-05 | NA | NA | NA | NA | NA |
| OF_diamond | 0.01093007 | 1.31E-10 | NA | NA | NA | NA |
| OF_mmseqs | 0.03094254 | 0.03744162 | 1.16E-05 | NA | NA | NA |
| ON | 0 | 1.97E-208 | 2.76E-305 | 3.19E-241 | NA | NA |

| | | | | | | |
|---|---|---|---|---|---|---|
| SP_diamond | 1.55E-10 | 1.44E-24 | 0.00063819 | 5.46E-16 | 0 | NA |
| SP_mmseqs | 6.56E-10 | 8.76E-24 | 0.00138369 | 2.62E-15 | 0 | 0.79169267 |

***Brassica***

Kruskal-Wallis rank sum test

number of genes per species per orthogroup

| | chi.squared | df | p.value |
|---|---|---|---|
| program | 2588.6 | 6 | <2.20E-16 |

Wilcoxon rank sum test

| | BR | OF_blast | OF_diamond | OF_mmseqs | ON | SP_diamond |
|---|---|---|---|---|---|---|
| OF_blast | 0.03115289 | NA | NA | NA | NA | NA |
| OF_diamond | 8.28E-07 | 7.82E-12 | NA | NA | NA | NA |
| OF_mmseqs | 0.0392962 | 0.89784742 | 1.64E-11 | NA | NA | NA |
| ON | 2.66E-212 | 2.98E-168 | 1.88E-264 | 4.50E-170 | NA | NA |
| SP_diamond | 7.57E-49 | 2.10E-58 | 8.00E-20 | 8.81E-58 | 0 | NA |
| SP_mmseqs | 8.50E-45 | 2.59E-54 | 2.20E-17 | 1.05E-53 | 0 | 0.50743934 |

***Cardamine***

Kruskal-Wallis rank sum test

number of genes per species per orthogroup

| | chi.squared | df | p.value |
|---|---|---|---|
| program | 3436.6 | 6 | <2.20E-16 |

Wilcoxon rank sum test

| | BR | OF_blast | OF_diamond | OF_mmseqs | ON | SP_diamond |
|---|---|---|---|---|---|---|
| OF_blast | 2.88E-16 | NA | NA | NA | NA | NA |
| OF_diamond | 0.01626824 | 6.90E-08 | NA | NA | NA | NA |
| OF_mmseqs | 1.92E-11 | 0.16594817 | 6.02E-05 | NA | NA | NA |
| ON | 0 | 6.01E-191 | 2.57E-265 | 4.71E-211 | NA | NA |
| SP_diamond | 1.39E-40 | 5.12E-96 | 4.84E-51 | 8.14E-84 | 0 | NA |
| SP_mmseqs | 3.80E-40 | 1.16E-95 | 1.41E-50 | 2.09E-83 | 0 | 0.88589981 |

***Capsella***

Kruskal-Wallis rank sum test

number of genes per species per orthogroup

| | chi.squared | df | p.value |
|---|---|---|---|
| program | 2067.7 | 6 | <2.20E-16 |

Wilcoxon rank sum test

| | BR | OF_blast | OF_diamond | OF_mmseqs | ON | SP_diamond |
|---|---|---|---|---|---|---|
| OF_blast | 1.48E-06 | NA | NA | NA | NA | NA |
| OF_diamond | 0.01874751 | 2.42E-11 | NA | NA | NA | NA |
| OF_mmseqs | 0.0358246 | 0.01275314 | 3.04E-05 | NA | NA | NA |
| ON | 1.24E-276 | 1.80E-173 | 1.56E-267 | 9.67E-209 | NA | NA |
| SP_diamond | 1.60E-06 | 2.60E-20 | 0.03322174 | 5.69E-11 | 0 | NA |
| SP_mmseqs | 2.81E-06 | 5.07E-20 | 0.04050995 | 1.02E-10 | 0 | 0.8865349 |

***Camelina***

Kruskal-Wallis rank sum test

number of genes per species per orthogroup

| | chi.squared | df | p.value |
|---|---|---|---|
| program | 7740.6 | 6 | <2.20E-16 |

Wilcoxon rank sum test

| | BR | OF_blast | OF_diamond | OF_mmseqs | ON | SP_diamond |
|---|---|---|---|---|---|---|
| OF_blast | 3.79E-127 | NA | NA | NA | NA | NA |
| OF_diamond | 0 | 1.12E-50 | NA | NA | NA | NA |
| OF_mmseqs | 2.08E-169 | 0.00068975 | 2.66E-31 | NA | NA | NA |
| ON | 5.13E-253 | 0 | 0 | 0 | NA | NA |
| SP_diamond | 0 | 4.82E-27 | 1.72E-07 | 5.89E-13 | 0 | NA |
| SP_mmseqs | 0 | 3.88E-26 | 4.74E-08 | 2.56E-12 | 0 | 0.79457526 |

***Sinapis***

Kruskal-Wallis rank sum test

number of genes per species per orthogroup

| | chi.squared | df | p.value |
|---|---|---|---|
| program | 2623.6 | 6 | <2.20E-16 |

Wilcoxon rank sum test

| | BR | OF_blast | OF_diamond | OF_mmseqs | ON | SP_diamond |
|---|---|---|---|---|---|---|
| OF_blast | 0.01133354 | NA | NA | NA | NA | NA |
| OF_diamond | 2.16E-18 | 2.64E-09 | NA | NA | NA | NA |
| OF_mmseqs | 0.00213363 | 0.58926534 | 5.30E-08 | NA | NA | NA |
| ON | 3.43E-196 | 1.61E-216 | 0 | 1.35E-224 | NA | NA |
| SP_diamond | 1.01E-54 | 1.33E-35 | 1.81E-10 | 7.03E-33 | 0 | NA |

| | | | | | | |
|---|---|---|---|---|---|---|
| SP_mmseqs | 2.20E-49 | 1.54E-31 | 2.43E-08 | 5.61E-29 | 0 | 0.40088478 |

***Thlaspi***

| | Kruskal-Wallis rank sum test | | |
|---|---|---|---|
| | number of genes per species per orthogroup | | |
| | chi.squared | df | p.value |
| program | 2935.6 | 6 | <2.20E-16 |

Wilcoxon rank sum test

| | BR | OF_blast | OF_diamond | OF_mmseqs | ON | SP_diamond |
|---|---|---|---|---|---|---|
| OF_blast | 0.00297203 | NA | NA | NA | NA | NA |
| OF_diamond | 0.29427495 | 0.00013157 | NA | NA | NA | NA |
| OF_mmseqs | 0.04176322 | 0.39341302 | 0.00297203 | NA | NA | NA |
| ON | 1.45E-261 | 1.34E-191 | 4.70E-246 | 5.40E-205 | NA | NA |
| SP_diamond | 7.98E-54 | 3.83E-68 | 3.25E-41 | 6.58E-62 | 0 | NA |
| SP_mmseqs | 9.87E-50 | 7.16E-64 | 7.85E-38 | 8.64E-58 | 0 | 0.50787134 |

Appendix S9-Fig. S5: Orthogroup gene compositions are more similar across algorithms tested for diploid species than for those from higher ploidy species. The Rand Score (RS) and the Adjusted Rand Score (ARS) was calculated for all algorithms in a pairwise manner. The upper left triangle represents the number of genes in orthogroups with the exact same gene composition (RS or ARS = 1) with the numbers in parentheses and the red color gradient representing the proportion of orthogroups with the same composition. The lower right triangle represents the mean RS value (top) or ARS value (bottom) with the numbers in parentheses representing the standard error; the gray gradient represents the mean values. (A) Diploid set. (B) Diploid+higher ploidy set. Corresponds to Appendix S10-Table S5.

| | (A) Diploid set | (B) Diploid+higher ploidy set |
|---|---|---|
| RS | | |

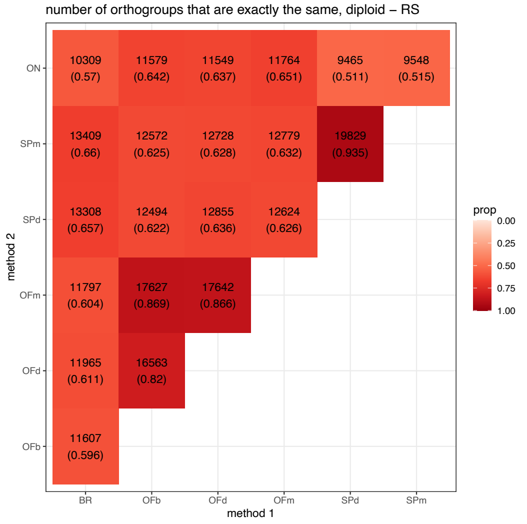

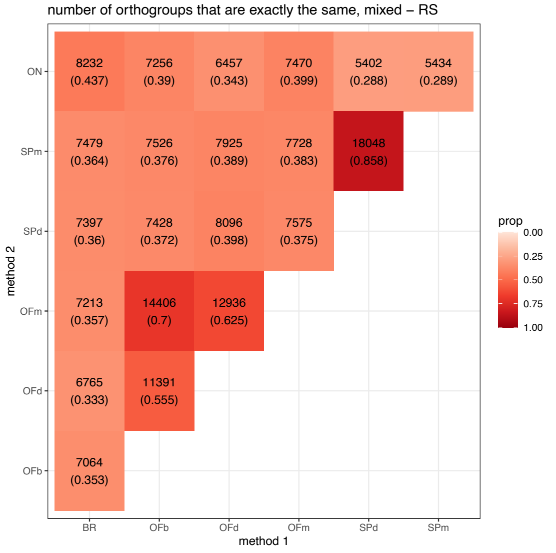

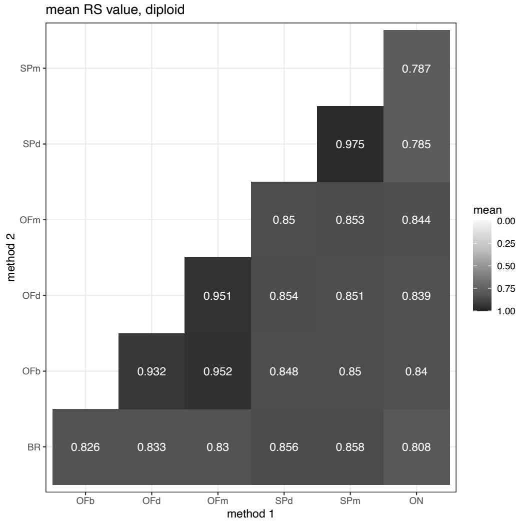

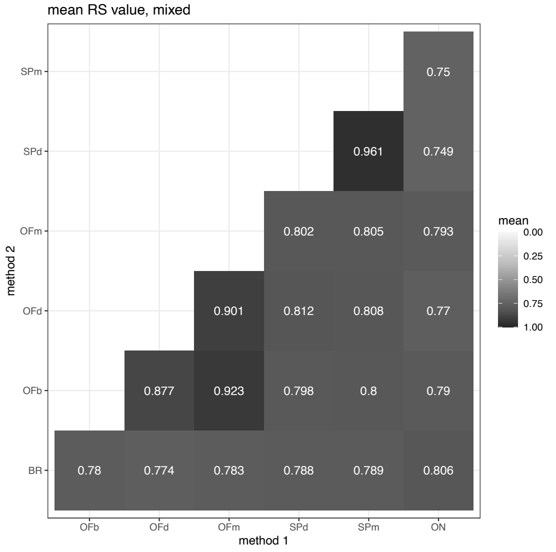

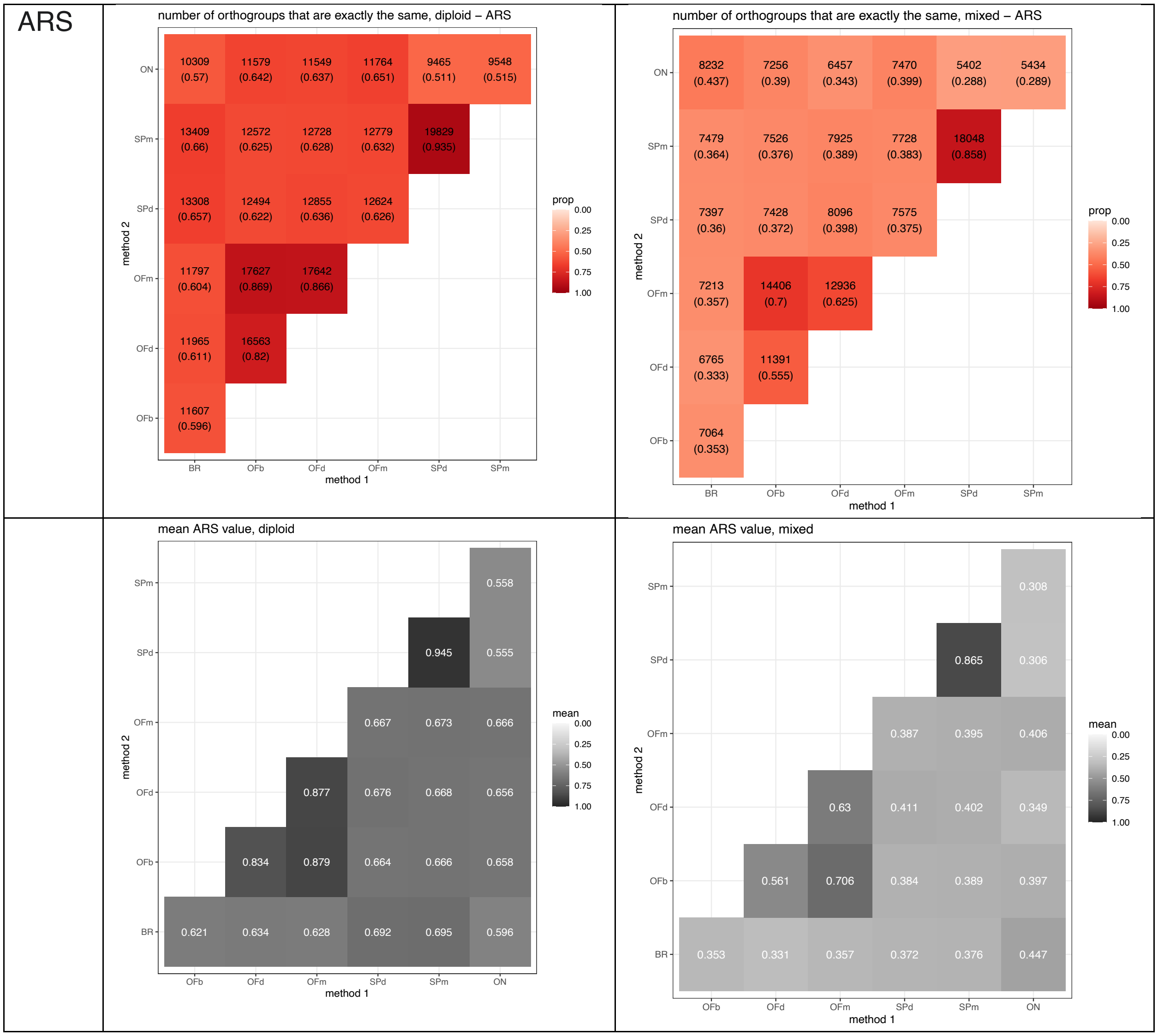
ARS
number of orthogroups that are exactly the same, diploid – ARS
ON: 10309 (0.57) 11579 (0.642) 11549 (0.637) 11764 (0.651) 9465 (0.511) 9548 (0.515)
SPm: 13409 (0.66) 12572 (0.625) 12728 (0.628) 12779 (0.632) 19829 (0.935)
SPd: 13308 (0.657) 12494 (0.622) 12855 (0.636) 12624 (0.626)
OFm: 11797 (0.604) 17627 (0.869) 17642 (0.868)
OFd: 11965 (0.611) 16563 (0.82)
OFb: 11607 (0.596)
method 1: BR OFb OFd OFm SPd SPm
method 2
prop 0.00 0.25 0.50 0.75 1.00
number of orthogroups that are exactly the same, mixed – ARS
ON: 8232 (0.437) 7256 (0.39) 6457 (0.343) 7470 (0.399) 5402 (0.288) 5434 (0.289)
SPm: 7479 (0.364) 7526 (0.376) 7925 (0.389) 7728 (0.383) 18048 (0.858)
SPd: 7397 (0.36) 7428 (0.372) 8096 (0.398) 7575 (0.375)
OFm: 7213 (0.357) 14406 (0.7) 12936 (0.625)
OFd: 6765 (0.333) 11391 (0.555)
OFb: 7064 (0.353)
method 1: BR OFb OFd OFm SPd SPm
method 2
prop 0.00 0.25 0.50 0.75 1.00
mean ARS value, diploid
SPm: 0.558
SPd: 0.945 0.555
OFm: 0.667 0.673 0.666
OFd: 0.877 0.676 0.668 0.656
OFb: 0.834 0.879 0.664 0.666 0.658
BR: 0.621 0.634 0.628 0.692 0.695 0.596
method 1: OFb OFd OFm SPd SPm ON
method 2
mean 0.00 0.25 0.50 0.75 1.00
mean ARS value, mixed
SPm: 0.308
SPd: 0.865 0.306
OFm: 0.387 0.395 0.406
OFd: 0.63 0.411 0.402 0.349
OFb: 0.561 0.706 0.384 0.389 0.397
BR: 0.353 0.331 0.357 0.372 0.376 0.447
method 1: OFb OFd OFm SPd SPm ON
method 2
mean 0.00 0.25 0.50 0.75 1.00

Appendix S10-Table S5: Summary statistics for metrics calculated for all-against-all comparisons of orthogroup compositions among all algorithms. The column “num of OG exact same” refers to instances when the metric is equal to one. (A-C) Metrics for the diploid set. (D-E) Metrics for the diploid+higher ploidy set. Corresponds to Fig. 4, Appendix S9.

(A) Diploid set - RS

| algorithm1 | algorithm2 | num of OG exact same | total num OG | proportion of OG exact same | mean | median | sd | se |
|---|---|---|---|---|---|---|---|---|
| BR | OFb | 11607 | 19471 | 0.596 | 0.826 | 1.000 | 0.239 | 0.002 |
| BR | OFd | 11965 | 19571 | 0.611 | 0.833 | 1.000 | 0.236 | 0.002 |
| BR | OFm | 11797 | 19530 | 0.604 | 0.830 | 1.000 | 0.237 | 0.002 |
| BR | ON | 10309 | 18092 | 0.570 | 0.808 | 1.000 | 0.251 | 0.002 |
| BR | SPd | 13308 | 20248 | 0.657 | 0.856 | 1.000 | 0.229 | 0.002 |
| BR | SPm | 13409 | 20303 | 0.660 | 0.858 | 1.000 | 0.228 | 0.002 |
| OFb | OFd | 16563 | 20208 | 0.820 | 0.932 | 1.000 | 0.163 | 0.001 |
| OFb | OFm | 17627 | 20273 | 0.869 | 0.952 | 1.000 | 0.139 | 0.001 |
| OFb | ON | 11579 | 18026 | 0.642 | 0.840 | 1.000 | 0.237 | 0.002 |
| OFb | SPd | 12494 | 20093 | 0.622 | 0.848 | 1.000 | 0.227 | 0.002 |
| OFb | SPm | 12572 | 20131 | 0.625 | 0.850 | 1.000 | 0.226 | 0.002 |
| OFd | OFm | 17642 | 20372 | 0.866 | 0.951 | 1.000 | 0.141 | 0.001 |
| OFd | ON | 11549 | 18122 | 0.637 | 0.839 | 1.000 | 0.238 | 0.002 |
| OFd | SPd | 12855 | 20225 | 0.636 | 0.854 | 1.000 | 0.223 | 0.002 |
| OFd | SPm | 12728 | 20273 | 0.628 | 0.851 | 1.000 | 0.225 | 0.002 |
| OFm | ON | 11764 | 18064 | 0.651 | 0.844 | 1.000 | 0.236 | 0.002 |
| OFm | SPd | 12624 | 20169 | 0.626 | 0.850 | 1.000 | 0.226 | 0.002 |
| OFm | SPm | 12779 | 20206 | 0.632 | 0.853 | 1.000 | 0.224 | 0.002 |
| SPd | ON | 9465 | 18521 | 0.511 | 0.785 | 1.000 | 0.258 | 0.002 |
| SPd | SPm | 19829 | 21197 | 0.935 | 0.975 | 1.000 | 0.107 | 0.001 |
| SPm | ON | 9548 | 18527 | 0.515 | 0.787 | 1.000 | 0.257 | 0.002 |

(B) Diploid set - ARS

| algorithm1 | algorithm2 | num of OG exact same | total num OG | proportion of OG exact same | mean | median | sd | se |
|---|---|---|---|---|---|---|---|---|
| BR | OFb | 11607 | 19471 | 0.596 | 0.621 | 1.000 | 0.491 | 0.004 |
| BR | OFd | 11965 | 19571 | 0.611 | 0.634 | 1.000 | 0.488 | 0.003 |
| BR | OFm | 11797 | 19530 | 0.604 | 0.628 | 1.000 | 0.489 | 0.003 |
| BR | ON | 10309 | 18092 | 0.570 | 0.596 | 1.000 | 0.493 | 0.004 |
| BR | SPd | 13308 | 20248 | 0.657 | 0.692 | 1.000 | 0.467 | 0.003 |
| BR | SPm | 13409 | 20303 | 0.660 | 0.695 | 1.000 | 0.466 | 0.003 |
| OFb | OFd | 16563 | 20208 | 0.820 | 0.834 | 1.000 | 0.375 | 0.003 |

| | | | | | | | | |
|---|---|---|---|---|---|---|---|---|
| OFb | OFm | 17627 | 20273 | 0.869 | 0.879 | 1.000 | 0.327 | 0.002 |
| OFb | ON | 11579 | 18026 | 0.642 | 0.658 | 1.000 | 0.476 | 0.004 |
| OFb | SPd | 12494 | 20093 | 0.622 | 0.664 | 1.000 | 0.475 | 0.003 |
| OFb | SPm | 12572 | 20131 | 0.625 | 0.666 | 1.000 | 0.474 | 0.003 |
| OFd | OFm | 17642 | 20372 | 0.866 | 0.877 | 1.000 | 0.330 | 0.002 |
| OFd | ON | 11549 | 18122 | 0.637 | 0.656 | 1.000 | 0.477 | 0.004 |
| OFd | SPd | 12855 | 20225 | 0.636 | 0.676 | 1.000 | 0.471 | 0.003 |
| OFd | SPm | 12728 | 20273 | 0.628 | 0.668 | 1.000 | 0.474 | 0.003 |
| OFm | ON | 11764 | 18064 | 0.651 | 0.666 | 1.000 | 0.474 | 0.004 |
| OFm | SPd | 12624 | 20169 | 0.626 | 0.667 | 1.000 | 0.474 | 0.003 |
| OFm | SPm | 12779 | 20206 | 0.632 | 0.673 | 1.000 | 0.471 | 0.003 |
| SPd | ON | 9465 | 18521 | 0.511 | 0.555 | 1.000 | 0.498 | 0.004 |
| SPd | SPm | 19829 | 21197 | 0.935 | 0.945 | 1.000 | 0.230 | 0.002 |
| SPm | ON | 9548 | 18527 | 0.515 | 0.558 | 1.000 | 0.498 | 0.004 |

(C) Diploid set - Jaccard Index (JI)

| algorithm1 | algorithm2 | num of OG exact same | total num OG | proportion of OG exact same | mean | median | sd | se |
|---|---|---|---|---|---|---|---|---|
| BR | OFb | 11607 | 19471 | 0.596 | 0.846 | 1.000 | 0.236 | 0.002 |
| BR | OFd | 11965 | 19571 | 0.611 | 0.853 | 1.000 | 0.233 | 0.002 |
| BR | OFm | 11797 | 19530 | 0.604 | 0.851 | 1.000 | 0.233 | 0.002 |
| BR | ON | 10309 | 18092 | 0.570 | 0.829 | 1.000 | 0.244 | 0.002 |
| BR | SPd | 13308 | 20248 | 0.657 | 0.861 | 1.000 | 0.242 | 0.002 |
| BR | SPm | 13409 | 20303 | 0.660 | 0.863 | 1.000 | 0.240 | 0.002 |
| OFb | OFd | 16563 | 20208 | 0.820 | 0.940 | 1.000 | 0.160 | 0.001 |
| OFb | OFm | 17627 | 20273 | 0.869 | 0.958 | 1.000 | 0.134 | 0.001 |
| OFb | ON | 11579 | 18026 | 0.642 | 0.859 | 1.000 | 0.229 | 0.002 |
| OFb | SPd | 12494 | 20093 | 0.622 | 0.853 | 1.000 | 0.239 | 0.002 |
| OFb | SPm | 12572 | 20131 | 0.625 | 0.855 | 1.000 | 0.238 | 0.002 |
| OFd | OFm | 17642 | 20372 | 0.866 | 0.956 | 1.000 | 0.139 | 0.001 |
| OFd | ON | 11549 | 18122 | 0.637 | 0.856 | 1.000 | 0.232 | 0.002 |
| OFd | SPd | 12855 | 20225 | 0.636 | 0.859 | 1.000 | 0.236 | 0.002 |
| OFd | SPm | 12728 | 20273 | 0.628 | 0.857 | 1.000 | 0.238 | 0.002 |
| OFm | ON | 11764 | 18064 | 0.651 | 0.862 | 1.000 | 0.227 | 0.002 |
| OFm | SPd | 12624 | 20169 | 0.626 | 0.855 | 1.000 | 0.239 | 0.002 |
| OFm | SPm | 12779 | 20206 | 0.632 | 0.859 | 1.000 | 0.235 | 0.002 |
| SPd | ON | 9465 | 18521 | 0.511 | 0.793 | 1.000 | 0.268 | 0.002 |
| SPd | SPm | 19829 | 21197 | 0.935 | 0.975 | 1.000 | 0.115 | 0.001 |
| SPm | ON | 9548 | 18527 | 0.515 | 0.797 | 1.000 | 0.265 | 0.002 |

(D) Diploid+higher ploidy set - RS

| algorithm1 | algorithm2 | num of OG exact same | total num OG | proportion of OG exact same | mean | median | sd | se |
|---|---|---|---|---|---|---|---|---|
| BR | OFb | 7064 | 20034 | 0.353 | 0.780 | 0.818 | 0.210 | 0.001 |
| BR | OFd | 6765 | 20316 | 0.333 | 0.774 | 0.800 | 0.207 | 0.001 |
| BR | OFm | 7213 | 20178 | 0.357 | 0.783 | 0.818 | 0.209 | 0.001 |
| BR | ON | 8232 | 18828 | 0.437 | 0.806 | 0.846 | 0.221 | 0.002 |
| BR | SPd | 7397 | 20520 | 0.360 | 0.788 | 0.818 | 0.206 | 0.001 |
| BR | SPm | 7479 | 20552 | 0.364 | 0.789 | 0.818 | 0.207 | 0.001 |
| OFb | OFd | 11391 | 20506 | 0.555 | 0.877 | 1.000 | 0.168 | 0.001 |
| OFb | OFm | 14406 | 20568 | 0.700 | 0.923 | 1.000 | 0.141 | 0.001 |
| OFb | ON | 7256 | 18621 | 0.390 | 0.790 | 0.833 | 0.216 | 0.002 |
| OFb | SPd | 7428 | 19976 | 0.372 | 0.798 | 0.818 | 0.203 | 0.001 |
| OFb | SPm | 7526 | 19997 | 0.376 | 0.800 | 0.818 | 0.203 | 0.001 |
| OFd | OFm | 12936 | 20692 | 0.625 | 0.901 | 1.000 | 0.154 | 0.001 |
| OFd | ON | 6457 | 18808 | 0.343 | 0.770 | 0.809 | 0.217 | 0.002 |
| OFd | SPd | 8096 | 20361 | 0.398 | 0.812 | 0.833 | 0.198 | 0.001 |
| OFd | SPm | 7925 | 20349 | 0.389 | 0.808 | 0.818 | 0.199 | 0.001 |
| OFm | ON | 7470 | 18704 | 0.399 | 0.793 | 0.833 | 0.216 | 0.002 |
| OFm | SPd | 7575 | 20181 | 0.375 | 0.802 | 0.818 | 0.201 | 0.001 |
| OFm | SPm | 7728 | 20188 | 0.383 | 0.805 | 0.818 | 0.200 | 0.001 |
| SPd | ON | 5402 | 18773 | 0.288 | 0.749 | 0.800 | 0.218 | 0.002 |
| SPd | SPm | 18048 | 21043 | 0.858 | 0.961 | 1.000 | 0.115 | 0.001 |
| SPm | ON | 5434 | 18772 | 0.289 | 0.750 | 0.800 | 0.218 | 0.002 |

(E) Diploid+higher ploidy set - ARS

| algorithm1 | algorithm2 | num of OG exact same | total num OG | proportion of OG exact same | mean | median | sd | se |
|---|---|---|---|---|---|---|---|---|
| BR | OFb | 7064 | 20034 | 0.353 | 0.353 | 0.000 | 0.492 | 0.003 |
| BR | OFd | 6765 | 20316 | 0.333 | 0.331 | 0.000 | 0.485 | 0.003 |
| BR | OFm | 7213 | 20178 | 0.357 | 0.357 | 0.000 | 0.493 | 0.003 |
| BR | ON | 8232 | 18828 | 0.437 | 0.447 | 0.000 | 0.502 | 0.004 |
| BR | SPd | 7397 | 20520 | 0.360 | 0.372 | 0.000 | 0.492 | 0.003 |
| BR | SPm | 7479 | 20552 | 0.364 | 0.376 | 0.000 | 0.492 | 0.003 |
| OFb | OFd | 11391 | 20506 | 0.555 | 0.561 | 1.000 | 0.501 | 0.003 |
| OFb | OFm | 14406 | 20568 | 0.700 | 0.706 | 1.000 | 0.458 | 0.003 |
| OFb | ON | 7256 | 18621 | 0.390 | 0.397 | 0.000 | 0.493 | 0.004 |
| OFb | SPd | 7428 | 19976 | 0.372 | 0.384 | 0.000 | 0.497 | 0.004 |
| OFb | SPm | 7526 | 19997 | 0.376 | 0.389 | 0.000 | 0.498 | 0.004 |
| OFd | OFm | 12936 | 20692 | 0.625 | 0.630 | 1.000 | 0.485 | 0.003 |

| | | | | | | | | |
|---|---|---|---|---|---|---|---|---|
| OFd | ON | 6457 | 18808 | 0.343 | 0.349 | 0.000 | 0.480 | 0.003 |
| OFd | SPd | 8096 | 20361 | 0.398 | 0.411 | 0.000 | 0.503 | 0.004 |
| OFd | SPm | 7925 | 20349 | 0.389 | 0.402 | 0.000 | 0.502 | 0.004 |
| OFm | ON | 7470 | 18704 | 0.399 | 0.406 | 0.000 | 0.494 | 0.004 |
| OFm | SPd | 7575 | 20181 | 0.375 | 0.387 | 0.000 | 0.498 | 0.004 |
| OFm | SPm | 7728 | 20188 | 0.383 | 0.395 | 0.000 | 0.499 | 0.004 |
| SPd | ON | 5402 | 18773 | 0.288 | 0.306 | 0.000 | 0.463 | 0.003 |
| SPd | SPm | 18048 | 21043 | 0.858 | 0.865 | 1.000 | 0.343 | 0.002 |
| SPm | ON | 5434 | 18772 | 0.289 | 0.308 | 0.000 | 0.464 | 0.003 |

(F) Diploid+higher ploidy set - Jaccard Index

| algorithm1 | algorithm2 | num of OG exact same | total num OG | proportion of OG exact same | mean | median | sd | se |
|---|---|---|---|---|---|---|---|---|
| BR | OFb | 7064 | 20034 | 0.353 | 0.823 | 0.900 | 0.219 | 0.002 |
| BR | OFd | 6765 | 20316 | 0.333 | 0.824 | 0.900 | 0.212 | 0.001 |
| BR | OFm | 7213 | 20178 | 0.357 | 0.828 | 0.900 | 0.214 | 0.002 |
| BR | ON | 8232 | 18828 | 0.437 | 0.839 | 0.923 | 0.220 | 0.002 |
| BR | SPd | 7397 | 20520 | 0.360 | 0.823 | 0.900 | 0.226 | 0.002 |
| BR | SPm | 7479 | 20552 | 0.364 | 0.824 | 0.900 | 0.225 | 0.002 |
| OFb | OFd | 11391 | 20506 | 0.555 | 0.908 | 1.000 | 0.164 | 0.001 |
| OFb | OFm | 14406 | 20568 | 0.700 | 0.942 | 1.000 | 0.136 | 0.001 |
| OFb | ON | 7256 | 18621 | 0.390 | 0.832 | 0.909 | 0.214 | 0.002 |
| OFb | SPd | 7428 | 19976 | 0.372 | 0.826 | 0.909 | 0.227 | 0.002 |
| OFb | SPm | 7526 | 19997 | 0.376 | 0.827 | 0.909 | 0.226 | 0.002 |
| OFd | OFm | 12936 | 20692 | 0.625 | 0.927 | 1.000 | 0.146 | 0.001 |
| OFd | ON | 6457 | 18808 | 0.343 | 0.821 | 0.900 | 0.214 | 0.002 |
| OFd | SPd | 8096 | 20361 | 0.398 | 0.838 | 0.909 | 0.223 | 0.002 |
| OFd | SPm | 7925 | 20349 | 0.389 | 0.835 | 0.909 | 0.224 | 0.002 |
| OFm | ON | 7470 | 18704 | 0.399 | 0.837 | 0.917 | 0.210 | 0.002 |
| OFm | SPd | 7575 | 20181 | 0.375 | 0.830 | 0.909 | 0.224 | 0.002 |
| OFm | SPm | 7728 | 20188 | 0.383 | 0.833 | 0.909 | 0.222 | 0.002 |
| SPd | ON | 5402 | 18773 | 0.288 | 0.783 | 0.867 | 0.240 | 0.002 |
| SPd | SPm | 18048 | 21043 | 0.858 | 0.965 | 1.000 | 0.123 | 0.001 |
| SPm | ON | 5434 | 18772 | 0.289 | 0.785 | 0.867 | 0.238 | 0.002 |

Appendix S11-Fig. S6: Distribution of pairwise comparisons between orthology inference algorithms. The Jaccard Index (top row) and Rand Score (bottom row) were calculated for all algorithms in a pairwise manner. (A) Diploid set. (B) Diploid+higher ploidy set. BR: Broccoli. OFb: OrthoFinder-BLAST. OFd: OrthoFinder-DIAMOND. OFm: OrthoFinder-MMseqs2. SPd: SonicParanoid-DIAMOND. SPm: SonicParanoid-MMseqs2. ON: OrthNet. Corresponds to Figure 4, Table S5.

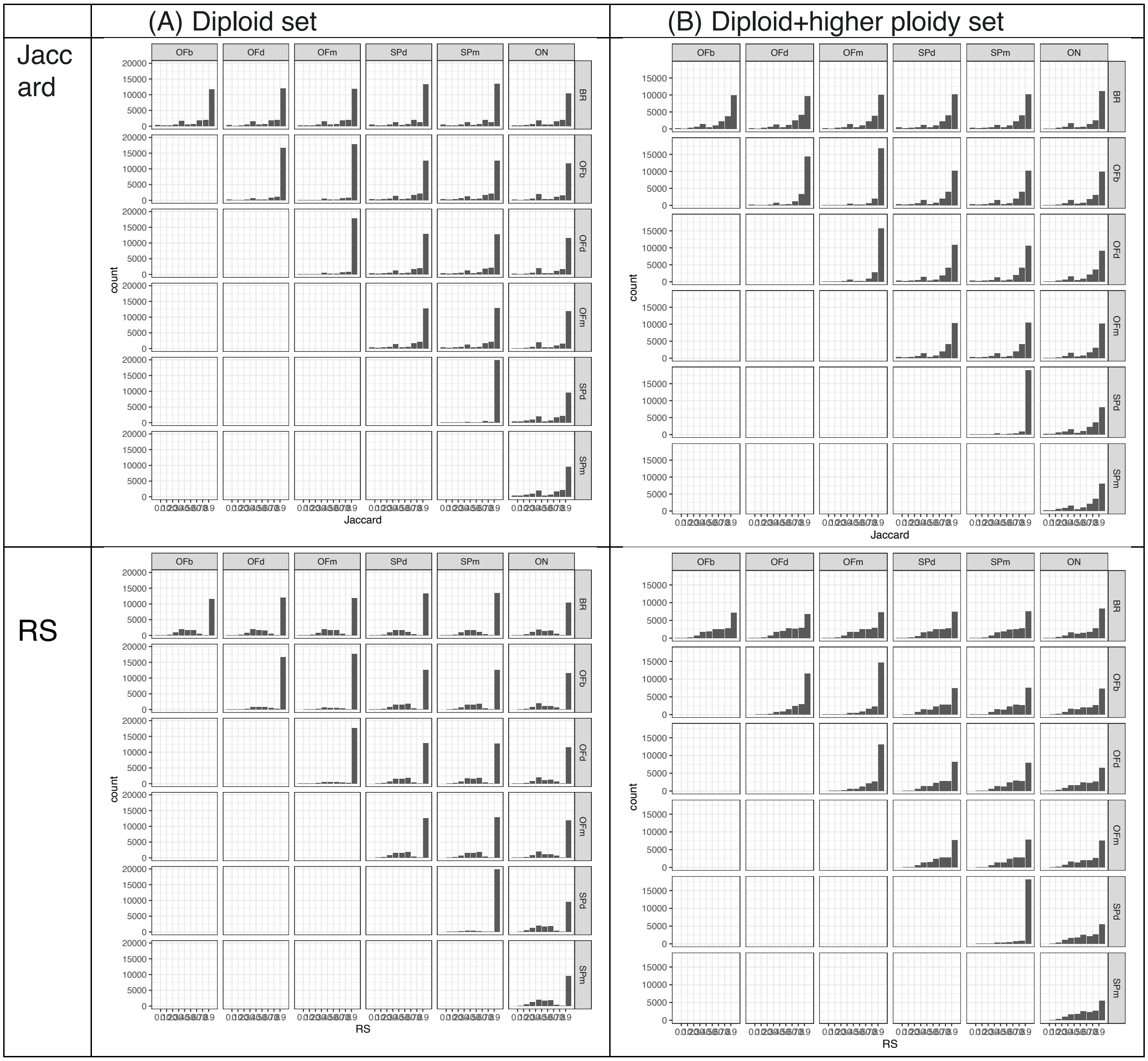

Appendix S12-Fig. S7: Proportion of predicted orthology relationships between all species pairs across algorithms for the diploid set and the diploid+higher ploidy set. Stacked bar plots displaying (A-B) diploid-diploid species pairs from the (A) diploid set and the (B) diploid+higher species set; (C) diploid-mesopolyploid species pair; (D) diploid-hexaploid species pairs; (E) mesopolyploid-hexaploid species pair. Each stacked bar represents the algorithm used, and the colors represent the category of orthology relationships: one:one (one-to-one), 1:M (one to many), M:1 (many to one), M:M (many to many). OFb_base: OrthoFinder-BLAST-MCL, the baseline to compare the results from all other algorithms. Species abbreviations can be found in Table 1. BR: Broccoli. OFb: OrthoFinder-BLAST. OFd: OrthoFinder-DIAMOND. OFm: OrthoFinder-MMseqs2. SPd: SonicParanoid-DIAMOND. SPm: SonicParanoid-MMseqs2. ON: OrthNet.

(A) diploid-diploid species pairs (diploid set)

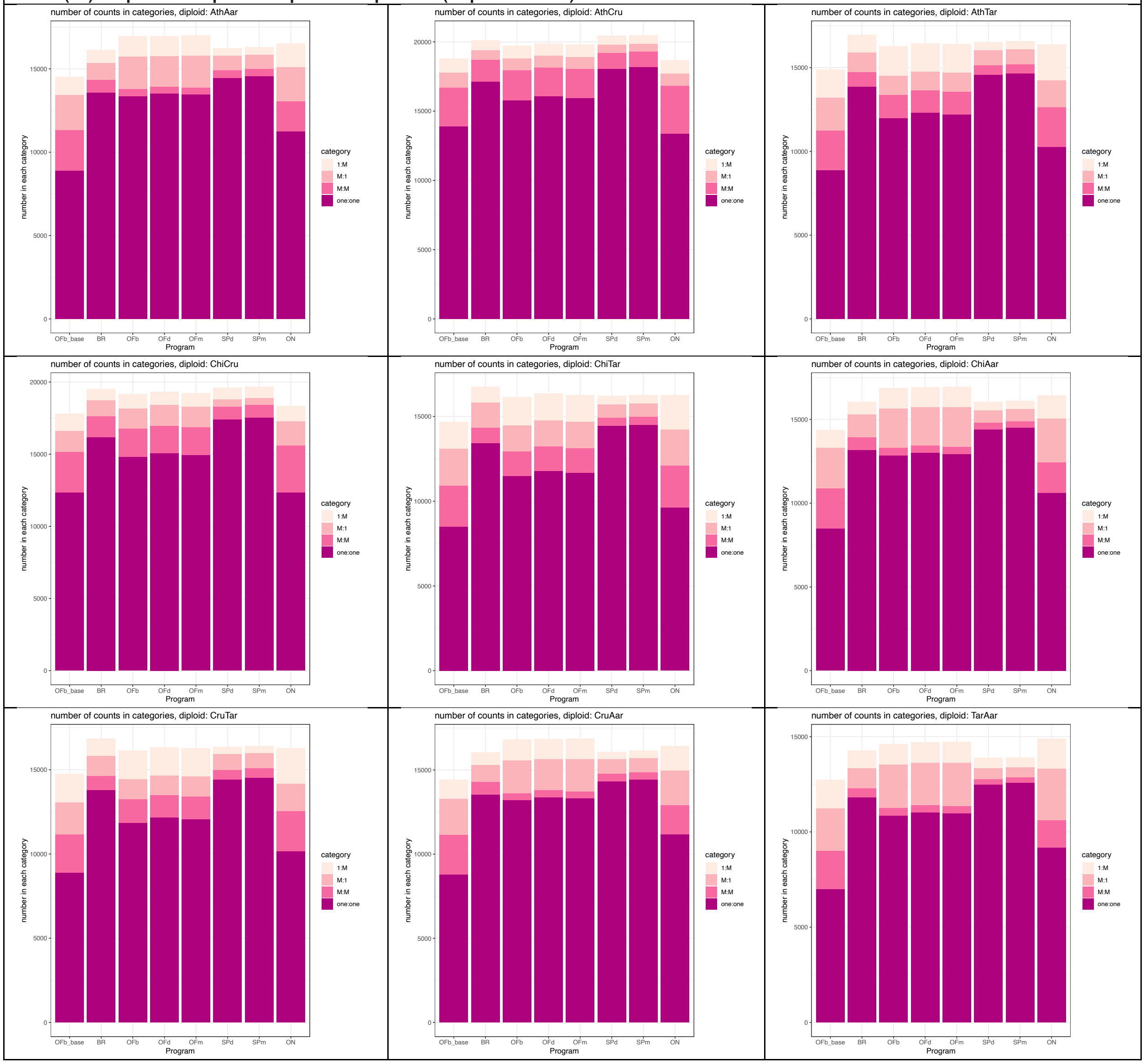

#### (B) diploid-diploid species pairs (diploid+higher ploidy set)

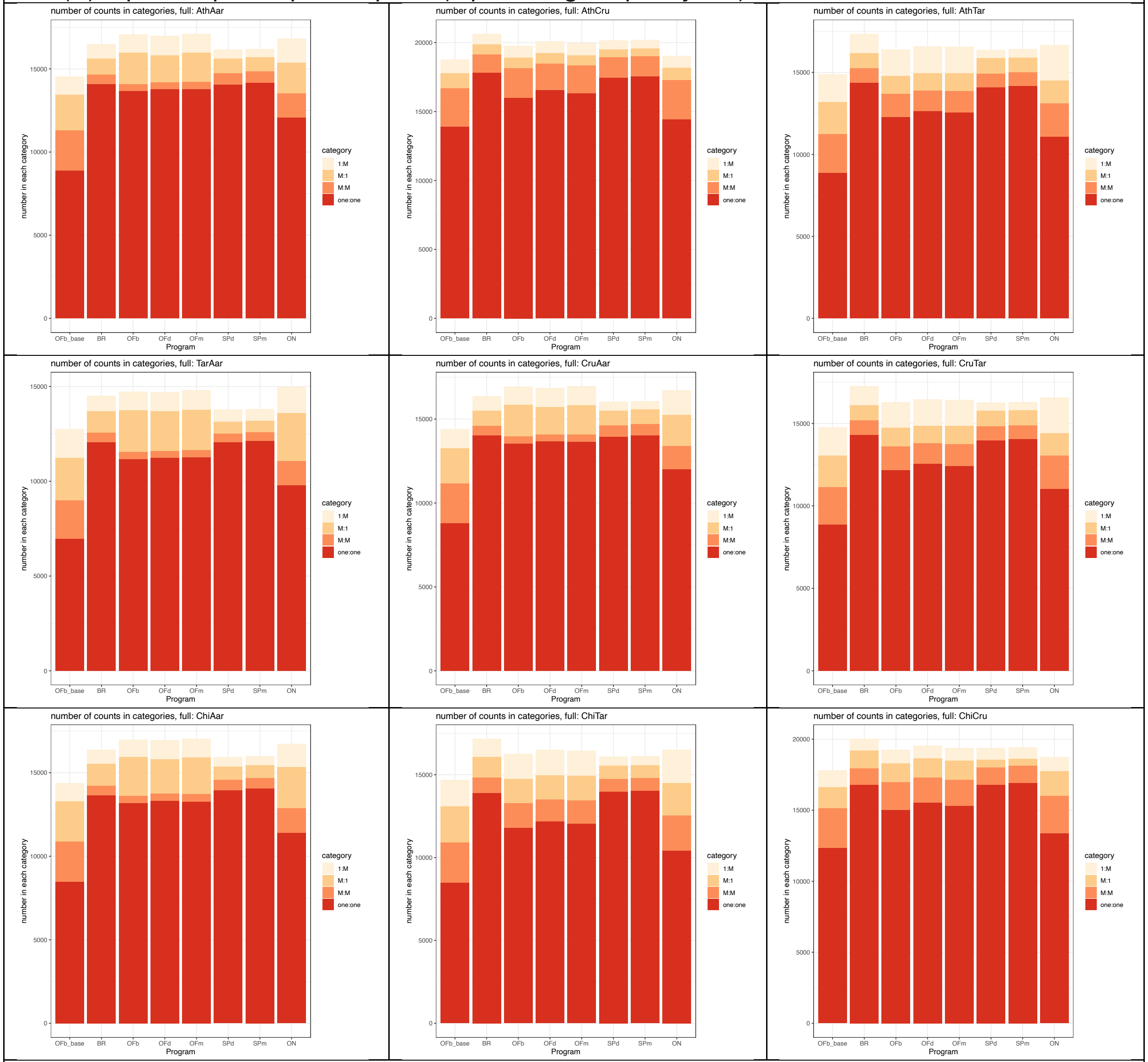

#### (C) diploid-mesopolyploid species pairs

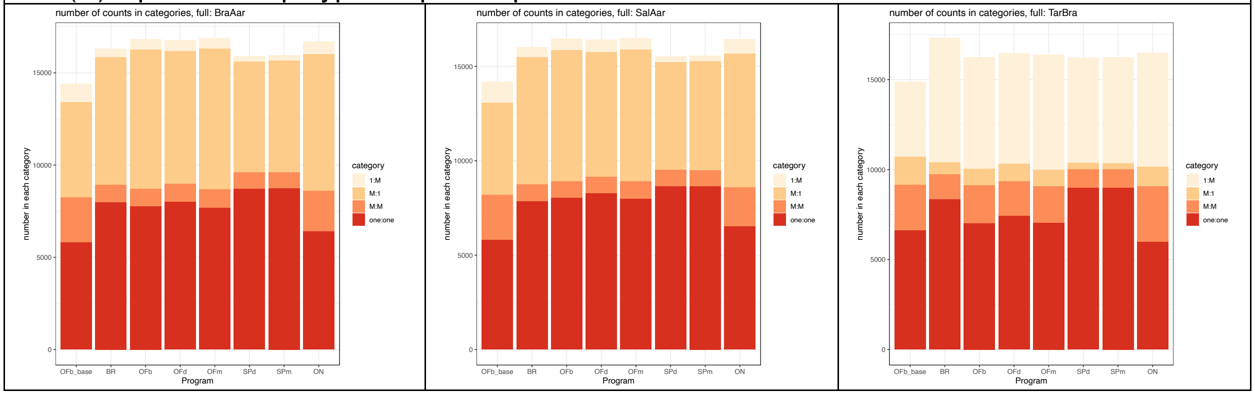

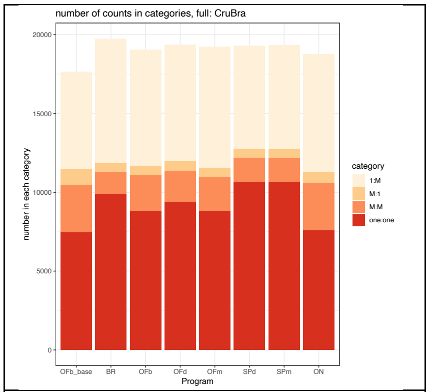

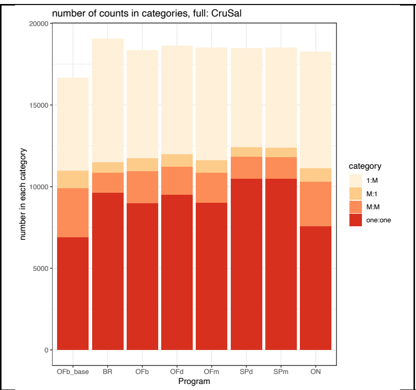

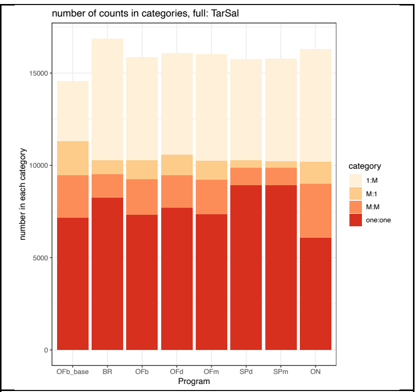

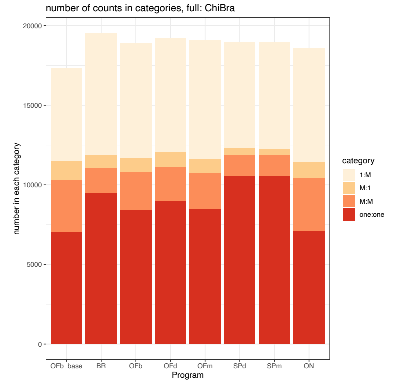

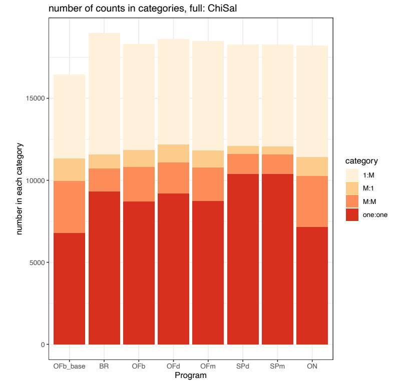

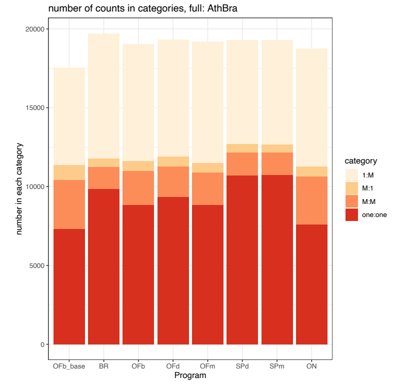

(D) diploid-hexaploid species pair

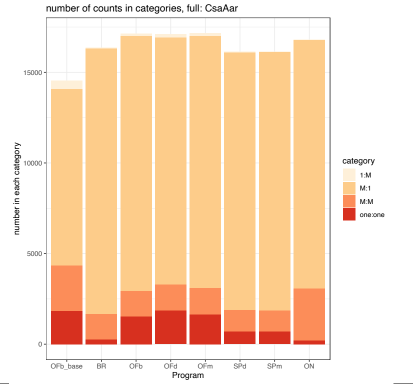

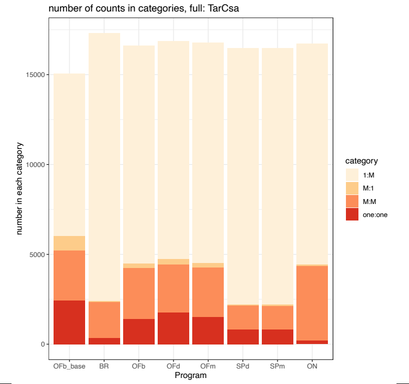

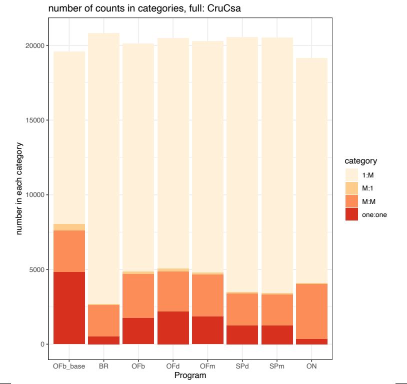

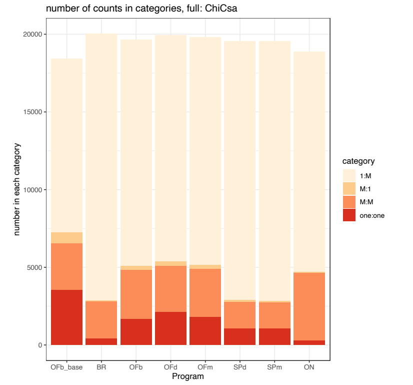

(E) mesopolyploid-hexaploid species pair

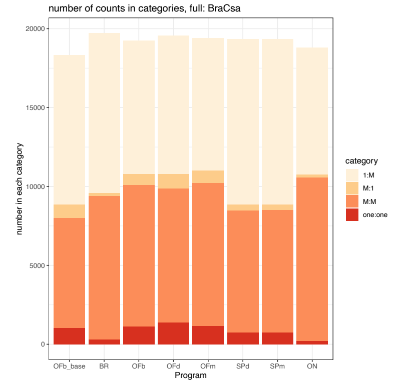

Appendix S13-Table S6: Species pair ortholog ratios across algorithms. Ratios consist of four categori
– one-to-many orthologs, M:1 – many-to-one orthologs, M:M – many-to-many orthologs. Each row repr
the shorthand found in Table 1, in which the order of the species is the same as the ratio order. All sev
algorithms and the baseline algorithm (OrthoFinder-BLAST-MCL) were tested. (A) Diploid set. (B) Dipl
Corresponds to Fig. 5 and Appendix S13.

| SpPair | baseline: Orthofinder-BLAST -MCL ONLY | | | | diploid broccoli | | | | diploid OI | |
|---|---|---|---|---|---|---|---|---|---|---|
| | one:one | 1:M | M:1 | M:M | one:one | 1:M | M:1 | M:M | one:one | 1:M |
| **AthChi** | 12389 | 1412 | 1140 | 3013 | 16491 | 1210 | 780 | 1061 | 16016 | 1507 |
| **AthCru** | 13899 | 1025 | 1084 | 2784 | 17494 | 817 | 707 | 1167 | 16954 | 926 |
| **AthTar** | 8885 | 1712 | 1946 | 2357 | 13835 | 1071 | 945 | 656 | 13376 | 1964 |
| **ChiCru** | 12359 | 1173 | 1471 | 2792 | 16227 | 832 | 1176 | 1037 | 15815 | 984 |
| **ChiTar** | 8500 | 1614 | 2157 | 2423 | 13298 | 1015 | 1307 | 676 | 12775 | 1912 |
| **CruTar** | 8877 | 1701 | 1911 | 2268 | 13709 | 1037 | 984 | 676 | 13271 | 1938 |

| SpPair | baseline: Orthofinder-BLAST -MCL ONLY | | | | diploid broccoli | | | | diploid OI | |
|---|---|---|---|---|---|---|---|---|---|---|
| | one:one | 1:M | M:1 | M:M | one:one | 1:M | M:1 | M:M | one:one | 1:M |
| AthChi | 12389 | 1412 | 1140 | 3013 | 16340 | 1136 | 752 | 1489 | 15032 | 1429 |
| AthCru | 13899 | 1025 | 1084 | 2784 | 17138 | 697 | 687 | 1576 | 15789 | 917 |
| AthTar | 8885 | 1712 | 1946 | 2357 | 13881 | 1048 | 1166 | 855 | 11979 | 1749 |
| AthAar | 8890 | 1109 | 2120 | 2416 | 13579 | 784 | 1034 | 747 | 13356 | 1244 |
| ChiCru | 12359 | 1173 | 1471 | 2792 | 16171 | 770 | 1131 | 1429 | 14801 | 993 |
| ChiTar | 8500 | 1614 | 2157 | 2423 | 13419 | 944 | 1480 | 907 | 11490 | 1647 |
| ChiAar | 8495 | 1075 | 2406 | 2385 | 13180 | 748 | 1359 | 756 | 12849 | 1223 |
| CruTar | 8877 | 1701 | 1911 | 2268 | 13790 | 1038 | 1183 | 839 | 11856 | 1700 |
| CruAar | 8789 | 1144 | 2121 | 2359 | 13542 | 763 | 989 | 745 | 13217 | 1234 |
| TarAar | 6988 | 1515 | 2233 | 2007 | 11813 | 948 | 1043 | 473 | 10850 | 1079 |

| F blast | | diploid OF diamond | | | | diploid OF mmseqs | | | | diploid OrthNet | | | |
|---|---|---|---|---|---|---|---|---|---|---|---|---|---|
| M:1 | M:M | one:one | 1:M | M:1 | M:M | one:one | 1:M | M:1 | M:M | one:one | 1:M | M:1 | M:M |
| 945 | 1461 | 16344 | 1517 | 887 | 1383 | 16130 | 1542 | 937 | 1449 | 13937 | 1930 | 1055 | 2587 |
| 889 | 1611 | 17272 | 916 | 872 | 1486 | 17064 | 932 | 887 | 1575 | 14985 | 1032 | 976 | 2700 |
| 1633 | 422 | 13691 | 1922 | 1544 | 397 | 13588 | 1940 | 1621 | 411 | 11456 | 2370 | 1610 | 1761 |
| 1492 | 1402 | 16137 | 933 | 1480 | 1315 | 15917 | 980 | 1507 | 1384 | 13762 | 1123 | 1916 | 2501 |
| 2116 | 472 | 13109 | 1858 | 2083 | 465 | 12978 | 1890 | 2130 | 472 | 10736 | 2189 | 2206 | 1905 |
| 1659 | 410 | 13591 | 1867 | 1568 | 404 | 13479 | 1890 | 1630 | 414 | 11378 | 2323 | 1595 | 1771 |

| F blast | | diploid OF diamond | | | | diploid OF mmseqs | | | | diploid OrthNet | | | |
|---|---|---|---|---|---|---|---|---|---|---|---|---|---|
| M:1 | M:M | one:one | 1:M | M:1 | M:M | one:one | 1:M | M:1 | M:M | one:one | 1:M | M:1 | M:M |
| 937 | 2013 | 15286 | 1449 | 883 | 1950 | 15165 | 1459 | 903 | 1976 | 12508 | 1711 | 1009 | 3309 |
| 839 | 2173 | 16084 | 902 | 824 | 2061 | 15950 | 933 | 837 | 2098 | 13383 | 950 | 906 | 3429 |
| 1167 | 1380 | 12302 | 1688 | 1120 | 1335 | 12195 | 1741 | 1150 | 1348 | 10258 | 2162 | 1607 | 2368 |
| 1952 | 414 | 13522 | 1221 | 1821 | 410 | 13479 | 1255 | 1890 | 407 | 11263 | 1428 | 2055 | 1787 |
| 1414 | 1957 | 15073 | 931 | 1425 | 1891 | 14925 | 961 | 1433 | 1928 | 12357 | 1053 | 1703 | 3233 |
| 1533 | 1452 | 11786 | 1585 | 1543 | 1430 | 11671 | 1613 | 1554 | 1449 | 9639 | 2033 | 2118 | 2470 |
| 2364 | 437 | 13002 | 1205 | 2300 | 430 | 12921 | 1226 | 2379 | 424 | 10610 | 1385 | 2626 | 1820 |
| 1190 | 1385 | 12160 | 1663 | 1170 | 1326 | 12047 | 1679 | 1194 | 1354 | 10177 | 2137 | 1613 | 2362 |
| 1948 | 405 | 13393 | 1204 | 1826 | 407 | 13322 | 1235 | 1909 | 399 | 11164 | 1456 | 2067 | 1732 |
| 2295 | 386 | 11032 | 1088 | 2217 | 365 | 10977 | 1084 | 2280 | 372 | 9178 | 1578 | 2705 | 1436 |

| diploid SP diamond | | | | diploid SP mmseqs | | | |
|---|---|---|---|---|---|---|---|
| one:one | 1:M | M:1 | M:M | one:one | 1:M | M:1 | M:M |
| 18637 | 458 | 671 | 529 | 18715 | 426 | 646 | 534 |
| 19133 | 584 | 518 | 725 | 19172 | 579 | 520 | 711 |
| 15260 | 403 | 712 | 326 | 15307 | 385 | 711 | 322 |
| 18357 | 698 | 446 | 495 | 18447 | 676 | 405 | 503 |
| 15077 | 431 | 576 | 268 | 15135 | 413 | 559 | 267 |
| 15111 | 387 | 708 | 339 | 15161 | 372 | 693 | 336 |

| diploid SP diamond | | | | diploid SP mmseqs | | | |
|---|---|---|---|---|---|---|---|
| one:one | 1:M | M:1 | M:M | one:one | 1:M | M:1 | M:M |
| 17688 | 530 | 762 | 919 | 17817 | 513 | 743 | 889 |
| 18065 | 658 | 590 | 1131 | 18177 | 647 | 577 | 1091 |
| 14571 | 478 | 916 | 559 | 14656 | 477 | 902 | 541 |
| 14447 | 470 | 883 | 445 | 14563 | 459 | 856 | 426 |
| 17393 | 807 | 517 | 885 | 17546 | 779 | 486 | 860 |
| 14435 | 519 | 785 | 483 | 14515 | 506 | 766 | 477 |
| 14394 | 510 | 745 | 396 | 14502 | 491 | 725 | 381 |
| 14413 | 459 | 936 | 568 | 14512 | 439 | 897 | 563 |
| 14320 | 455 | 860 | 452 | 14427 | 457 | 846 | 420 |
| 12499 | 568 | 552 | 272 | 12575 | 539 | 532 | 276 |

| | baseline: Orthofinder-BLAST -MCL ONLY | | | | full broccoli | | | | full OF l | |
|---|---|---|---|---|---|---|---|---|---|---|
| SpPair | one:one | 1:M | M:1 | M:M | one:one | 1:M | M:1 | M:M | one:one | 1:M |
| **AthChi** | 12389 | 1412 | 1140 | 3013 | 16959 | 1252 | 808 | 1191 | 15229 | 1366 |
| **AthCru** | 13899 | 1025 | 1084 | 2784 | 17812 | 779 | 736 | 1305 | 15992 | 870 |
| **AthTar** | 8885 | 1712 | 1946 | 2357 | 14392 | 1186 | 909 | 866 | 12276 | 1603 |
| **AthSal** | 6828 | 5828 | 1056 | 3077 | 9685 | 7577 | 624 | 1220 | 9074 | 6675 |
| **AthBra** | 7321 | 6168 | 957 | 3094 | 9862 | 7924 | 567 | 1365 | 8832 | 7430 |
| **AthCsa** | 3572 | 12222 | 409 | 2793 | 497 | 18081 | 77 | 1987 | 1742 | 15288 |
| **AthAar** | 8890 | 1109 | 2120 | 2416 | 14107 | 885 | 948 | 552 | 13676 | 1089 |
| **ChiCru** | 12359 | 1173 | 1471 | 2792 | 16797 | 841 | 1245 | 1142 | 15020 | 955 |
| **ChiTar** | 8500 | 1614 | 2157 | 2423 | 13903 | 1089 | 1262 | 914 | 11811 | 1511 |
| **ChiSal** | 6804 | 5120 | 1349 | 3162 | 9323 | 7389 | 873 | 1374 | 8706 | 6467 |
| **ChiBra** | 7054 | 5852 | 1208 | 3205 | 9496 | 7692 | 803 | 1540 | 8458 | 7192 |
| **ChiCsa** | 3554 | 11159 | 726 | 2969 | 438 | 17161 | 88 | 2353 | 1680 | 14535 |
| **ChiAar** | 8495 | 1075 | 2406 | 2385 | 13647 | 865 | 1320 | 549 | 13188 | 1060 |
| **CruTar** | 8877 | 1701 | 1911 | 2268 | 14307 | 1143 | 915 | 877 | 12166 | 1555 |
| **CruSal** | 6892 | 5729 | 1070 | 2998 | 9638 | 7570 | 641 | 1217 | 9005 | 6631 |
| **CruBra** | 7472 | 6202 | 970 | 3004 | 9874 | 7912 | 571 | 1397 | 8836 | 7369 |
| **CruCsa** | 4851 | 11538 | 431 | 2756 | 526 | 18132 | 75 | 2079 | 1773 | 15262 |
| **CruAar** | 8789 | 1144 | 2121 | 2359 | 14038 | 871 | 911 | 548 | 13546 | 1069 |
| **TarSal** | 7154 | 3254 | 1851 | 2291 | 8251 | 6609 | 746 | 1261 | 7329 | 5590 |
| **TarBra** | 6634 | 4167 | 1572 | 2513 | 8343 | 6893 | 684 | 1380 | 7039 | 6190 |
| **TarCsa** | 2424 | 9033 | 829 | 2772 | 336 | 14903 | 64 | 2002 | 1397 | 12137 |
| **TarAar** | 6988 | 1515 | 2233 | 2007 | 12053 | 824 | 1148 | 492 | 11181 | 976 |
| **SalBra** | 9238 | 2766 | 1745 | 6594 | 9655 | 1704 | 1382 | 7684 | 8278 | 2112 |
| **SalCsa** | 765 | 9549 | 701 | 6274 | 287 | 10027 | 151 | 8627 | 1073 | 8908 |
| **SalAar** | 5807 | 1110 | 4886 | 2387 | 7871 | 516 | 6751 | 878 | 8060 | 617 |
| **BraCsa** | 1026 | 9446 | 872 | 6962 | 317 | 10122 | 189 | 9080 | 1145 | 8448 |
| **BraAar** | 5804 | 992 | 5168 | 2437 | 7984 | 471 | 6899 | 948 | 7767 | 569 |
| **CsaAar** | 1834 | 448 | 9760 | 2489 | 264 | 35 | 14661 | 1396 | 1549 | 153 |

| blast | | full OF diamond | | | | full OF mmseqs | | | | full OrthNet | | |
|---|---|---|---|---|---|---|---|---|---|---|---|---|
| M:1 | M:M | one:one | 1:M | M:1 | M:M | one:one | 1:M | M:1 | M:M | one:on | 1:M | M:1 |
| 877 | 1991 | 15738 | 1369 | 855 | 1797 | 15515 | 1405 | 855 | 1860 | 13512 | 1698 | 963 |
| 766 | 2140 | 16550 | 851 | 785 | 1910 | 16327 | 863 | 756 | 2006 | 14451 | 882 | 888 |
| 1111 | 1401 | 12660 | 1651 | 1049 | 1231 | 12562 | 1633 | 1066 | 1309 | 11085 | 2152 | 1394 |
| 731 | 1930 | 9582 | 6664 | 757 | 1692 | 9104 | 6939 | 718 | 1806 | 7615 | 7150 | 777 |
| 620 | 2153 | 9357 | 7411 | 628 | 1924 | 8849 | 7734 | 597 | 2036 | 7598 | 7500 | 639 |
| 144 | 2790 | 2165 | 15424 | 186 | 2549 | 1844 | 15526 | 154 | 2633 | 342 | 15065 | 68 |
| 1896 | 411 | 13784 | 1171 | 1649 | 392 | 13796 | 1143 | 1743 | 421 | 12092 | 1450 | 1843 |
| 1343 | 1938 | 15557 | 886 | 1354 | 1741 | 15305 | 909 | 1360 | 1826 | 13393 | 998 | 1728 |
| 1467 | 1466 | 12196 | 1538 | 1455 | 1323 | 12063 | 1519 | 1465 | 1403 | 10427 | 2024 | 1943 |
| 1025 | 2097 | 9210 | 6445 | 1081 | 1873 | 8765 | 6679 | 1013 | 2016 | 7161 | 6792 | 1166 |
| 869 | 2358 | 8979 | 7166 | 911 | 2148 | 8471 | 7439 | 879 | 2282 | 7109 | 7144 | 1032 |
| 263 | 3151 | 2134 | 14581 | 292 | 2946 | 1811 | 14689 | 260 | 3072 | 297 | 14163 | 80 |
| 2301 | 430 | 13316 | 1141 | 2076 | 420 | 13280 | 1110 | 2193 | 443 | 11414 | 1391 | 2455 |
| 1139 | 1415 | 12541 | 1606 | 1078 | 1245 | 12419 | 1580 | 1100 | 1325 | 11032 | 2144 | 1382 |
| 778 | 1937 | 9514 | 6635 | 781 | 1695 | 9029 | 6892 | 763 | 1815 | 7583 | 7160 | 800 |
| 612 | 2242 | 9365 | 7381 | 631 | 1983 | 8843 | 7672 | 604 | 2113 | 7605 | 7498 | 637 |
| 157 | 2927 | 2196 | 15444 | 191 | 2662 | 1873 | 15471 | 155 | 2780 | 350 | 15059 | 57 |
| 1894 | 416 | 13684 | 1147 | 1629 | 389 | 13658 | 1120 | 1750 | 422 | 12023 | 1485 | 1834 |
| 1021 | 1920 | 7721 | 5530 | 1095 | 1740 | 7348 | 5784 | 1035 | 1852 | 6097 | 6081 | 1193 |
| 908 | 2093 | 7444 | 6143 | 969 | 1915 | 7042 | 6416 | 910 | 2028 | 5978 | 6361 | 1091 |
| 248 | 2832 | 1757 | 12143 | 309 | 2664 | 1509 | 12266 | 251 | 2752 | 220 | 12295 | 78 |
| 2192 | 376 | 11240 | 1033 | 2076 | 360 | 11257 | 1019 | 2132 | 381 | 9784 | 1412 | 2541 |
| 1298 | 7527 | 8896 | 2203 | 1380 | 7223 | 8326 | 2122 | 1275 | 7700 | 6734 | 1741 | 1339 |
| 695 | 7943 | 1381 | 9194 | 824 | 7552 | 1117 | 8905 | 754 | 8010 | 231 | 8245 | 124 |
| 6929 | 860 | 8291 | 656 | 6604 | 871 | 7995 | 615 | 6972 | 914 | 6524 | 779 | 7085 |
| 691 | 8951 | 1391 | 8777 | 902 | 8488 | 1178 | 8425 | 771 | 9046 | 218 | 8055 | 159 |
| 7557 | 930 | 8019 | 608 | 7200 | 953 | 7679 | 569 | 7634 | 998 | 6424 | 683 | 7384 |
| 14052 | 1383 | 1875 | 185 | 13630 | 1410 | 1635 | 151 | 13929 | 1443 | 196 | 28 | 13724 |

| | full SP diamond | | | | full SP mmseqs | | | |
|---|---|---|---|---|---|---|---|---|
| **M:M** | **one:one** | **1:M** | **M:1** | **M:M** | **one:one** | **1:M** | **M:1** | **M:M** |
| 2746 | 17084 | 562 | 728 | 1255 | 17195 | 523 | 706 | 1234 |
| 2829 | 17464 | 660 | 559 | 1488 | 17560 | 638 | 536 | 1454 |
| 2034 | 14103 | 522 | 935 | 830 | 14186 | 518 | 904 | 817 |
| 2778 | 10540 | 6120 | 543 | 1326 | 10553 | 6194 | 514 | 1294 |
| 3013 | 10710 | 6598 | 514 | 1455 | 10737 | 6647 | 490 | 1425 |
| 3677 | 1170 | 17059 | 99 | 1969 | 1182 | 17103 | 92 | 1921 |
| 1437 | 14074 | 537 | 876 | 674 | 14169 | 505 | 858 | 664 |
| 2625 | 16800 | 814 | 542 | 1220 | 16928 | 789 | 503 | 1192 |
| 2120 | 13974 | 552 | 820 | 756 | 14048 | 540 | 797 | 747 |
| 3089 | 10410 | 6165 | 486 | 1192 | 10409 | 6233 | 465 | 1160 |
| 3302 | 10567 | 6636 | 416 | 1321 | 10587 | 6694 | 403 | 1283 |
| 4326 | 1072 | 16666 | 101 | 1700 | 1086 | 16715 | 90 | 1652 |
| 1474 | 13962 | 572 | 793 | 624 | 14060 | 545 | 773 | 612 |
| 2000 | 13964 | 498 | 955 | 836 | 14045 | 490 | 924 | 830 |
| 2728 | 10480 | 6071 | 571 | 1348 | 10487 | 6157 | 557 | 1315 |
| 3011 | 10686 | 6568 | 550 | 1506 | 10701 | 6631 | 544 | 1463 |
| 3679 | 1255 | 17072 | 110 | 2110 | 1255 | 17111 | 97 | 2061 |
| 1382 | 13939 | 524 | 886 | 677 | 14028 | 506 | 881 | 658 |
| 2908 | 8939 | 5489 | 381 | 936 | 8933 | 5551 | 372 | 923 |
| 3076 | 9001 | 5857 | 338 | 1019 | 9015 | 5896 | 332 | 1006 |
| 4143 | 823 | 14260 | 71 | 1310 | 836 | 14291 | 63 | 1295 |
| 1268 | 12052 | 645 | 634 | 454 | 12120 | 613 | 618 | 455 |
| 8667 | 10416 | 1899 | 1466 | 6069 | 10438 | 1857 | 1449 | 6132 |
| 9817 | 718 | 10458 | 322 | 7130 | 715 | 10434 | 323 | 7176 |
| 2057 | 8670 | 317 | 5696 | 861 | 8674 | 312 | 5771 | 827 |
| 10360 | 755 | 10502 | 349 | 7730 | 753 | 10492 | 352 | 7737 |
| 2185 | 8722 | 308 | 5984 | 890 | 8743 | 299 | 6042 | 863 |
| 2860 | 712 | 44 | 14200 | 1177 | 714 | 38 | 14266 | 1137 |

Appendix S14-Fig. S8: The Jaccard Index was calculated from orthology inference results from each algorithm compared to the baseline orthology inference results (OrthoFinder-BLAST-MCL) for each species pair. Top row: the number of orthogroups that are exactly the same, as calculated with the Jaccard Index; blue gradient represents the proportion of orthogroups that are exactly the same. Middle row: the proportion of orthogroups that are exactly same; the blue gradient reflects the same metric. Bottom row: the mean Jaccard Index between algorithms; the green gradient reflects the same metric. (A) Diploid set. (B) diploid+higher ploidy set. Each row represents a species pair using the three letter code (Table 1), and each column represents an algorithm: BR: Broccoli. OFb: OrthoFinder-BLAST. OFd: OrthoFinder-DIAMOND. OFm: OrthoFinder-MMseqs2. SPd: SonicParanoid-DIAMOND. SPm: SonicParanoid-MMseqs2. ON: OrthNet. Corresponds to Table S7.

| | (A) Diploid set | (B) diploid+higher ploidy set |
|---|---|---|
| Jaccard | | |

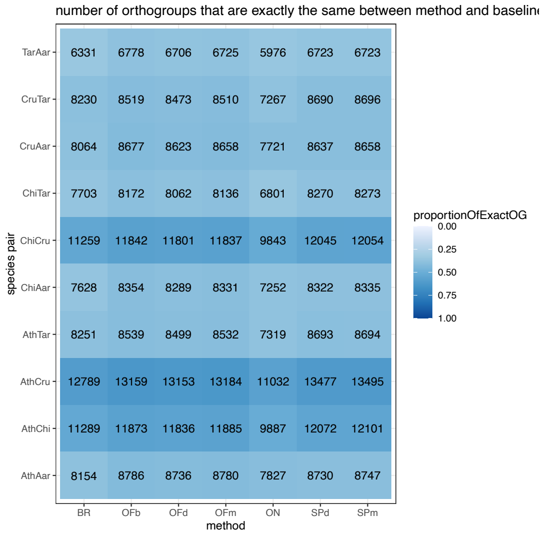

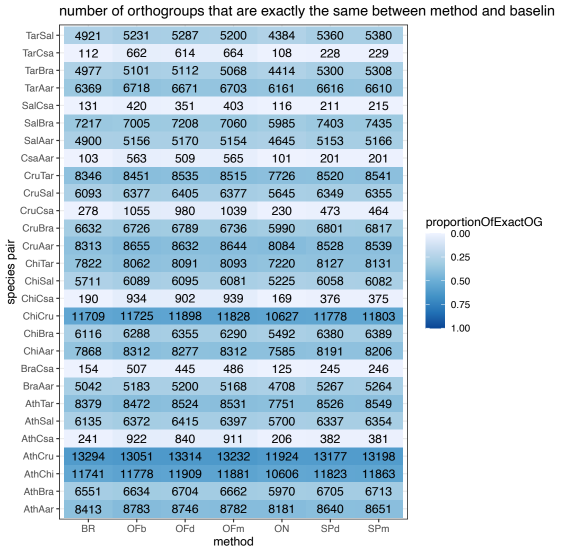

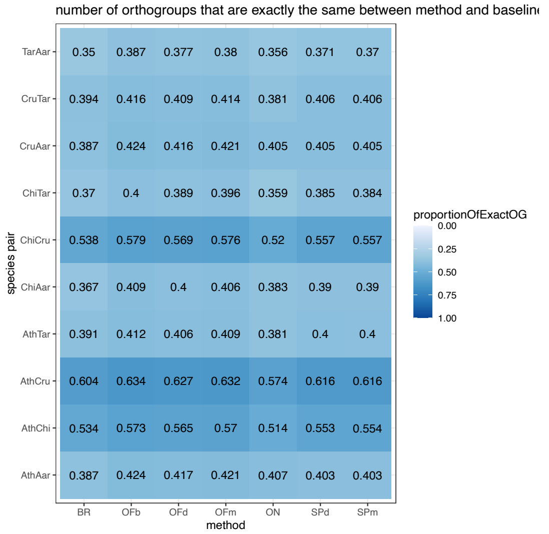

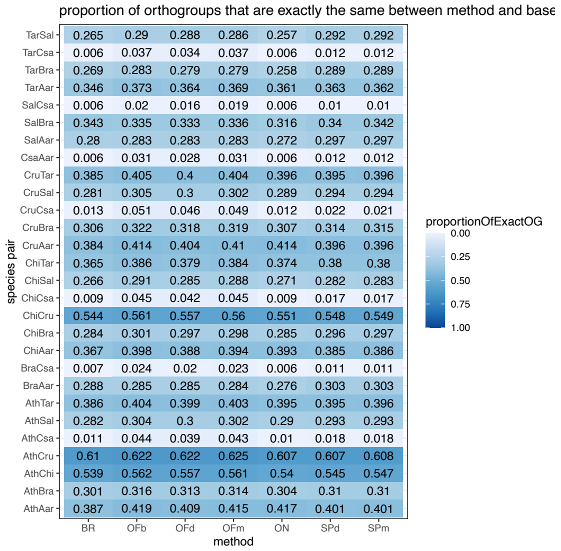

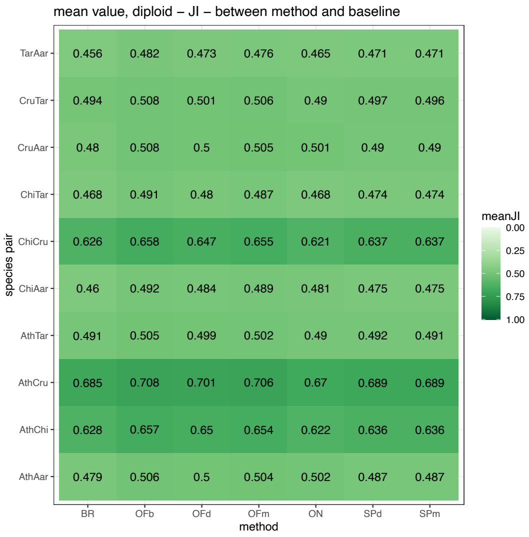
mean value, diploid – JI – between method and baseline
species pair
method
BR OFb OFd OFm ON SPd SPm
TarAar 0.456 0.482 0.473 0.476 0.465 0.471 0.471
CruTar 0.494 0.508 0.501 0.506 0.49 0.497 0.496
CruAar 0.48 0.508 0.5 0.505 0.501 0.49 0.49
ChiTar 0.468 0.491 0.48 0.487 0.468 0.474 0.474
ChiCru 0.626 0.658 0.647 0.655 0.621 0.637 0.637
ChiAar 0.46 0.492 0.484 0.489 0.481 0.475 0.475
AthTar 0.491 0.505 0.499 0.502 0.49 0.492 0.491
AthCru 0.685 0.708 0.701 0.706 0.67 0.689 0.689
AthChi 0.628 0.657 0.65 0.654 0.622 0.636 0.636
AthAar 0.479 0.506 0.5 0.504 0.502 0.487 0.487
meanJI
0.00
0.25
0.50
0.75
1.00

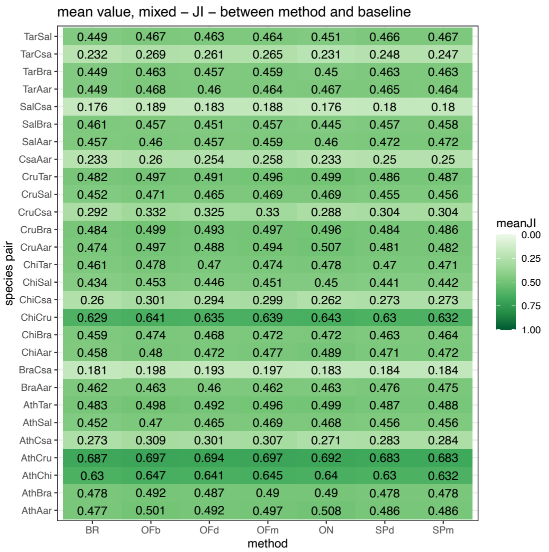
mean value, mixed – JI – between method and baseline
species pair
method
BR OFb OFd OFm ON SPd SPm
TarSal 0.449 0.467 0.463 0.464 0.451 0.466 0.467
TarCsa 0.232 0.269 0.261 0.265 0.231 0.248 0.247
TarBra 0.449 0.463 0.457 0.459 0.45 0.463 0.463
TarAar 0.449 0.468 0.46 0.464 0.467 0.465 0.464
SalCsa 0.176 0.189 0.183 0.188 0.176 0.18 0.18
SalBra 0.461 0.457 0.451 0.457 0.445 0.457 0.458
SalAar 0.457 0.46 0.457 0.459 0.46 0.472 0.472
CsaAar 0.233 0.26 0.254 0.258 0.233 0.25 0.25
CruTar 0.482 0.497 0.491 0.496 0.499 0.486 0.487
CruSal 0.452 0.471 0.465 0.469 0.469 0.455 0.456
CruCsa 0.292 0.332 0.325 0.33 0.288 0.304 0.304
CruBra 0.484 0.499 0.493 0.497 0.496 0.484 0.486
CruAar 0.474 0.497 0.488 0.494 0.507 0.481 0.482
ChiTar 0.461 0.478 0.47 0.474 0.478 0.47 0.471
ChiSal 0.434 0.453 0.446 0.451 0.45 0.441 0.442
ChiCsa 0.26 0.301 0.294 0.299 0.262 0.273 0.273
ChiCru 0.629 0.641 0.635 0.639 0.643 0.63 0.632
ChiBra 0.459 0.474 0.468 0.472 0.472 0.463 0.464
ChiAar 0.458 0.48 0.472 0.477 0.489 0.471 0.472
BraCsa 0.181 0.198 0.193 0.197 0.183 0.184 0.184
BraAar 0.462 0.463 0.46 0.462 0.463 0.476 0.475
AthTar 0.483 0.498 0.492 0.496 0.499 0.487 0.488
AthSal 0.452 0.47 0.465 0.469 0.468 0.456 0.456
AthCsa 0.273 0.309 0.301 0.307 0.271 0.283 0.284
AthCru 0.687 0.697 0.694 0.697 0.692 0.683 0.683
AthChi 0.63 0.647 0.641 0.645 0.64 0.63 0.632
AthBra 0.478 0.492 0.487 0.49 0.49 0.478 0.478
AthAar 0.477 0.501 0.492 0.497 0.508 0.486 0.486
meanJI
0.00
0.25
0.50
0.75
1.00

Appendix S15-Table S7: Species pair orthogroup composition comparisons between the orthology inference algorithms and the baseline orthology algorithm (OrthoFinder-BLAST-MCL) using the Jaccard Index. Each row represents a species pair, written in the shorthand found in Table 1, in which the order of the species is the same as the ratio order. BR: Broccoli. OFb: OrthoFinder-BLAST. OFd: OrthoFinder-DIAMOND. OFm: OrthoFinder-MMseqs2. SPd: SonicParanoid-DIAMOND. SPm: SonicParanoid-MMseqs2. ON: OrthNet. (A) Diploid species set. (B) Diploid+higher ploidy species set. Corresponds to Appendix S14.

(A) Diploid set: Species Pairs compared with baseline orthology output

| algorithm | SpeciesPair | numberOfExactOG | totalOG | proportionOfExactOG | meanJI |
|---|---|---|---|---|---|
| OFb | AthChi | 11873 | 20722 | 0.57296593 | 0.656672475 |
| OFb | AthCru | 13159 | 20741 | 0.634443855 | 0.707837192 |
| OFb | AthTar | 8539 | 20718 | 0.412153683 | 0.505029168 |
| OFb | AthAar | 8786 | 20726 | 0.423911995 | 0.506361905 |
| OFb | ChiCru | 11842 | 20435 | 0.579495963 | 0.658346634 |
| OFb | ChiTar | 8172 | 20412 | 0.400352734 | 0.490671527 |
| OFb | ChiAar | 8354 | 20441 | 0.40868842 | 0.491732706 |
| OFb | CruTar | 8519 | 20463 | 0.416312369 | 0.508105335 |
| OFb | CruAar | 8677 | 20479 | 0.423702329 | 0.507571137 |
| OFb | TarAar | 6778 | 17526 | 0.386739701 | 0.481510581 |
| OFd | AthChi | 11836 | 20940 | 0.565234002 | 0.649806102 |
| OFd | AthCru | 13153 | 20965 | 0.627378965 | 0.70083595 |
| OFd | AthTar | 8499 | 20932 | 0.406029046 | 0.499115513 |
| OFd | AthAar | 8736 | 20929 | 0.417411248 | 0.500151144 |
| OFd | ChiCru | 11801 | 20750 | 0.568722892 | 0.647307667 |
| OFd | ChiTar | 8062 | 20726 | 0.388980025 | 0.479974873 |
| OFd | ChiAar | 8289 | 20719 | 0.400067571 | 0.484013826 |
| OFd | CruTar | 8473 | 20714 | 0.409047021 | 0.500994764 |
| OFd | CruAar | 8623 | 20720 | 0.416167954 | 0.500240784 |
| OFd | TarAar | 6706 | 17772 | 0.377335134 | 0.473073738 |
| OFm | AthChi | 11885 | 20837 | 0.570379613 | 0.654045914 |
| OFm | AthCru | 13184 | 20853 | 0.63223517 | 0.705785029 |
| OFm | AthTar | 8532 | 20839 | 0.409424636 | 0.502233025 |
| OFm | AthAar | 8780 | 20835 | 0.421406287 | 0.503795934 |
| OFm | ChiCru | 11837 | 20547 | 0.576093834 | 0.654836988 |
| OFm | ChiTar | 8136 | 20532 | 0.396259497 | 0.486864903 |
| OFm | ChiAar | 8331 | 20545 | 0.405500122 | 0.489019513 |
| OFm | CruTar | 8510 | 20563 | 0.413850119 | 0.505856876 |
| OFm | CruAar | 8658 | 20574 | 0.420822397 | 0.504835662 |
| OFm | TarAar | 6725 | 17675 | 0.380480905 | 0.475968908 |
| ON | AthChi | 9887 | 19223 | 0.51433179 | 0.622483926 |
| ON | AthCru | 11032 | 19226 | 0.573806304 | 0.670473151 |
| ON | AthTar | 7319 | 19223 | 0.38074182 | 0.490232521 |
| ON | AthAar | 7827 | 19220 | 0.40723205 | 0.5021104 |
| ON | ChiCru | 9843 | 18922 | 0.520188141 | 0.621144179 |
| ON | ChiTar | 6801 | 18922 | 0.359422894 | 0.467523674 |
| ON | ChiAar | 7252 | 18922 | 0.383257584 | 0.480851549 |
| ON | CruTar | 7267 | 19068 | 0.381109713 | 0.489924226 |
| ON | CruAar | 7721 | 19068 | 0.404919236 | 0.500992651 |
| ON | TarAar | 5976 | 16784 | 0.356053384 | 0.464872869 |
| SPd | AthChi | 12072 | 21821 | 0.553228541 | 0.636025607 |
| SPd | AthCru | 13477 | 21868 | 0.616288641 | 0.688952433 |
| SPd | AthTar | 8693 | 21706 | 0.400488344 | 0.491627927 |

| | | | | | |
|---|---|---|---|---|---|
| SPd | AthAar | 8730 | 21658 | 0.403084311 | 0.487292642 |
| SPd | ChiCru | 12045 | 21608 | 0.557432432 | 0.63686439 |
| SPd | ChiTar | 8270 | 21495 | 0.384740637 | 0.474090484 |
| SPd | ChiAar | 8322 | 21355 | 0.389697963 | 0.474987179 |
| SPd | CruTar | 8690 | 21390 | 0.40626461 | 0.496896415 |
| SPd | CruAar | 8637 | 21352 | 0.404505433 | 0.490098699 |
| SPd | TarAar | 6723 | 18127 | 0.370883213 | 0.471318135 |
| SPm | AthChi | 12101 | 21857 | 0.553644141 | 0.636383634 |
| SPm | AthCru | 13495 | 21905 | 0.616069391 | 0.688946251 |
| SPm | AthTar | 8694 | 21752 | 0.399687385 | 0.490777729 |
| SPm | AthAar | 8747 | 21695 | 0.403180456 | 0.487423026 |
| SPm | ChiCru | 12054 | 21632 | 0.55723003 | 0.637389982 |
| SPm | ChiTar | 8273 | 21530 | 0.384254529 | 0.473843165 |
| SPm | ChiAar | 8335 | 21390 | 0.389668069 | 0.474879869 |
| SPm | CruTar | 8696 | 21434 | 0.405710553 | 0.496434889 |
| SPm | CruAar | 8658 | 21392 | 0.40473074 | 0.490178792 |
| SPm | TarAar | 6723 | 18150 | 0.370413223 | 0.470898979 |
| BR | AthChi | 11289 | 21147 | 0.533834586 | 0.628279664 |
| BR | AthCru | 12789 | 21171 | 0.604081054 | 0.684808105 |
| BR | AthTar | 8251 | 21088 | 0.391265175 | 0.491339537 |
| BR | AthAar | 8154 | 21070 | 0.386995729 | 0.478838839 |
| BR | ChiCru | 11259 | 20917 | 0.538270306 | 0.626328216 |
| BR | ChiTar | 7703 | 20828 | 0.369838679 | 0.468236812 |
| BR | ChiAar | 7628 | 20794 | 0.366836587 | 0.459931215 |
| BR | CruTar | 8230 | 20862 | 0.394497172 | 0.493553566 |
| BR | CruAar | 8064 | 20841 | 0.38692961 | 0.480076004 |
| BR | TarAar | 6331 | 18108 | 0.349624475 | 0.456369556 |

| sdJI | seJI |
|---|---|
| 0.420200707 | 0.002919046 |
| 0.404676218 | 0.002809913 |
| 0.443238379 | 0.003079381 |
| 0.447868928 | 0.003110951 |
| 0.422252044 | 0.002953823 |
| 0.443534516 | 0.00310445 |
| 0.447074306 | 0.003127005 |
| 0.443643542 | 0.003101341 |
| 0.447076787 | 0.00312412 |
| 0.439498482 | 0.003319831 |
| 0.422718773 | 0.002921213 |
| 0.408596433 | 0.002821936 |
| 0.443684623 | 0.003066684 |
| 0.448099241 | 0.003097419 |
| 0.426791845 | 0.002962832 |
| 0.443408752 | 0.00307997 |
| 0.446791903 | 0.003103994 |
| 0.444220578 | 0.003086503 |
| 0.447396056 | 0.003108116 |
| 0.439101753 | 0.003293798 |
| 0.42133011 | 0.002918804 |
| 0.405824986 | 0.002810312 |
| 0.443594173 | 0.003072893 |
| 0.448107633 | 0.003104457 |
| 0.423665248 | 0.00295562 |
| 0.443543031 | 0.003095424 |
| 0.446866117 | 0.003117628 |
| 0.443855017 | 0.003095265 |
| 0.447112479 | 0.003117148 |
| 0.439105822 | 0.003302855 |
| 0.415731083 | 0.002998485 |
| 0.405856305 | 0.002927034 |
| 0.431579799 | 0.003112795 |
| 0.438611146 | 0.003163756 |
| 0.418913264 | 0.003045374 |
| 0.43004324 | 0.003126285 |
| 0.435981026 | 0.003169451 |
| 0.431994468 | 0.003128424 |
| 0.437989887 | 0.003171842 |
| 0.428458066 | 0.003307201 |
| 0.432099365 | 0.002925138 |
| 0.418390459 | 0.002829289 |
| 0.448132732 | 0.003041703 |

| | |
|---|---|
| 0.450977773 | 0.003064404 |
| 0.435444121 | 0.002962273 |
| 0.449507029 | 0.003065969 |
| 0.451132106 | 0.003087123 |
| 0.448626044 | 0.003067461 |
| 0.450233876 | 0.003081193 |
| 0.443581996 | 0.003294663 |
| 0.432099288 | 0.002922727 |
| 0.418355003 | 0.002826658 |
| 0.44808258 | 0.003038145 |
| 0.451030853 | 0.00306215 |
| 0.435030724 | 0.002957819 |
| 0.449367095 | 0.003062522 |
| 0.451194737 | 0.003085025 |
| 0.448573246 | 0.003063951 |
| 0.450407011 | 0.003079495 |
| 0.443554336 | 0.003292369 |
| 0.426557445 | 0.002933277 |
| 0.414131107 | 0.002846211 |
| 0.441922729 | 0.003043187 |
| 0.444547023 | 0.003062566 |
| 0.430823931 | 0.00297886 |
| 0.441282015 | 0.003057683 |
| 0.442438976 | 0.003068205 |
| 0.442639714 | 0.00306459 |
| 0.443958351 | 0.003075268 |
| 0.43680451 | 0.003246025 |

(B) Diploid + higher ploidy set: Species Pairs compared with baseline orthology output

| algorithm | SpeciesPair | numberOfExactOG | totalOG | proportionOfExactOG | meanJI |
|---|---|---|---|---|---|
| OFb | AthChi | 11778 | 20962 | 0.561873867 | 0.64662516 |
| OFb | AthCru | 13051 | 20971 | 0.622335606 | 0.696991301 |
| OFb | AthTar | 8472 | 20952 | 0.404352806 | 0.498080469 |
| OFb | AthSal | 6372 | 20978 | 0.303746782 | 0.469707706 |
| OFb | AthBra | 6634 | 20967 | 0.316401965 | 0.492358644 |
| OFb | AthCsa | 922 | 20904 | 0.044106391 | 0.308990869 |
| OFb | AthAar | 8783 | 20951 | 0.419216267 | 0.501025658 |
| OFb | ChiCru | 11725 | 20906 | 0.560843777 | 0.640516698 |
| OFb | ChiTar | 8062 | 20883 | 0.386055643 | 0.477603453 |
| OFb | ChiSal | 6089 | 20914 | 0.291144688 | 0.453204613 |
| OFb | ChiBra | 6288 | 20906 | 0.300774897 | 0.474239281 |
| OFb | ChiCsa | 934 | 20826 | 0.044847786 | 0.300855541 |
| OFb | ChiAar | 8312 | 20893 | 0.397836596 | 0.480362425 |
| OFb | CruTar | 8451 | 20892 | 0.404508903 | 0.496999603 |
| OFb | CruSal | 6377 | 20914 | 0.304915368 | 0.470762673 |
| OFb | CruBra | 6726 | 20904 | 0.321756602 | 0.499060143 |
| OFb | CruCsa | 1055 | 20874 | 0.050541343 | 0.331635883 |
| OFb | CruAar | 8655 | 20900 | 0.414114833 | 0.497229942 |
| OFb | TarSal | 5231 | 18027 | 0.290175847 | 0.467446528 |
| OFb | TarBra | 5101 | 18004 | 0.283325928 | 0.463100786 |
| OFb | TarCsa | 662 | 17908 | 0.036966719 | 0.268801037 |
| OFb | TarAar | 6718 | 17992 | 0.373388173 | 0.467813935 |
| OFb | SalBra | 7005 | 20883 | 0.33544031 | 0.457094291 |
| OFb | SalCsa | 420 | 20763 | 0.020228291 | 0.18943443 |
| OFb | SalAar | 5156 | 18201 | 0.283281138 | 0.459895087 |
| OFb | BraCsa | 507 | 21124 | 0.024001136 | 0.198443008 |
| OFb | BraAar | 5183 | 18184 | 0.285030796 | 0.46328175 |
| OFb | CsaAar | 563 | 18141 | 0.031034673 | 0.259711735 |
| OFd | AthChi | 11909 | 21392 | 0.556703441 | 0.640741692 |
| OFd | AthCru | 13314 | 21412 | 0.621800859 | 0.693518464 |
| OFd | AthTar | 8524 | 21382 | 0.398653073 | 0.491540503 |
| OFd | AthSal | 6415 | 21404 | 0.299710335 | 0.464577437 |
| OFd | AthBra | 6704 | 21401 | 0.31325639 | 0.486597282 |
| OFd | AthCsa | 840 | 21370 | 0.03930744 | 0.300726363 |
| OFd | AthAar | 8746 | 21380 | 0.409073901 | 0.491711855 |
| OFd | ChiCru | 11898 | 21361 | 0.556996395 | 0.634573163 |
| OFd | ChiTar | 8091 | 21335 | 0.379235997 | 0.469799626 |
| OFd | ChiSal | 6095 | 21358 | 0.285373162 | 0.445945561 |
| OFd | ChiBra | 6355 | 21364 | 0.297463022 | 0.468272456 |
| OFd | ChiCsa | 902 | 21317 | 0.042313646 | 0.294131698 |
| OFd | ChiAar | 8277 | 21339 | 0.387881344 | 0.47168766 |
| OFd | CruTar | 8535 | 21337 | 0.400009373 | 0.491230326 |
| OFd | CruSal | 6405 | 21354 | 0.299943804 | 0.46464993 |

| | | | | | |
|---|---|---|---|---|---|
| OFd | CruBra | 6789 | 21355 | 0.317911496 | 0.49297433 |
| OFd | CruCsa | 980 | 21332 | 0.045940371 | 0.324737223 |
| OFd | CruAar | 8632 | 21344 | 0.404422789 | 0.488125835 |
| OFd | TarSal | 5287 | 18347 | 0.288167003 | 0.462914699 |
| OFd | TarBra | 5112 | 18335 | 0.278811017 | 0.456933953 |
| OFd | TarCsa | 614 | 18297 | 0.033557414 | 0.260833144 |
| OFd | TarAar | 6671 | 18313 | 0.364276743 | 0.460460816 |
| OFd | SalBra | 7208 | 21638 | 0.333117663 | 0.451321117 |
| OFd | SalCsa | 351 | 21484 | 0.01633774 | 0.183023687 |
| OFd | SalAar | 5170 | 18268 | 0.28300854 | 0.456919467 |
| OFd | BraCsa | 445 | 21857 | 0.02035961 | 0.192763777 |
| OFd | BraAar | 5200 | 18255 | 0.284853465 | 0.45968109 |
| OFd | CsaAar | 509 | 18245 | 0.027898054 | 0.253731085 |
| OFm | AthChi | 11881 | 21174 | 0.561112685 | 0.645069014 |
| OFm | AthCru | 13232 | 21187 | 0.624533912 | 0.69721403 |
| OFm | AthTar | 8531 | 21166 | 0.403052065 | 0.495955855 |
| OFm | AthSal | 6397 | 21190 | 0.301887683 | 0.468521529 |
| OFm | AthBra | 6662 | 21188 | 0.314423258 | 0.49025571 |
| OFm | AthCsa | 911 | 21139 | 0.0430957 | 0.306857771 |
| OFm | AthAar | 8782 | 21166 | 0.414910706 | 0.49725725 |
| OFm | ChiCru | 11828 | 21109 | 0.560329717 | 0.638734589 |
| OFm | ChiTar | 8093 | 21094 | 0.383663601 | 0.47440422 |
| OFm | ChiSal | 6081 | 21118 | 0.287953405 | 0.450783471 |
| OFm | ChiBra | 6290 | 21117 | 0.29786428 | 0.471734587 |
| OFm | ChiCsa | 939 | 21063 | 0.044580544 | 0.298742251 |
| OFm | ChiAar | 8312 | 21096 | 0.394008343 | 0.477045092 |
| OFm | CruTar | 8515 | 21075 | 0.404033215 | 0.495584998 |
| OFm | CruSal | 6377 | 21092 | 0.30234212 | 0.469098483 |
| OFm | CruBra | 6736 | 21087 | 0.319438517 | 0.497207179 |
| OFm | CruCsa | 1039 | 21068 | 0.049316499 | 0.329682048 |
| OFm | CruAar | 8644 | 21086 | 0.409940245 | 0.493785809 |
| OFm | TarSal | 5200 | 18196 | 0.285777094 | 0.464293158 |
| OFm | TarBra | 5068 | 18184 | 0.278706555 | 0.459389913 |
| OFm | TarCsa | 664 | 18119 | 0.036646614 | 0.265348439 |
| OFm | TarAar | 6703 | 18168 | 0.368945399 | 0.464196291 |
| OFm | SalBra | 7060 | 21023 | 0.33582267 | 0.456728758 |
| OFm | SalCsa | 403 | 20919 | 0.019264783 | 0.188219774 |
| OFm | SalAar | 5154 | 18228 | 0.28275181 | 0.459069553 |
| OFm | BraCsa | 486 | 21254 | 0.022866284 | 0.196686541 |
| OFm | BraAar | 5168 | 18213 | 0.283753363 | 0.46167231 |
| OFm | CsaAar | 565 | 18199 | 0.031045662 | 0.257551851 |
| ON | AthChi | 10606 | 19640 | 0.540020367 | 0.640055302 |
| ON | AthCru | 11924 | 19640 | 0.60712831 | 0.692321218 |
| ON | AthTar | 7751 | 19640 | 0.394653768 | 0.499156128 |
| ON | AthSal | 5700 | 19640 | 0.290224033 | 0.468319528 |

| | | | | | |
|---|---|---|---|---|---|
| ON | AthBra | 5970 | 19639 | 0.303986965 | 0.490159503 |
| ON | AthCsa | 206 | 19644 | 0.010486663 | 0.271451213 |
| ON | AthAar | 8181 | 19638 | 0.416590284 | 0.508435285 |
| ON | ChiCru | 10627 | 19290 | 0.550907206 | 0.643105024 |
| ON | ChiTar | 7220 | 19290 | 0.374287195 | 0.478324449 |
| ON | ChiSal | 5225 | 19290 | 0.270865734 | 0.449643945 |
| ON | ChiBra | 5492 | 19291 | 0.284692344 | 0.47247787 |
| ON | ChiCsa | 169 | 19293 | 0.008759654 | 0.261778907 |
| ON | ChiAar | 7585 | 19290 | 0.393208917 | 0.488513745 |
| ON | CruTar | 7726 | 19524 | 0.395718091 | 0.49926027 |
| ON | CruSal | 5645 | 19528 | 0.289072102 | 0.468985586 |
| ON | CruBra | 5990 | 19527 | 0.30675475 | 0.495583026 |
| ON | CruCsa | 230 | 19528 | 0.01177796 | 0.288136513 |
| ON | CruAar | 8084 | 19525 | 0.414033291 | 0.506845208 |
| ON | TarSal | 4384 | 17080 | 0.256674473 | 0.451213138 |
| ON | TarBra | 4414 | 17081 | 0.258415784 | 0.450144955 |
| ON | TarCsa | 108 | 17079 | 0.006323555 | 0.231319101 |
| ON | TarAar | 6161 | 17077 | 0.360777654 | 0.466507285 |
| ON | SalBra | 5985 | 18914 | 0.316432272 | 0.444752584 |
| ON | SalCsa | 116 | 18910 | 0.00613432 | 0.175612447 |
| ON | SalAar | 4645 | 17064 | 0.272210502 | 0.46040289 |
| ON | BraCsa | 125 | 19312 | 0.006472659 | 0.182595459 |
| ON | BraAar | 4708 | 17065 | 0.275886317 | 0.462929995 |
| ON | CsaAar | 101 | 17062 | 0.005919587 | 0.233188251 |
| SPd | AthChi | 11823 | 21687 | 0.545165306 | 0.62991578 |
| SPd | AthCru | 13177 | 21704 | 0.607123111 | 0.682639163 |
| SPd | AthTar | 8526 | 21602 | 0.394685677 | 0.487012615 |
| SPd | AthSal | 6337 | 21650 | 0.292702079 | 0.455518124 |
| SPd | AthBra | 6705 | 21657 | 0.309599668 | 0.478084645 |
| SPd | AthCsa | 382 | 21690 | 0.017611803 | 0.283404379 |
| SPd | AthAar | 8640 | 21556 | 0.400816478 | 0.48564328 |
| SPd | ChiCru | 11778 | 21501 | 0.547788475 | 0.629760602 |
| SPd | ChiTar | 8127 | 21410 | 0.379588977 | 0.469945563 |
| SPd | ChiSal | 6058 | 21482 | 0.282003538 | 0.441126399 |
| SPd | ChiBra | 6380 | 21528 | 0.296358231 | 0.463364713 |
| SPd | ChiCsa | 376 | 21676 | 0.017346374 | 0.272697261 |
| SPd | ChiAar | 8191 | 21288 | 0.384770763 | 0.471396637 |
| SPd | CruTar | 8520 | 21579 | 0.394828305 | 0.485718297 |
| SPd | CruSal | 6349 | 21631 | 0.293513938 | 0.45501834 |
| SPd | CruBra | 6801 | 21641 | 0.31426459 | 0.484293706 |
| SPd | CruCsa | 473 | 21727 | 0.021770148 | 0.303739814 |
| SPd | CruAar | 8528 | 21561 | 0.395528964 | 0.481232123 |
| SPd | TarSal | 5360 | 18386 | 0.291526161 | 0.46627987 |
| SPd | TarBra | 5300 | 18369 | 0.288529588 | 0.463322612 |
| SPd | TarCsa | 228 | 18357 | 0.01242033 | 0.24751153 |

| | | | | | |
|---|---|---|---|---|---|
| SPd | TarAar | 6616 | 18212 | 0.36327696 | 0.465360906 |
| SPd | SalBra | 7403 | 21758 | 0.340242669 | 0.456893115 |
| SPd | SalCsa | 211 | 21473 | 0.009826293 | 0.180366708 |
| SPd | SalAar | 5153 | 17377 | 0.296541405 | 0.471879244 |
| SPd | BraCsa | 245 | 22513 | 0.010882601 | 0.183711496 |
| SPd | BraAar | 5267 | 17372 | 0.30318904 | 0.475879127 |
| SPd | CsaAar | 201 | 17360 | 0.011578341 | 0.250189398 |
| SPm | AthChi | 11863 | 21682 | 0.547135873 | 0.631707839 |
| SPm | AthCru | 13198 | 21705 | 0.608062658 | 0.683386501 |
| SPm | AthTar | 8549 | 21603 | 0.395732074 | 0.487570775 |
| SPm | AthSal | 6354 | 21652 | 0.293460188 | 0.456156989 |
| SPm | AthBra | 6713 | 21660 | 0.309926131 | 0.478447379 |
| SPm | AthCsa | 381 | 21685 | 0.017569749 | 0.284121314 |
| SPm | AthAar | 8651 | 21554 | 0.401364016 | 0.486317587 |
| SPm | ChiCru | 11803 | 21481 | 0.549462316 | 0.631756421 |
| SPm | ChiTar | 8131 | 21397 | 0.380006543 | 0.47059776 |
| SPm | ChiSal | 6082 | 21466 | 0.28333178 | 0.442342916 |
| SPm | ChiBra | 6389 | 21506 | 0.297079885 | 0.464160535 |
| SPm | ChiCsa | 375 | 21664 | 0.017309823 | 0.273266538 |
| SPm | ChiAar | 8206 | 21277 | 0.385674672 | 0.47227283 |
| SPm | CruTar | 8541 | 21563 | 0.396095163 | 0.48682949 |
| SPm | CruSal | 6355 | 21612 | 0.294049602 | 0.455925256 |
| SPm | CruBra | 6817 | 21612 | 0.315426615 | 0.486077688 |
| SPm | CruCsa | 464 | 21715 | 0.021367718 | 0.304424631 |
| SPm | CruAar | 8539 | 21546 | 0.396314861 | 0.482079903 |
| SPm | TarSal | 5380 | 18413 | 0.292184869 | 0.46671814 |
| SPm | TarBra | 5308 | 18385 | 0.288713625 | 0.463449072 |
| SPm | TarCsa | 229 | 18387 | 0.012454452 | 0.247417887 |
| SPm | TarAar | 6610 | 18242 | 0.362350619 | 0.464064052 |
| SPm | SalBra | 7435 | 21747 | 0.341886237 | 0.458478842 |
| SPm | SalCsa | 215 | 21471 | 0.010013507 | 0.180491628 |
| SPm | SalAar | 5166 | 17401 | 0.29687949 | 0.47197427 |
| SPm | BraCsa | 246 | 22515 | 0.010926049 | 0.183748209 |
| SPm | BraAar | 5264 | 17389 | 0.30272011 | 0.475495707 |
| SPm | CsaAar | 201 | 17383 | 0.011563021 | 0.250182583 |
| BR | AthChi | 11741 | 21771 | 0.539295393 | 0.62968356 |
| BR | AthCru | 13294 | 21783 | 0.61029243 | 0.687052115 |
| BR | AthTar | 8379 | 21735 | 0.385507246 | 0.483268193 |
| BR | AthSal | 6135 | 21751 | 0.282055997 | 0.452223508 |
| BR | AthBra | 6551 | 21756 | 0.301112337 | 0.477754222 |
| BR | AthCsa | 241 | 21758 | 0.011076386 | 0.273120173 |
| BR | AthAar | 8413 | 21728 | 0.387196244 | 0.476839141 |
| BR | ChiCru | 11709 | 21517 | 0.544174374 | 0.628790862 |
| BR | ChiTar | 7822 | 21447 | 0.364713013 | 0.461465583 |
| BR | ChiSal | 5711 | 21498 | 0.265652619 | 0.433568739 |

| | | | | | |
|---|---|---|---|---|---|
| BR | ChiBra | 6116 | 21512 | 0.284306434 | 0.458991995 |
| BR | ChiCsa | 190 | 21558 | 0.008813434 | 0.260167886 |
| BR | ChiAar | 7868 | 21426 | 0.367217399 | 0.458405687 |
| BR | CruTar | 8346 | 21679 | 0.384980857 | 0.482147524 |
| BR | CruSal | 6093 | 21689 | 0.280925815 | 0.452041321 |
| BR | CruBra | 6632 | 21701 | 0.305608036 | 0.483790205 |
| BR | CruCsa | 278 | 21713 | 0.01280339 | 0.291581278 |
| BR | CruAar | 8313 | 21675 | 0.383529412 | 0.474007835 |
| BR | TarSal | 4921 | 18535 | 0.265497707 | 0.448822911 |
| BR | TarBra | 4977 | 18514 | 0.268823593 | 0.449272533 |
| BR | TarCsa | 112 | 18518 | 0.006048169 | 0.23182341 |
| BR | TarAar | 6369 | 18432 | 0.345540365 | 0.449036075 |
| BR | SalBra | 7217 | 21020 | 0.343339676 | 0.460767588 |
| BR | SalCsa | 131 | 20877 | 0.006274848 | 0.175648528 |
| BR | SalAar | 4900 | 17490 | 0.280160091 | 0.457157672 |
| BR | BraAar | 5042 | 17494 | 0.288213102 | 0.462326906 |
| BR | CsaAar | 103 | 17505 | 0.005884033 | 0.233240474 |
| BR | BraCsa | 154 | 22015 | 0.006995231 | 0.180599117 |

| sdJI | seJI |
|---|---|
| 0.423789489 | 0.002927075 |
| 0.409877358 | 0.002830377 |
| 0.443486492 | 0.003063851 |
| 0.394575323 | 0.002724256 |
| 0.38875323 | 0.002684762 |
| 0.229525989 | 0.001587513 |
| 0.448579438 | 0.00309911 |
| 0.428668701 | 0.002964738 |
| 0.44331817 | 0.003067744 |
| 0.396419412 | 0.002741172 |
| 0.389894156 | 0.002696567 |
| 0.235163003 | 0.001629544 |
| 0.447378173 | 0.003095098 |
| 0.444203876 | 0.003073211 |
| 0.394849264 | 0.002730315 |
| 0.388284978 | 0.002685566 |
| 0.236650428 | 0.001637964 |
| 0.447916735 | 0.003098305 |
| 0.396048551 | 0.00294976 |
| 0.389777277 | 0.002904906 |
| 0.229629409 | 0.001715947 |
| 0.439554354 | 0.003276973 |
| 0.410957202 | 0.002843807 |
| 0.189282487 | 0.001313607 |
| 0.388541447 | 0.002879982 |
| 0.197167932 | 0.001356589 |
| 0.386311742 | 0.002864793 |
| 0.217494991 | 0.0016148 |
| 0.426802482 | 0.002918107 |
| 0.41371295 | 0.002827291 |
| 0.444355192 | 0.003038828 |
| 0.395736223 | 0.002704944 |
| 0.390970712 | 0.002672558 |
| 0.226449254 | 0.001549061 |
| 0.448758099 | 0.003069082 |
| 0.432220791 | 0.002957296 |
| 0.443813698 | 0.003038466 |
| 0.397112881 | 0.002717276 |
| 0.391700893 | 0.002679867 |
| 0.235409741 | 0.001612358 |
| 0.447013694 | 0.003060087 |
| 0.445465058 | 0.003049629 |
| 0.395997367 | 0.002709896 |

| | |
|---|---|
| 0.390422465 | 0.002671684 |
| 0.234592394 | 0.001606195 |
| 0.448200783 | 0.003067854 |
| 0.397326225 | 0.002933356 |
| 0.39068411 | 0.002885262 |
| 0.227775791 | 0.001683905 |
| 0.438534246 | 0.003240588 |
| 0.413876525 | 0.002813599 |
| 0.185423865 | 0.001265051 |
| 0.390167125 | 0.002886723 |
| 0.193757089 | 0.001310576 |
| 0.388313255 | 0.00287403 |
| 0.215192907 | 0.001593148 |
| 0.424972121 | 0.002920512 |
| 0.41112719 | 0.002824499 |
| 0.444191971 | 0.003053172 |
| 0.394544179 | 0.00271038 |
| 0.388960598 | 0.002672149 |
| 0.228412284 | 0.001571004 |
| 0.448749347 | 0.003084498 |
| 0.430123228 | 0.002960459 |
| 0.443716393 | 0.003055104 |
| 0.396024534 | 0.002725183 |
| 0.389580906 | 0.002680906 |
| 0.235359009 | 0.001621701 |
| 0.447371509 | 0.003080124 |
| 0.444978779 | 0.003065177 |
| 0.394539899 | 0.00271664 |
| 0.388123316 | 0.002672775 |
| 0.235318445 | 0.001621229 |
| 0.447888771 | 0.003084417 |
| 0.395401194 | 0.002931231 |
| 0.389072702 | 0.002885267 |
| 0.229578784 | 0.001705551 |
| 0.439226989 | 0.003258633 |
| 0.41096307 | 0.002834363 |
| 0.187845517 | 0.001298764 |
| 0.388647973 | 0.002878637 |
| 0.194996005 | 0.001337536 |
| 0.386048347 | 0.002860559 |
| 0.217753283 | 0.001614139 |
| 0.416070361 | 0.002968903 |
| 0.403904408 | 0.002882092 |
| 0.435798086 | 0.003109672 |
| 0.384412538 | 0.002743006 |

| | |
|---|---|
| 0.380135279 | 0.002712555 |
| 0.179175042 | 0.001278388 |
| 0.441642813 | 0.003151538 |
| 0.417918366 | 0.003009022 |
| 0.434296042 | 0.003126942 |
| 0.382910812 | 0.002756967 |
| 0.378019722 | 0.00272168 |
| 0.180186486 | 0.001297246 |
| 0.438792743 | 0.003159318 |
| 0.436443926 | 0.003123518 |
| 0.383955741 | 0.002747592 |
| 0.378954727 | 0.002711874 |
| 0.183653365 | 0.001314226 |
| 0.441174491 | 0.003157293 |
| 0.381394749 | 0.002918306 |
| 0.378319365 | 0.002894689 |
| 0.177530698 | 0.001358446 |
| 0.431175624 | 0.003299502 |
| 0.40083113 | 0.002914538 |
| 0.154545435 | 0.001123855 |
| 0.378699082 | 0.002899037 |
| 0.157003667 | 0.001129786 |
| 0.377734563 | 0.002891569 |
| 0.169537464 | 0.001297928 |
| 0.432458662 | 0.0029366 |
| 0.418889126 | 0.002843343 |
| 0.446912379 | 0.003040713 |
| 0.397969718 | 0.002704712 |
| 0.395530018 | 0.002687697 |
| 0.203018756 | 0.001378499 |
| 0.450081352 | 0.00306554 |
| 0.435658003 | 0.002971094 |
| 0.448161309 | 0.003062852 |
| 0.400582574 | 0.002733095 |
| 0.397167429 | 0.002706897 |
| 0.209548602 | 0.001423296 |
| 0.449685247 | 0.003082061 |
| 0.447696221 | 0.003047669 |
| 0.398385373 | 0.002708726 |
| 0.394995732 | 0.002685058 |
| 0.210987018 | 0.001431383 |
| 0.449398749 | 0.003060535 |
| 0.402290936 | 0.002966857 |
| 0.397896036 | 0.002935803 |
| 0.202247475 | 0.001492733 |

| | |
|---|---|
| 0.441637622 | 0.003272557 |
| 0.418311506 | 0.002835896 |
| 0.174615122 | 0.001191614 |
| 0.393512516 | 0.002985184 |
| 0.180224042 | 0.001201147 |
| 0.393380725 | 0.002984614 |
| 0.192494678 | 0.001460979 |
| 0.432043139 | 0.002934117 |
| 0.418833671 | 0.002842901 |
| 0.447182416 | 0.00304248 |
| 0.398019789 | 0.002704928 |
| 0.395489932 | 0.002687238 |
| 0.202747328 | 0.001376814 |
| 0.450107481 | 0.00306586 |
| 0.435017452 | 0.002968106 |
| 0.448109877 | 0.003063431 |
| 0.400777895 | 0.002735446 |
| 0.397037967 | 0.002707399 |
| 0.209542939 | 0.001423652 |
| 0.4497897 | 0.003083574 |
| 0.447773318 | 0.003049324 |
| 0.398233527 | 0.002708884 |
| 0.394650989 | 0.002684514 |
| 0.210412996 | 0.001427883 |
| 0.449488642 | 0.003062213 |
| 0.40247449 | 0.002966034 |
| 0.397928831 | 0.002934767 |
| 0.202488226 | 0.001493291 |
| 0.441569545 | 0.003269361 |
| 0.418427226 | 0.002837398 |
| 0.174794989 | 0.001192897 |
| 0.393821335 | 0.002985466 |
| 0.180321015 | 0.00120174 |
| 0.393354813 | 0.002982958 |
| 0.192672754 | 0.001461362 |
| 0.428812887 | 0.002906221 |
| 0.415785553 | 0.002817154 |
| 0.443360286 | 0.003007301 |
| 0.391672632 | 0.002655728 |
| 0.389971964 | 0.002643893 |
| 0.184427677 | 0.001250307 |
| 0.446145123 | 0.003026678 |
| 0.432249488 | 0.002946752 |
| 0.442069998 | 0.003018615 |
| 0.391774939 | 0.002672007 |

| | |
|---|---|
| 0.389928826 | 0.002658551 |
| 0.186923437 | 0.001273091 |
| 0.443744395 | 0.003031533 |
| 0.443788153 | 0.003014089 |
| 0.391185258 | 0.002656212 |
| 0.389141358 | 0.002641603 |
| 0.189164225 | 0.001283746 |
| 0.445489055 | 0.00302592 |
| 0.393355863 | 0.002889278 |
| 0.390919708 | 0.002873012 |
| 0.183397798 | 0.001347712 |
| 0.436617113 | 0.003215989 |
| 0.414698822 | 0.002860332 |
| 0.159720742 | 0.00110542 |
| 0.389788537 | 0.002947367 |
| 0.39077828 | 0.002954513 |
| 0.17636345 | 0.001332992 |
| 0.163419519 | 0.001101399 |

Appendix S16-Fig. S9 RAxML tree of all genes from the six Arabidopsis YABBY orthogroups, after 1000 bootstraps. The YABBY sequence from a *Picea glauca* YABBY (BT115385) was used as the outgroup.

A

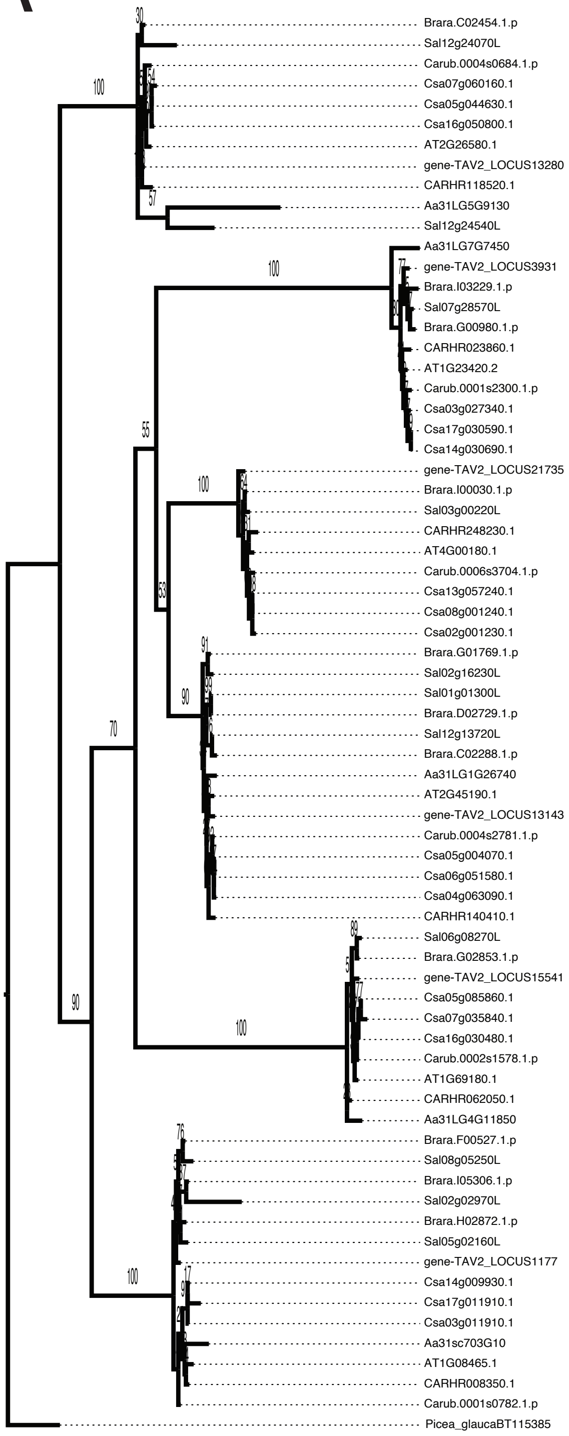

Appendix S17-Fig. S10: Screenshots of YABBY sequence alignments reveal sequence features that could affect whether an orthology inference algorithm incorporates the sequence into an orthogroup. (A) AT2G45190.1 (FIL/YAB1). (B) AT1G08465.1 (YAB2). (C) AT4G00180.1 (YAB3). (D) AT2G26580.1 (YAB5). (E) AT1G69180.1 (CRC). (F) AT1G23420.2 (INO).

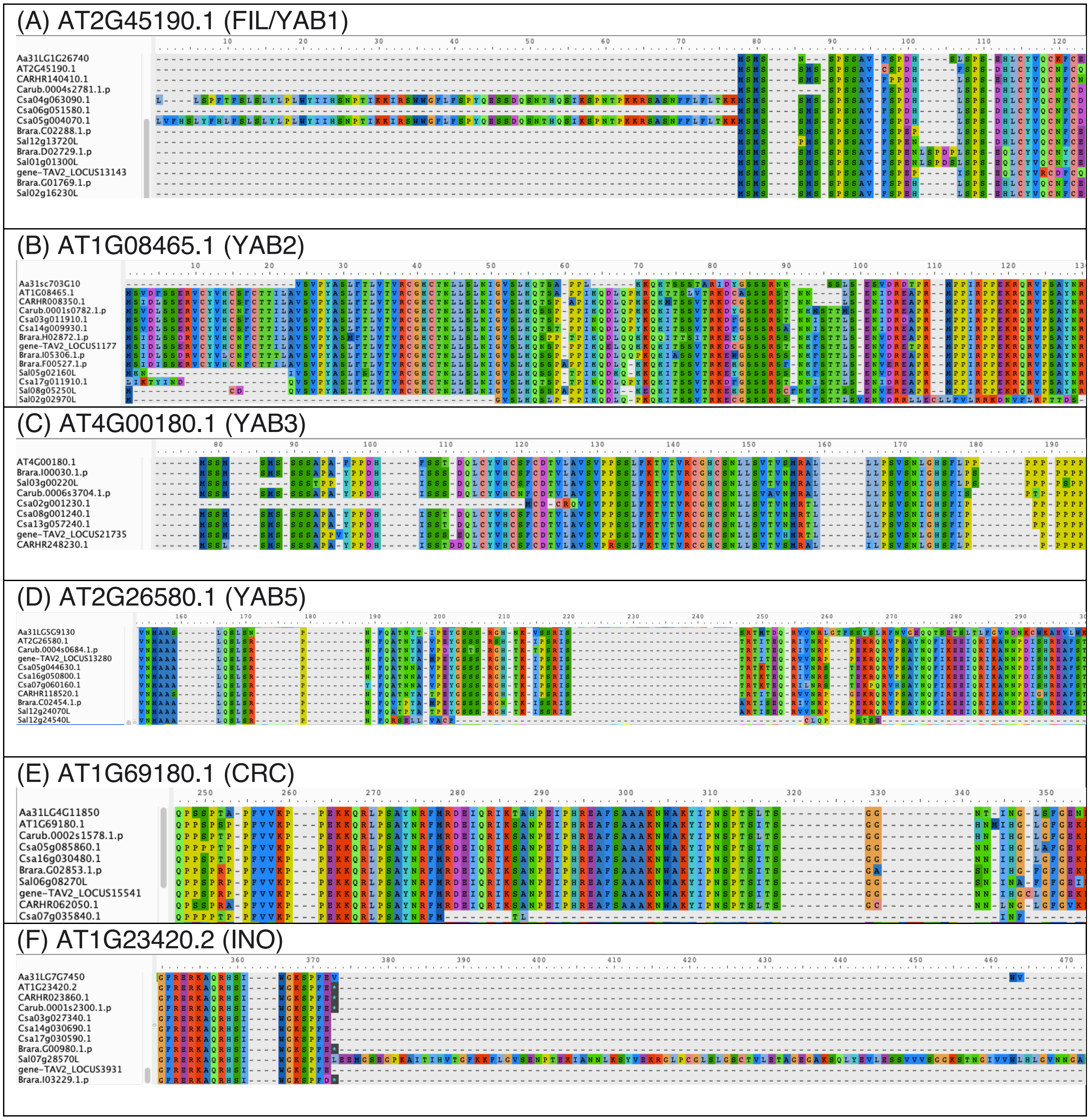

Appendix S18-Table S8: Orthogroup composition outputs of select YABBYs from the diploid set. White indicates the gene is found in the same orthogroup, and black indicates that the gene was not included in any orthogroups. (A) FIL/YAB1. Notice the extra *Aethionema* gene that was only included in OFb. (B) YAB3. Notice that there no *Aethionema* gene included in this orthogroup. BR: Broccoli. OFb: OrthoFinder-BLAST. OFd: OrthoFinder-DIAMOND. OFm: OrthoFinder-MMseqs2. SPd: SonicParanoid-DIAMOND. SPm: SonicParanoid-MMseqs2. ON: OrthNet.

(A) FIL/YAB1

| **gene** | **BR** | **OFb** | **OFd** | **OFm** | **SPd** | **SPm** |
|---|---|---|---|---|---|---|
| *C. rubella* - Carub.0004s2781.1.p | | | | | | |
| ***A. thaliana* - AT2G45190.1 (FIL/YAB1)** | | | | | | |
| *A. arabicum* - Aa31LG1G26740 | | | | | | |
| *C. hirsuta* - CARHR140410.1 | | | | | | |
| *T. arvense* - gene-TAV2_LOCUS13143 | | | | | | |
| *A. arabicum* - Aa31LG2G250 | ■ | | ■ | ■ | ■ | ■ |

(B) YAB3

| **gene** | **BR** | **OFb** | **OFd** | **OFm** | **SPd** | **SPm** |
|---|---|---|---|---|---|---|
| *C. rubella* - Carub.0006s3704.1.p | | | | | | |
| ***A. thaliana* - AT4G00180.1 (YAB3)** | | | | | | |
| *C. hirsuta* -CARHR248230.1 | | | | | | |
| *T. arvense* - gene-TAV2_LOCUS21735 | | | | | | |
